# Homoeologous non-reciprocal translocation explains a major QTL for seed lignin content in oilseed rape (*Brassica napus* L.)

**DOI:** 10.1101/2023.03.10.531521

**Authors:** Hanna Marie Schilbert, Karin Holzenkamp, Prisca Viehöver, Daniela Holtgräwe, Christian Möllers

**Affiliations:** Genetics and Genomics of Plants, CeBiTec and Faculty of Biology, Bielefeld University, Bielefeld, Germany; Graduate School DILS, Bielefeld Institute for Bioinformatics Infrastructure (BIBI), Faculty of Technology, Bielefeld University, Germany; Department of Crop Sciences, Division of Crop Plant Genetics, Georg-August-University, Göttingen, Germany

**Author notes:** Corresponding author: Hanna Marie Schilbert. These authors contributed equally to this work (Hanna Marie Schilbert, Karin Holzenkamp).

**Keywords:** lignin, HNRT, *PAL4*, phenylpropanoid, QTL, seed coat

## Abstract

Oilseed rape is a major oil crop and a valuable protein source for animal and human nutrition. Lignin is a non-digestible, major component of the seed coat with negative effect on sensory quality, bioavailability and usage of oilseed rape’s protein. Hence, seed lignin reduction is of economic and nutritional importance. In this study, the major QTL for reduced lignin content found on chromosome C05 in the DH population SGDH14 x Express 617 was further examined. SGDH14 had lower seed lignin content than Express 617. Harvested seeds from a F2 population of the same cross were additionally field tested and used for seed quality analysis. The F2 population showed a bimodal distribution for seed lignin content. F2 plants with low lignin content had thinner seed coats compared to high lignin lines. Both groups showed a dark seed colour with a slightly lighter colour in the low lignin group indicating that a low lignin content is not necessarily associated with yellow seed colour. Mapping of genomic long-reads from SGDH14 against the Express 617 genome assembly revealed a homoeologous non-reciprocal translocation (HNRT) in the confidence interval of the major QTL for lignin content. A homologous A05 region is duplicated and replaced the C05 region in SGDH14. As consequence several genes located in the C05 region were lost in SGDH14. Thus, a HNRT was identified in the major QTL region for reduced lignin content in the low lignin line SGDH14. The most promising candidate gene related to lignin biosynthesis on C05, *PAL4*, was deleted.

**Key message:** A homoeologous non-reciprocal translocation was identified in the major QTL for seed lignin content in the low lignin line SGDH14. The lignin biosynthetic gene *PAL4* was deleted.

## Introduction

Oilseed rape (*Brassica napus* L.) is one of the most important oil crops worldwide. The allopolyploid species *B. napus* (AACC, 2n=38) was formed by a spontaneous hybridisation and genome doubling between the two diploid species *Brassica rapa* (AA, 2n=20) and *Brassica oleracea* (CC, 2n=18) more than 7,500 years ago (Chalhoub et al. 2014, Lu et al. 2019, Nagaharu 1935). Recombination is one of the major sources of genetic variation and for novel allelic combinations. In many allopolyploid species homoeologous exchanges (HEs) can occur. Thus, mispairing between ancestral chromosomes can lead to deletions, duplications and translocations (Sharpe et al. 1995, Mason and Wendel 2020). Meiotic chromosome pairing that occurs between homoeologous chromosomes with a high sequence similarity leads to enhanced HEs and gene conversion events both initiated by the repair of double-strand breaks (DSB) throughout the *B. napus* genome. These HEs range in size from large segments to single SNPs. HEs are highly enriched in transcriptional regions generating novel gene combinations and can potentially cause phenotypic alterations (Chalhoub et al. 2014, Gaeta and Pires 2010, Stein et al. 2017, Zhang et al. 2020). HEs can result in reciprocal exchange between homoeologous chromosomes (crossover) and non-reciprocal exchange (non-crossover). Non-reciprocal exchanges known as homoeologous non-reciprocal translocations (HNRTs) are duplication/deletion events where an additional copy of a DNA sequence has replaced its homoeologous copy leading to gain and loss of genetic material (reviewed in Gaeta and Pires (2010) and Mason and Wendel (2020)).

Synthetic polyploids and recent allopolyploids contain frequent HEs. For established *B. napus* (Chalhoub et al. 2014) and resynthesised *B. napus* (Szadkowski et al. 2010, Gaeta and Pires 2010) HEs are documented and indicate a diversification in the polyploid genome. In the *B. napus* genome, genomic segments and genes can be found in multiple copies. Two copies of the A and C subgenomes which share the most recent ancestry are primary homologues, while the remaining ones are secondary homologues and paralogous to one or the other copy (Nicolas et al. 2007, Mason and Wendel 2020). It was demonstrated that regions of primary homology show a high recombination rate of chromosomal rearrangements. This represents an up to 100-fold increase compared to the frequency of homologous recombination measured in euploid lines (Nicolas et al. 2007). Further, for *B. napus* a subgenome bias towards replacing larger C-subgenome segments by smaller, homoeologous A-subgenome segments was observed (Samans et al. 2017, Sochorová et al. 2017, Higgins et al. 2018). The reciprocal allele gain and loss observed between the A and C subgenomes ranged from small segments to entire chromosomes, with low quantity for aneuploidy across all gametes in *B. napus* (Higgins et al. 2018). HNRTs and HEs have been conclusively linked to phenotypic changes in *B. napus* like seed yield, seed quality, seed colour, seed fibre content, flowering time, disease resistance and fertility (Hurgobin et al. 2018, Schiessl et al. 2019, Stein et al. 2017). Having both adaptive and agronomic importance, HNRTs and HEs could play an important role in the breeding of improved allopolyploid crops.

Due to a growing demand of plant-based protein for food and feed (Ismail et al. 2020), the breeding aim in *B. napus* is increasing the seed oil and protein content (Si et al. 2003). However, in comparison to soybean meal, which is the most important vegetable protein source, oilseed rape meal has a lower protein content as well as higher anti-nutritive fibre compounds (Nesi et al. 2008). Therefore, genetic reduction of anti-nutritive and non-digestible compounds, such as lignin, is required to develop genotypes with improved nutritional quality for the production of optimised vegetable protein products (Wittkop et al. 2009). Lignin is mainly found in the seed coat. Hence, a low lignin content is associated with less seed coat phenolic compounds and a lighter seed colour (Carré et al. 2016, Wittkop et al. 2012). Attempts made to develop competitive agronomic yellow seeded winter oilseed rape cultivars with low lignin contents under various environmental conditions were unsuccessful to date (Rahman and McVetty 2011, Simbaya et al. 1995).

In diverse populations a large number of quantitative trait loci (QTL) for reduced lignin content on A02, A05, A07, A09, C02, C03, C05 and C07 (Behnke et al. 2018, Wang et al. 2015, Körber et al. 2016) and yellow seed colour on A09 and C08 (Badani et al. 2006, Wang et al. 2017) or both traits (Stein et al. 2013, Liu et al. 2013) were detected on different linkage groups. A major QTL for low lignin content was found on C05 explaining 81 % of the phenotypic variation in the doubled haploid (DH) population derived from the cross of genotype SGDH14 with inbred line 617 of the cultivar Express (Express 617) (Behnke et al. 2018). This QTL co-located with a QTL for oil and protein content with opposite additive effects. The DH population, showed a large bimodal distribution for the Acid Detergent Lignin (ADL, synonym: lignin) content ranging from 6.5 – 14.5 %. SGDH14 was with 8.3 % the low lignin parental genotype compared to Express 617 with 12 %. The reduction in lignin content was accompanied by a significant reduction of the seed coat proportion (Behnke et al. 2018).

Lignin is a large ubiquitous aromatic polymer and a major cell wall component. There are three main types of lignin, *p*-hydroxyphenyl lignin (H-type lignin), syringyl lignin (S-type lignin) and guaiacyl lignin (G-type lignin). All of which are formed by polymerisation of different phenylpropanoids, predominantly the monolignols *p*-coumaryl, coniferyl and sinapyl alcohols varying in their degree of methoxylation (Fraser and Chapple 2011, Ding et al. 2021, Vanholme et al. 2010). The first step in the general phenylpropanoid metabolism is catalysed by phenylalanine ammonia-lyase (PAL) which plays an important role in the biosynthesis of flavonoid- and lignin-related metabolites. The *PAL* gene family in *Arabidopsis thaliana* consists of four paralogues named *AthPAL1* – *AthPAL4* (Raes et al. 2003). Expression analysis of the *PAL* genes showed that *AthPAL3* is expressed at basal levels in stems, *AthPAL1, AthPAL2* and *AthPAL4* are expressed with high levels in stems of plants in later growth stages, while *AthPAL2* and *AthPAL4* are expressed in seeds (Fraser and Chapple 2011, Raes et al. 2003). In oilseed rape, the corresponding *B. napus* homologue *PAL4* was found as the key candidate gene associated with lignin biosynthesis, possessing a positive influence on seed lignin content (Behnke et al. 2018, Wang et al. 2015).

The objective of this work was to further investigate the major QTL found on C05 for low lignin content in the parental genotypes and progeny of the low fibre line SGDH14 and the high fibre line Express 617. In-depth sequence analysis was conducted by mapping ONT long-reads derived from SGDH14 against the Express 617 genome assembly (Lee et al. 2020). RNA-Seq data of developing seeds were used to analyse transcriptional changes between the two genotypes. The sequencing results revealed a homologous non-reciprocal translocation (HNRT) of an A05 sequence fragment replacing a homoeologous fragment on C05, which is associated with a reduction of lignin content in SGDH14.

## Material & Methods

### Plant material

The plant material comprised the *B. napus* genotypes SGDH14 and Express 617 which were previously described by Behnke et al. (2018). The oil-rich genotype SGDH14 is derived from the cross between the cultivars Sollux and Gaoyou analysed by Zhao et al. (2005). Express 617, widely used in mapping populations, is the inbred line 617 of the German winter oilseed rape cultivar Express with low erucic acid, low glucosinolate and high oil content (Behnke et al. 2018, Lee et al. 2020). Phenotypic characteristics of SGDH14 in contrast to Express 617 are its high erucic acid and glucosinolate contents, as well as low lignin content in the seeds (Behnke et al. 2018). SGDH14 and Express 617 were used for nGBS, RNA-Seq analysis and SGDH14 was used for ONT sequencing. Moreover, previous published 60K SNP data of Express 617 and SGDH14 was analysed (Behnke et al. 2018). Additional *B. napus* cultivars included in the Illumina 60K SNP and nGBS analysis are Adriana, Sollux, SGEDH13, and Gaoyou. Moreover, a F2 population of the cross SGDH14 and Express 617 consisting of 195 lines, further referred to as SGEF2 population, was used for PCR analysis of the HNRT event and for the determination of the seed coat proportion.

### Field experiments

The SGEF2 population was tested in a field experiment in one environment, Göttingen-Reinshof (51°29’51.6”N 9°55’55.2”E), in north-western Germany during the growing season 2019/2020. F1:2 seeds were sown in single row observation plots. Fertiliser, fungicides and insecticides were applied following standard schemes. Individual F2 plants (n=195) were marked in the field and small amount of leaf material was collected at flowering time, freeze dried and used for DNA extraction (see polymerase chain reaction (PCR) method section). At maturity, F2:3 seeds were harvested from each open pollinated plant. Seeds were cleaned, dried to a moisture content of 5 to 8 % and stored at a dry place at room temperature for further analysis.

### Seed quality analysis - determination of crude fibre contents

Near Infrared Reflectance Spectroscopy (NIRS) was applied to predict seed quality and fibre traits. NIRS measurements were performed using 3 g F2:3 seed samples of the SGEF2 population in small ring cups and the FOSS monochromator model 6500 (NIRSystem Inc., Silverspring, MD, USA). Between 400 and 2498 nm absorbance values log at 2 nm intervals were recorded to create a NIR spectrum for each sample. Using the WinISI software (Version 4.12.0.15440, FOSS Analytical A/S, Denmark) and applying the calibration raps2020.eqa provided by VDLUFA Qualitätssicherung NIRS GmbH (Kassel, Germany) oil and protein content were determined in percent at 91 % seed dry matter. Protein content in the defatted meal (PidM) was calculated by using NIRS predicted seed oil and seed protein content (both at 91 % dry matter) as: % protein in the defatted meal (PidM) = [% protein / (100 − % oil)] × 100. The sum of seed oil and protein content (oil + protein) was obtained by forming the sum of oil and protein content at 91 % seed dry matter. Glucosinolate is determined as μMol · g^-1^ at 91 % seed dry matter. Erucic acid is determined in %. The fibre components of the Neutral Detergent Fibre (NDF), Acid Detergent Fibre (ADF) and lignin in the defatted meal (Van Soest et al. 1991) were estimated using the calibration equation valid600div7.eqa (Dimov et al. 2012). The calibration was extended by adding 252 selected samples with a wide variation in seed fibre components. The hemicellulose and cellulose contents were calculated by subtracting ADF from NDF and lignin from ADF contents, respectively. In validation, the standard error of prediction corrected for the bias (SEP(C)) of the extended calibration was 1.78 %, 1.62 %, 0.93 %, 1.14 % and 1.48 % for NDF, ADF, lignin, hemicellulose and cellulose content, respectively, in the defatted seed meal.

### Determination of other seed traits

The degree of seed colour pigmentation was visually scored according to a scale from (1) yellow, (3) yellow brown, (5) brown, (7) dark brown and (9) for black and using intermediate scores (Widiarsih et al. 2021). Seed coat content was determined following the method described by Dimov et al. (2012). The seed coat proportion in % was determined from 100 seeds (approx. 500 mg) per genotype. The thousand-kernel weight (TKW) was obtained from weight conversion of 500 seeds.

### PCR and KASP assay

DNA was isolated following the standard protocol of innuPREP PlantDNA Kit (Analytik Jena GmbH, Germany) using the SLS lysis solution. All PCRs were performed in 25 μL reactions in Bio-Rad 96-well thermocyclers. Each reaction contained 16.8 μL ddwater (HPLC grad, Chemsolute, Th. Geyer GmbH & Co. KG Germany), 5 μL 5X Biozym Reaction Buffer, 0.2 μL Biozym HS Taq DNA Polymerase (5 u/ μL) (Biozym Scientific GmbH, Germany), 1 μL 10 mM per oligonucleotide and 1 µL 10 ng DNA was added. The used PCR strategies and corresponding oligonucleotides and applied PCR parameters are shown in Figure S1, Table S1 and S2 of Supplementary Information 1. Gelelectrophoresis was performed on an 1.2 % agarose gel and 100 V for 30 min. Based on provided sequences anchored KASP primers were ordered from LGC, Pindar Road, Hoddesdon, Herts, EN11 0WZ, UK. KASP assay was performed according to the manufacturer’s instructions.

### Identification of the C05 homologous region in the Express 617 genome assembly

For the identification of the homologous C05 region, 30 kbp up- and downstream of the described borders of the C05 major QTL (Behnke et al. 2018) were extracted based on the Darmor-*bzh* reference genome sequence v4.1 (Chalhoub et al. 2014). Next, these sequences were used for a BLASTn against the Express 617 assembly (Lee et al. 2020) to infer the localisation of the respective homologous C05 region.

### ONT genomic reads analyses of SGDH14

To analyse the major QTL for seed lignin content on chromosome C05 (Behnke et al. 2018) in SGDH14 on genomic level, Oxford Nanopore Technology (ONT) was used for the generation of long-reads derived from a single SGDH14 plant. Genomic DNA from one young leaf was extracted as described before (Siadjeu et al. 2020). DNA quality was assessed via NanoDrop and PicoGreen (Fluostar) measurement, as well as running a 1 % agarose gel. The DNA was prepared with the short-read eliminator (SRE) XL kit, following library construction with the LSK109 kit and then loaded onto a MinION/GridION R9.4 flow cell. After a nuclease flush another library, prepared with the LSK109 kit and derived from a sample prepared with the SRE-XL kit, was loaded onto the same flow cell. A total output of 3.4 Gbp passed reads were generated by this run. For the third library generation and sequencing on another MinION/GridION R9.4 flow cell, the DNA was used without prior short fragment depletion and a library was prepared with the LSK109 kit. A total output of 7.67 Gbp passed reads were generated by this run. All reads were filtered for a min qscore >7. After basecalling with Guppy v4.2.3 (Wick et al. 2019), reads were mapped to the *B. napus* Express 617 assembly (Lee et al. 2020) using minimap2 (Li 2018) yielding ∼10x genome-wide coverage. Minimap2 was run in ONT mode (-ax map-ont) and no secondary alignments were allowed (--secondary=no). BAM files were sorted and indexed via samtools (Li et al. 2009). Manually curation of regions of interest were performed by using IGV (Robinson et al. 2011) and BLAST (Altschul et al. 1990). Synteny plots were generated with jcvi/MCscan (Tang et al. 2008).

### Determination and molecular validation of HNRT border sequences

For the identification of the border sequences of the HNRT event, a k-mer approach was used. First, border-spanning reads of SGDH14 were divided into 15 bp k-mers. Afterwards, each k-mer was assigned to C05, A05, or considered unspecific. If a k-mer was present in both subgenomes (unspecific) it was excluded from further analyses, yielding *bona fide* subgenome-specific k-mers. These k-mers were then mapped back to the border-spanning read sequences to narrow down the exact position of the left and right substitution border. Finally, manual curation was used to extract subgenome-specific SNPs around the borders of the HNRT. These SNPs were further used for oligonucleotide design to validate the HNRT borders via Sanger sequencing (Table S1, Supplementary Information 1).

### Coverage analysis

For the genomic coverage analysis BAM derived coverage files were generated as described before (Pucker and Brockington 2018). Next, the genomic coverage of specific regions was compared to the chromosome-wide coverage of the corresponding chromosome to identify duplicated or deleted regions (Figure S2 and S3, Supplementary Information 2).

### Functional annotation and *in silico* gene prediction

Genes were functionally annotated by transferring the Araport11 functional annotation to the *B. napus* Express 617 gene models as described before (Schilbert et al. 2021). Not annotated genes in the Express 617 annotation were identified via an *in silico* gene prediction using AUGUSTUS v3.3.3 Web server (Stanke and Morgenstern 2005). The predicted gene structures can be accessed via Table S3 of Supplementary Information 3. The functional annotation was assigned via a BLASTp search against the Arabidopsis database. Finally, the genes were incorporated into the Express 617 structural annotation file (.gff3) for further analysis.

### nGBS analysis

To identify SNPs and genotyping *Brassica napus* cultivars, normalised Genotyping-by-Sequencing (nGBS) was used (LGC, Berlin, Germany). Genomic DNA was extracted from leaves of the *B. napus* cultivars Express 617, SGDH14, Sollux, Adriana, SGEDH13, and Gaoyou. For each genotype 150 bp paired-end reads were generated via Illumina on the NextSeq500/550. After assessing read quality by FastQC (Andrews 2010), reads were mapped to the *B. napus* Express 617 assembly (Lee et al. 2020) using BWA-MEM v.0.7.13 (Li 2013). Default parameters were applied and the –M flag was set to avoid spurious mappings. Mapping statistics were calculated via the flagstat function of samtools (Li et al. 2009) (Table S4, Supplementary Information 3).

### Gene expression analysis: RNA extraction, library construction, and sequencing

For the gene expression analysis total RNA was extracted with the innuPREP Plant RNA Kit according to the manufacturer’s instructions with the PL Buffer (Analytik Jena, Germany). Seed samples were harvested at 35 days after flowering (DAF). Three biological replicates per genotype (SGDH14 and Express 617) were used. The RNA quality was validated using NanoDrop and Agilent DNF-472 HS RNA (15 nt) Kit (Agilent Technologies, CA, USA) on the 5200 Agilent system to confirm the purity, concentration, and integrity, respectively. Based on 2 µg of total RNA, sequencing libraries were constructed with removal of rRNA through PolyA selection for mRNA species (eukaryotic). Paired-end sequencing of 150 bp was performed on an Illumina NovaSeq 6000 following the 2×150 bp configuration at the Sequencing Facility of GENEWIZ Germany GmbH, Leipzig, Germany.

### RNA-Seq read mapping and differential gene expression analysis

Read quality was analysed via FastQC v0.11.9 (Andrews 2010). After trimming the reads with Trimmomatic v0.39 (Bolger et al. 2014), the reads revealed an overall good quality with a phred score of minimum 36. Reads were then mapped to the Express 617 genome assembly (Lee et al. 2020) by STAR v2.7.1a (Dobin et al. 2013) using basic mode allowing maximal 5% mismatches per read length and a minimum of 90 % matches per read length (see Table S5 of Supplementary Information 3 for mapping statistics). The read mappings were used to manually correct the functional annotation of Express 617 including e.g. the previously not annotated A05 *PAL4* homologue and subsequently used for downstream analyses. We used featureCounts v2.0.1 (Liao et al. 2014) to generate count tables applying the -O, -B, -p, and -fraction parameters based on the improved Express 617 annotation (see above). Raw count tables were subjected to differentially expressed gene analysis via DESeq2 (Love et al. 2014) to identify differentially expressed genes (DEGs) between SGDH14 and Express 617 seeds (35 DAF). DEGs with a log_2_ fold change > 1 and an adjusted p-value < 0.05 were selected for downstream analysis.

### Organ-specific gene expression analysis using newly generated and publicly available *B. napus* RNA-Seq data

For the analysis of organ-specific gene expression across various organs newly generated (see above) and publicly available *B. napus* RNA-Seq data sets were used as described before (Schilbert et al. 2021). Transcript abundance and corresponding count tables were analysed and generated via Kallisto v0.44.0 (Bray et al. 2016). Heatmaps were constructed as described previously (Schilbert et al. 2021).

### Identification of *PAL* gene family in *B. napus* Express 617

The *PAL* homologues in *B. napus* Express 617 were identified via KIPEs as described previously (Pucker et al. 2020). To reduce the number of fragmented peptides derived from possible mis-annotations, KIPEs was run with a minimal BLAST hit similarity of 40 %. The functional described PAL sequence collection described in KIPEs was used as bait peptide sequences. The proteome file of *B. napus* Express 617 was used as subject. The alignment was constructed with MAFFT v7 (Katoh and Standley 2013) and trimmed to minimal alignment column occupancy of 10 %. Next, FastTree v2.1.10 (Price et al. 2009) was used to build a phylogenetic tree using 10,000 rounds of bootstrapping. Visualisation of the phylogenetic tree was done with FigTree v1.4.3 (Rambaut 2006) (Figure S4, Supplementary Information 4). The *B. napus PAL* sequences were classified based on the corresponding *A. thaliana* homologues and their amino acid and coding sequences can be found in Table S6 of Supplementary Information 4.

### Development of a *PAL* specific marker

For the development of a marker, which can be applied in plant breeding, the subgenome-specific SNPs of the A05 and C05 *PAL4* homoeologues were harnessed. By using oligonucleotides (Table S1, Supplementary Information 1), which flank the subgenome-specific SNPs in the second exon, a subgenome-specific marker for the presence of the A05 and/or C05 *PAL4* homoeologue was developed.

## Results

### Phenotypic analysis – the F2 population derived from a cross of the low lignin line SGDH14 and high lignin line Express 617 reveals variation in fibre traits

SGDH14 had lower seed lignin content and higher contents of oil and the sum of oil and protein compared to Express 617 (Table 1). In the SGDH14 x Express 617 F2 population, namely SGEF2, the lignin content of seeds ranged from 3.6 to 13.8 %, hemicellulose from 1.6 to 7.9 % and cellulose from 12.5 to 17.7 % in the defatted meal, respectively (Table 1). Lignin showed a strong negative correlation to hemicellulose and a weak positive correlation to cellulose (Figure 1B). Particularly, the lignin content showed a bimodal, skewed distribution. The low and high lignin genotypes showed a 1:2 segregation (Figure 1A). Comparing the low and high lignin groups of the SGEF2 population, the low lignin group showed significantly higher contents of oil and protein in the defatted meal and hemicellulose, as well as for the sum of oil and protein compared to the high lignin group. The oil and protein content ranged from 38.7 to 47.1 % and from 17.1 to 22.3 %, respectively. The protein content in the defatted meal ranged from 31.4 to 37.4 % and the sum of oil and protein content varied between 60.6 and 66.1 %. The thousand-kernel weight (TKW) ranges from 3.8 to 6.8 g and is significantly higher in the low lignin group (Table 1). All those traits showed a weak negative correlation to lignin (Figure 1B). A positive correlation between lignin and darker seed colour as well as seed coat content was found (Figure 1B). The pigmentation of SGEF2 seed ranged from dark brown (7.0) to black (9.0) (Table 1). Transgressive segregation was observed for most of the traits in the F2 population. The high lignin group exhibited a significantly higher seed coat content (16.1 %) than the low lignin group (14.1 %; Table 1 and Figure 1C). To summarise the SGEF2 population was extensively phenotypically characterised and showed a large variation in seed lignin content, which was also identified in the parents with SGDH14 possessing a low lignin content and the Express 617 showing a high lignin content.

**Table 1:** Phenotypic trait measurements in the SGEF2 population and the respective low and high lignin groups. Minimum, maximum, mean values and standard deviation (Stdev) for contents of seed oil (%), protein (%), protein in defatted meal (PidM in %), the sum of oil and protein (%), erucic acid (%), glucosinolate (μMol · g^-1^ at 91 % seed dry matter), the fibre fractions (% in defatted meal), thousand-kernel weight (TKW) (g), seed colour and seed coat content (%) of the SGEF2 population (n=195). The lignin groups are divided at a lignin content of <7.5 % into low lignin group (n=67) and high lignin group (n=128). The mean values are shown. Significance levels were analysed with single factor ANOVA.

| Traits | Min | Max | Mean | Stdev | Lignin groups |  |  | Parental genotypes |  |
| --- | --- | --- | --- | --- | --- | --- | --- | --- | --- |
|  |  |  |  |  | Low |  | High | SGDH14 | Express 617 |
| Oil | 38.7 | 47.1 | 43.3 | 1.2 | 43.8 | ** | 43.0 | 46.2 | 42.7 |
| Protein | 17.1 | 22.3 | 19.6 | 0.8 | 19.6 | ns | 19.6 | 18.2 | 18.5 |
| PidM | 31.4 | 37.4 | 34.5 | 1.0 | 34.9 | ** | 34.3 | 33.9 | 32.3 |
| Oil + Protein | 60.6 | 66.1 | 62.9 | 0.9 | 63.4 | ** | 62.6 | 64.4 | 61.2 |
| Erucic acid | 0.0 | 27.4 | 13.8 | 6.0 | 13.4 | ns | 14.0 | 15.4 | 0.0 |
| Glucosinolate | 13.9 | 61.5 | 41.8 | 9.1 | 42.4 | ns | 41.4 | 44.1 | 13.7 |
| Lignin | 3.6 | 13.8 | 8.8 | 2.2 | 6.0 | ** | 10.3 | 7.2 | 10.7 |
| Hemicellulose | 1.6 | 7.9 | 6.1 | 0.8 | 6.7 | ** | 5.7 | 6.6 | 5.7 |
| Cellulose | 12.5 | 17.7 | 15.5 | 0.8 | 15.4 | ns | 15.5 | 15.4 | 16.0 |
| TKW | 3.8 | 6.8 | 5.2 | 0.5 | 5.3 | ** | 5.1 | 4.5 | 4.7 |
| Colour | 6.0 | 9.0 | 8.3 | 0.9 | 7.5 | ** | 8.8 | 7.8 | 9.0 |
| Seed coat | 11.2 | 25.3 | 15.2 | 2.2 | 14.1 | ** | 16.1 | 12.1 | 16.3<sup>§</sup> |

ns, \*, \*\* shows not significant and significance with $p < 0.05$ und $p < 0.01$
<sup>§</sup> data taken from Behnke et al. (2018)

**Figure 1:**
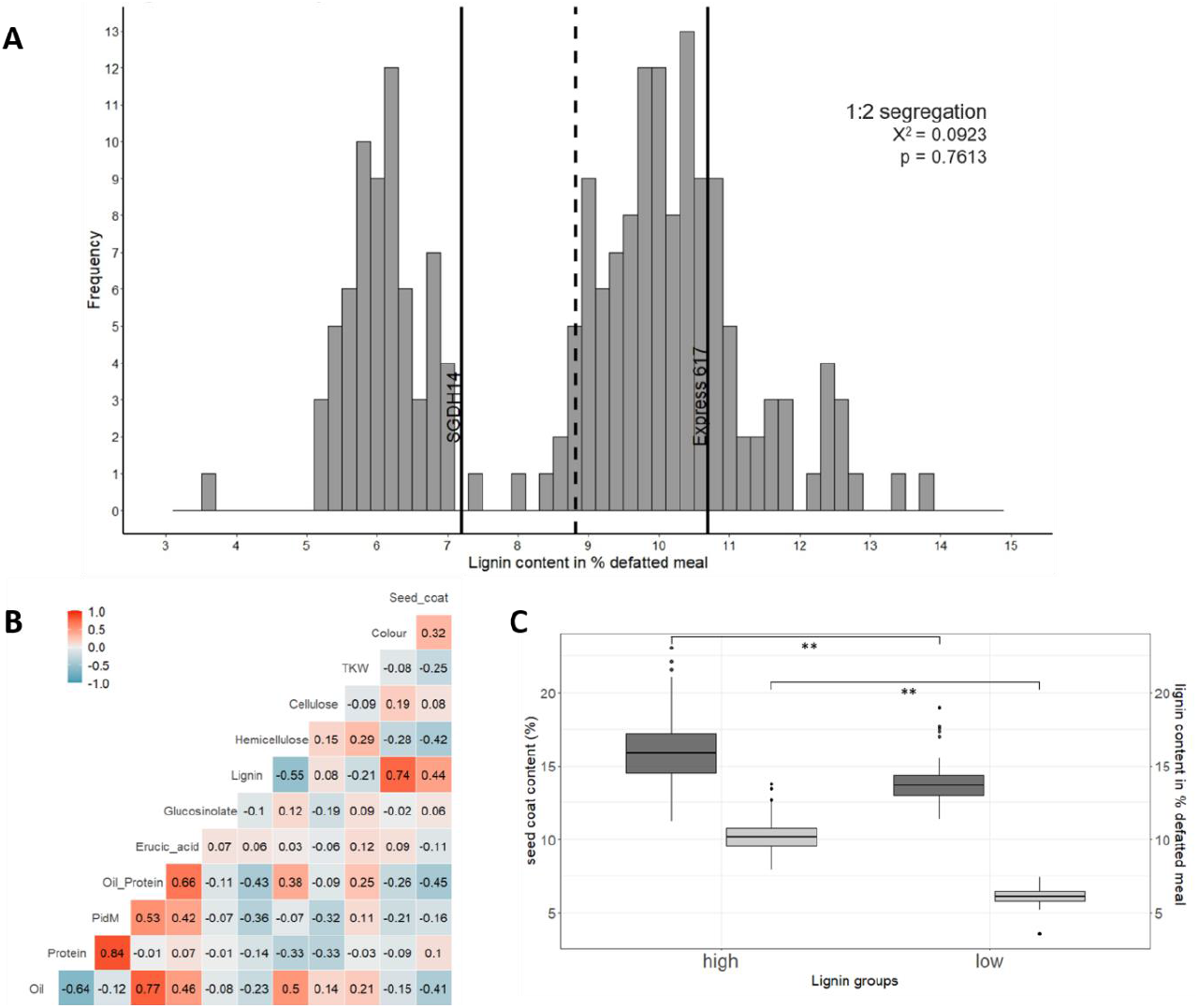
Phenotypic characterisation of the SGEF2 population. (A) Frequency distribution of lignin content in defatted meal of the SGEF2 population (F2:3 seeds). The lignin content of the parental genotypes SGDH14 and Express 617 is indicated by black vertical lines and the grey vertical dashed line marks the mean lignin content of the SGEF2 population (n=195). The chi-square result for a 1:2 segregation is displayed. The p-value is not significant so that H0, the 1:2 segregation, is accepted. (B) Correlation Matrix of the SGEF2 population (F2:3 seeds). Positive correlation is indicated in red and negative correlation is indicated in blue. The correlation coefficients are given for each trait. PidM is protein content in the defatted meal at 91 % dry matter. (C) Seed coat and lignin content (%) in high and low lignin groups of the SGEF2 population. The high lignin group (n=128) is shown on the left and the low lignin group (n=67) is displayed on the right side of the figure. Dark grey coloured boxplots represent the seed coat content and the light grey boxplots show the lignin content. The significance level was calculated with a t-test. ** shows significance p < 0.01.

### Detailed characterisation of the physical position of the major low lignin QTL

To analyse the genetic basis underlying the observed phenotypic variation of the parents, we used the genome assembly of the high lignin parent Express 617 (Lee et al. 2020) and generated genomic long-reads of the low lignin parent SGDH14. As a major QTL for low lignin content on C05 was identified before, based on the genome assembly of Darmor-*bzh* (Behnke et al. 2018), we first aimed to identify the syntenic genomic region in the Express 617 genome sequence. The physical position of the major low lignin QTL in the genome assembly of Express 617 ranged from 41,286,363 bp to 42,386,439 bp (∼1,100 kbp) on C05, while in Darmor-*bzh* the QTL ranged from 39,713,411 bp to 40,616,480 bp (∼903 kbp) (Figure 2). Thus, the QTL region is ∼197 kbp bigger in the Express 617 genome assembly. To identify genomic differences located near or inside the major QTL between the parental genotypes, the genomic long-reads of SGDH14 were mapped against the Express 617 genome assembly. A ∼208 kbp region ranging from 41,563 to 41,771 kbp on C05 located near the center of the major QTL revealed no read coverage in SGDH14 (Figure 2; Figure S2 of Supplementary Information 2). Corresponding flanking reads of SGDH14 were identified to be chimeric, having a C05 and an A05 sequence part. By separating the chimeric reads in subgenomic k-mers (Figure S3, Supplementary Information 2), the genomic border sequences were narrowed down and validated via Sanger sequencing (Figure 2). Ultimately, these results showed that the homoeologous A05 sequence ranging from 27,121 kbp to 27,289 kbp was inserted between 41,563 kbp and 41,771 kbp on C05 in SGDH14 (Figure 2). Further, this homoeologous A05 locus was identified to be duplicated, as the average coverage of the A05 homoeologous locus was ∼2-fold higher, compared to the average coverage of the whole A05 chromosome (Figure S2, Supplementary Information 2). Therefore, the name A05’’ is used for sequence features located within this duplicated homoeologous A05’’ region (Figure 2). A homoeologous non-reciprocal translocation (HNRT) is identified in SGDH14.

**Figure 2:**
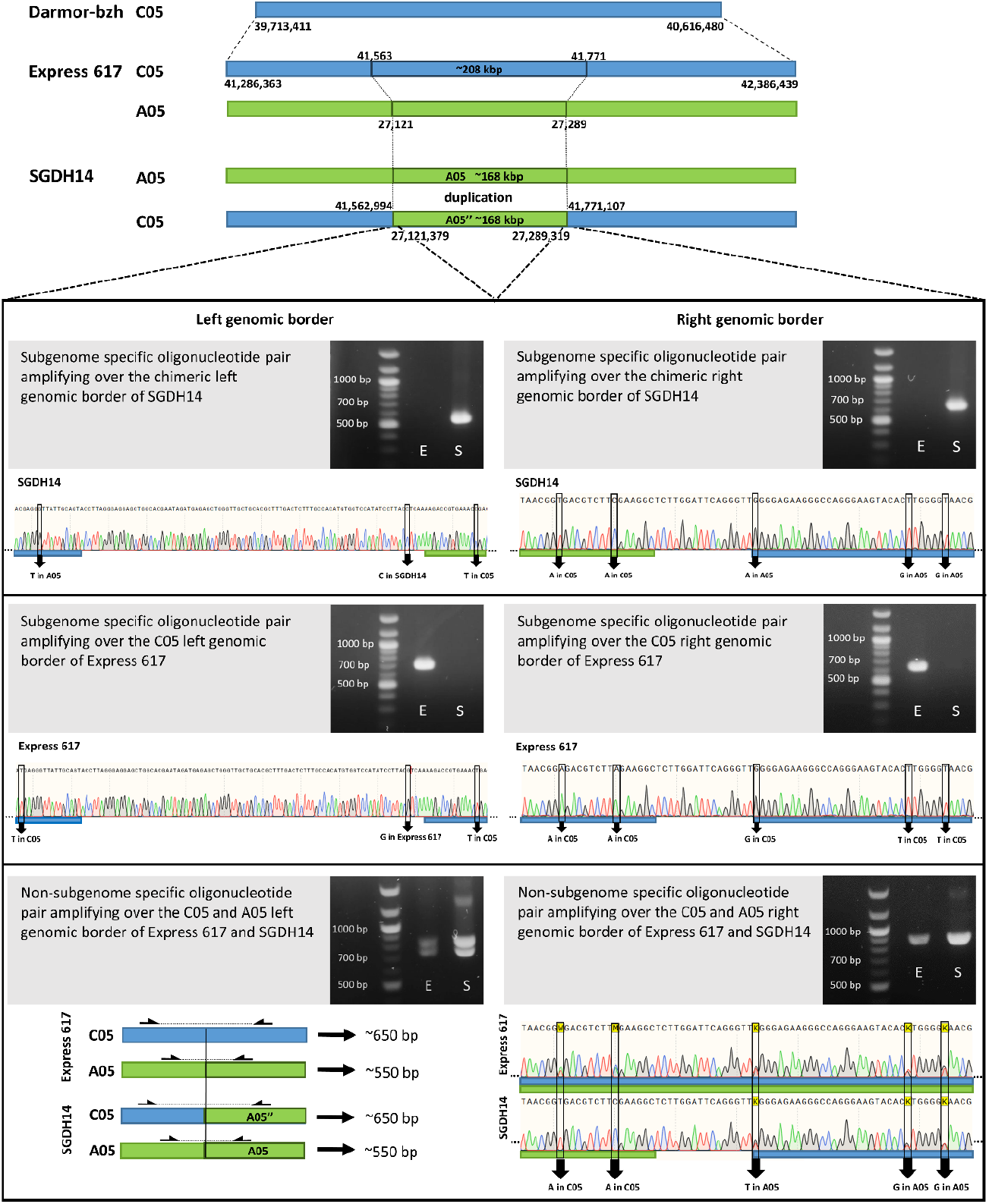
SGDH14 reveals a homoeologous non-reciprocal translocation in the major low lignin QTL region on C05. (Top) The location of the major low lignin QTL on C05 (blue) in the Darmor-*bzh*- and the Express 617 genome assembly is shown and the A05 (green) to C05 homoeologous non-reciprocal translocation region in SGDH14 is marked with a dark green outlined rectangle and dotted lines. (Bottom) Subgenome-specific and non subgenome-specific oligonucleotides were used for the validation of the substitution borders. The amplified products using Express 617 (E) and SGDH14 (S) genomic DNA, as well as sections of corresponding representative Sanger sequencing results are shown. Figure is not to scale.

Importantly, in the Express 617 genome assembly on chromosome A05 at position 27,213,244 to 27,213,343 a stretch of ambiguous bases was identified. This is indicative of an assembly break that could not be resolved in the Express 617 genome assembly. In accordance, the ONT long-reads of SGDH14 revealed a highly repetitive region located at this position inside the HNRT region, which could not be spanned. Although border spanning chimeric reads reaching into this repeat could be identified, their length was not sufficient to span this large repeat (Figure S3, Supplementary Information 2). As the region sizes were estimated based on the Express 617 genome assembly, irrespective of this repeat, they are likely underestimating the true sizes. The nGBS data support all above described findings. In summary, in SGDH14 a ∼208 kbp region on C05 of the major QTL is deleted and is substituted by a ∼168 kbp duplication of the homoeologous A05’’ sequence.

### HNRT event in the SGDH14 x Express 617 DH population explains seed lignin content

To analyse the association between the HNRT event and the lignin phenotype we screened a second independent population for the presence or absence of the HNRT event. We used the SGDH14 x Express 617 DH population of Behnke et al. (2018). SGDH14 shows the low lignin phenotype and harbours a HNRT event with a duplication of an A05 chromosome part and replacing the homoeologous C05 region. Analysis of 60K Illumina SNP data revealed that several markers in the QTL region were scored as “failed” not only in SGDH14, but also in its ancestral genotypes Sollux and Gaoyou, as well as in SGEDH13 and Zheyou 50, but not in Express 617 and Adriana (Table S8, Table S9, Supplementary Information 5). Carefully studying the frequency distribution of “failed” scored markers in the C05 region in the original SGDH14 x Express 617 DH genotyping table, allowed a clear separation of DH lines with and without “failed”-markers, i.e. with and without HNRT. Comparison of the DH lines with and without the HNRT event showed a clear, significant difference in several quality traits (Table 2). The lignin content was significantly reduced in lines with the HNRT event, whereas hemicellulose and cellulose showed increased contents. Oil and protein content in defatted meal and the sum of oil and protein content were increased in the low lignin lines. The seed protein content was not different between the groups. These results were concordant with the results of the high and low lignin groups of the SGEF2 population (Table 1).

**Table 2:** Phenotypic trait measurements in the SGDH14 x Express 617 DH population and the respective lines with and without HNRT. The groups with and without HNRT were formed by dividing the DH lines according to their SNP marker data in the C05 QTL region. Genotypes with HNRT showed several markers as “failed” in this region, equal to missing marker pattern of SGDH14. Oil, protein and crude fibre contents of the genotypes with the HNRT (n=71) and without HNRT (n=67) in the SGDH14 x Express 617 DH population of Behnke et al. (2018) are listed. The significance level was calculated with a single factor ANOVA.

| Traits | Range with HNRT | Range without HNRT | Genotypes with HNRT |  | Genotypes without HNRT |
| --- | --- | --- | --- | --- | --- |
| Oil | 43.0 - 50.8 | 42.5 - 48.6 | 47.0 | ** | 45.6 |
| Protein | 16.6 - 19.5 | 16.6 - 19.5 | 17.9 | ns | 17.9 |
| PidM | 30.3 - 36.9 | 30.3 - 36.3 | 33.8 | ** | 32.9 |
| Oil + Protein | 60.8 - 68.3 | 60.9 - 67.0 | 64.9 | ** | 63.5 |
| Erucic acid | 0.0 - 51.0 | 0.0 - 49.5 | 22.9 | ns | 20.2 |
| Glucosinolate | 18.2 - 74.6 | 18.7 - 69.0 | 53.1 | ns | 49.4 |
| Lignin | 6.1 - 9.8 | 8.4 - 14.0 | 7.3 | ** | 11.4 |
| Hemicellulose | 11.5 - 15.1 | 11.0 - 13.3 | 14.0 | ** | 12.4 |
| Cellulose | 10.7 - 12.5 | 9.9 - 11.6 | 11.4 | ** | 10.8 |
| TKW | 4.4 - 6.0 | 4.1 - 5.8 | 5.1 | ns | 5.0 |

Oil, Protein, Oil+Protein in % 9 % moisture, PidM = Protein in % in defatted meal; Hemicellulose, Cellulose und Lignin (% in defatted meal); \*\* shows significance with $p < 0.01$ .

### Candidate genes associated with phenylpropanoid and lignin biosynthesis

In order to identify candidate genes that might contribute to the observed lignin phenotype, e.g. involved in general phenylpropanoid and lignin biosynthesis, we screened the genomic regions affected by the HNRT event. Thus, the deleted region of the C05 chromosome and the substituted and thus duplicated A05’’ region were inspected for candidate genes. Based on the Express 617 genome assembly, 207 genes are located inside the major low lignin QTL region on C05 (41,286 to 42,386 kbp) of which 32 genes are deleted by the HNRT event (41,563 to 41,771 kbp) (Table S10, Supplementary Information 6). Additionally, *C05p041220*.*1* (homologue of *AthSEC8*) and *C05p041550*.*1* (homologue of *AthPIP5K9*) are located at the genomic borders of the HNRT event, leading to the loss or disruption of 26 of its 27 exons in the case of *C05p041220*.*1* and part of the first exon in the case of *C05p041550*.*1*. The homoeologous A05 region (27,121 to 27,289 kbp) duplicated in SGDH14 contains 31 genes based on the Express 617 genome assembly and shows a large proportion of overlapping genes to its C05 homoeologous region (Table S11, Supplementary Information 6).

We analysed the organ-specific expression of candidate genes, which *A. thaliana* homologues are associated with lignin biosynthesis (Table S12, Supplementary Information 7). Most candidate genes were very low or not at all expressed in Express 617 and SGDH14 seeds (35 DAF) except for the C05 *B. napus* homologue *AthPAL4 C05p041260*.*1*, which showed the highest expression in Express 617 but not in the HNRT deleted region of SGDH14. Moreover, the *B. napus* homologues of *AthCESA3* (*C05p040780*.*1*), *AthGALAK* (*C05p040910*.*1*), *AthMUCI21* (*C05p041270*.*1*), and *AthDEX1* (*C05p042190*.*1*) were the highest expressed candidate genes during seed coat development. In line with the HNRT event, *AthPAL4* (*C05p041260*.*1*) and three other genes located in the HNRT region were identified as lower expressed in SGDH14 compared to Express 617 seeds (Table S13, Supplementary Information 7), namely the *B. napus* homologues of *AthSEC8* (*C05p041220*.*1*), *AthSDP6* (*C05p041230*.*1*) and *AthMUCI21* (*C05p041270*.*1*).

### SGDH14 harbours two A05 *BnaPAL4* homologues

Due to the allotetraploid origin of *B. napus* homoeologous gene copies are frequent and could compensate for the loss of one homoeologue gene copy. Synteny analysis and manual inspection of RNA-Seq read coverage revealed an A05 *PAL* homoeologue in Express 617, which was not included in the original Express 617 annotation and named *A05p031299*.*1* (Figure S5, Supplementary Information 4). The respective gene was identified as *PAL4* homologue by *in silico* gene prediction followed by a BLASTp search. The organ-specific expression analysis revealed that both *BnaPAL4* homoeologues are much higher expressed in developing seeds than the other identified paralogues of Express 617 (Table S7, Supplementary Information 4).

In SGDH14, two A05 *PAL4* copies (A05 and A05’’ *PAL4*) were identified by inspecting SNPs of the genomic and transcriptomic SGDH14 read mappings against the Express 617 genome assembly. In total, 16 SNPs (12 in exon No. 3, 3 in 3’UTR, 1 upstream of 5’UTR) revealed a ∼50:50 ratio of reference vs alternative alleles (Figure 3, Figure S6 of Supplementary Information 4). Twelve of these SNPs are located in the third exon, of which six SNPs are identical to the C05 *PAL* allele, while the remaining six were specific for the new A05’’ *PAL4* allele as they differ from the A05 and C05 *PAL4* allele (Figure S6, Supplementary Information 4). Some of these SNPs result in amino acid exchanges compared to the A05 PAL4 amino acid sequence, of which three are specific to the A05’’ PAL4 (V586I, D596E, V638I), while the remaining two (E610G, V629G) are identical to the C05 PAL4 sequence. Analysis of the putative promoter sequence using 1 kbp upstream of the translational start side showed only one SNP differentiating the A05’’ *PAL* from the A05 *PAL* allele, while several subgenome-specific SNPs were identified (Figure S6, Supplementary Information 4). Finally, a marker for the presence or absence of the C05 *PAL4* homoeologue was designed by using non subgenome-specific oligonucleotides. Sequencing the PCR amplicons derived from the second exon showed the absence of C05 specific SNPs in SGDH14, while Express 617 revealed heterozygous SNPs for each homoeologue (Figure 3). In accordance with the genomic results stressing the loss of the C05 *PAL4* homologue in SGDH14, the C05 *PAL4* was not expressed in SGDH14 but in Express 617. The two A05 *PAL4* homologues of SGDH14 revealed a ∼1.5 higher expression compared to the combined expression of the C05 and A05 *PAL4* homoeologues of Express 617 (Table S7, Supplementary Information 4).

**Figure 3:**
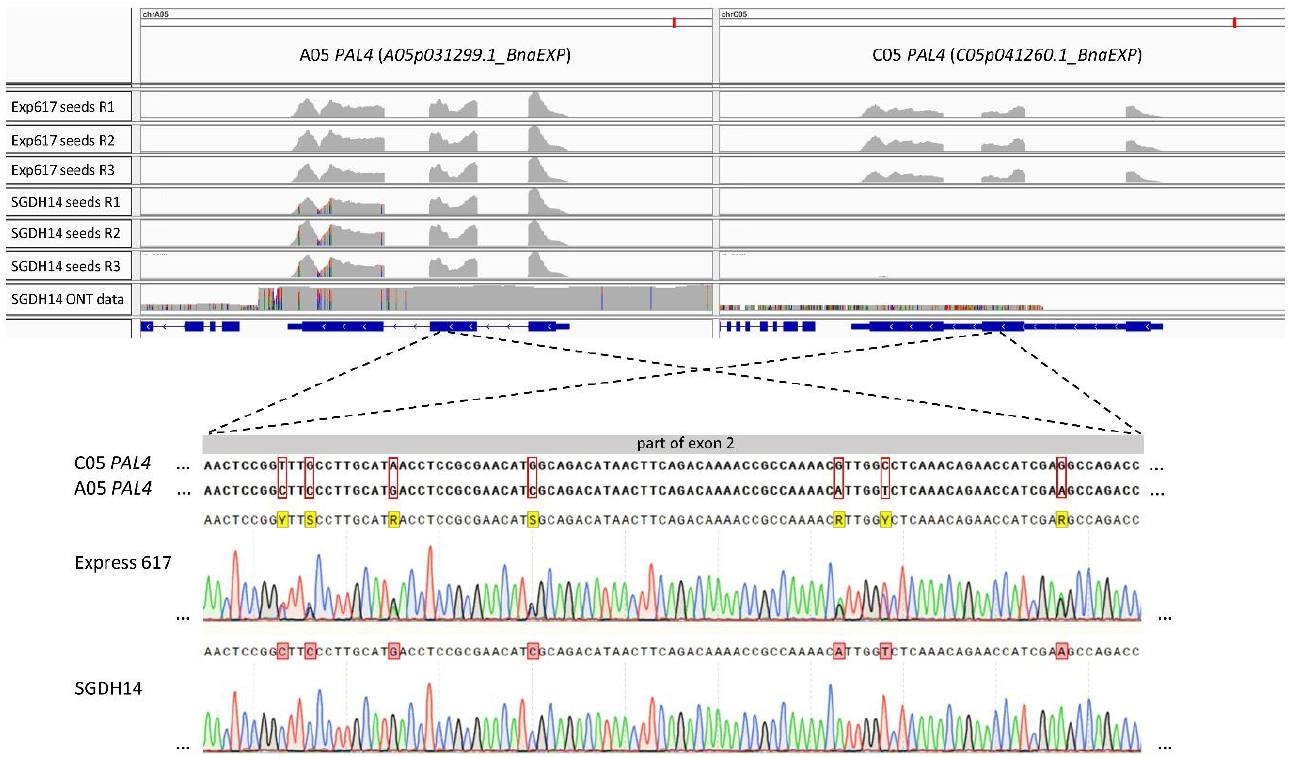
SGDH14 harbours a second A05 *PAL4* homologue, namely A05’’ *PAL4*, expressed in seeds, while the C05 *PAL4* is deleted. (Top) RNA-Seq data from seeds (35 DAF) of Express 617 and SGDH14 as well as ONT read mapping of SGDH14 against the Express 617 genome assembly. The three biological replicates of the RNA-Seq data are marked with R1, R2, and R3, respectively. Positions of SNPs differentiating from the Express 617 genome assembly are coloured as follows: adenine green, thymine red, cytosine blue, and guanine black. Twelve of sixteen identified SNPs are located in the third exon, of which six were specific for the new A05’’ *PAL4* allele as they differ from the A05 and C05 *PAL4* allele. (Bottom) Sequencing results of a part of the *PAL4* second exon using oligonucleotides, which can bind to both subgenomes, with Express 617 and SGDH14 gDNA. The C05 *PAL4* second exon sequence was used as reference and subgenome-specific SNPs are marked with a red rectangle. A representative sequenced section is shown.

## Discussion

### HNRT in *B. napus* and phenotypic changes

Widespread segmental duplications, deletions and homologous chromosome exchanges are already identified in a wide range of natural and synthetic genotypes of *B. napus*. Due to the presence of two additive subgenomes with high degree of homoeology, the meiotic chromosome pairing between homoeologous chromosomes causes the occurrence of homoeologous exchange (HE) and gene conversions. These homoeologous exchanges are the main drivers of diversification in polyploid genomes (Szadkowski et al. 2010, Chalhoub et al. 2014, Gaeta and Pires 2010). Changing the genome may subsequently cause modifications in the phenotypic appearance. The impact of homologous non-reciprocal translocation (HNRT) and HEs on the phenotype in *B. napus* were shown earlier. Seed yield, seed quality, seed colour, seed fibre content, flowering time, disease resistance and fertility were affected by homoeologous exchanges (Hurgobin et al. 2018, Schiessl et al. 2019, Stein et al. 2017). Looking at fibre content, Stein et al. (2013) concluded in their *B. napus* QTL mapping population that a HE causing a deletion or conversion is responsible for a lower seed fibre content and a yellow-coloured seed. The HE and subsequently important candidate genes were identified on chromosome A09. With higher marker density and sequence coverage, the borders and genetic position of the HNRT were confirmed (Stein et al. 2017). In the present study, the major QTL for lignin content on chromosome C05 (Behnke et al. 2018) was analysed in depth. With ONT- and RNA-Seq data, it was shown that HNRT of a A05 region replaces a homologous C05 region in SGDH14. This translocation was accompanied by a segmental C05 chromosome deletion and is responsible for the observed reduction in lignin content. The replacement of the C05 *BnaPAL4* by a A05 *BnaPAL4* homologue was identified as the likely cause for the observed changes in lignin content.

### A chromosome segment replaces C chromosome segment

As shown in previous studies, larger C-subgenome segments are replaced by smaller, homoeologous A-subgenome segments, driving postpolyploidisation genome size reduction (Samans et al. 2017, Sochorová et al. 2017, Higgins et al. 2018). Further, most of those chromosomal rearrangements occurred between regions of primary homology (Nicolas et al. 2007). In the present study, the allotetrapolyploid *B. napus* shows similar chromosomal rearrangements with a subgenome bias. In SGDH14 the 208 kbp large C05 chromosome segment was replaced by a smaller at least 168 kbp segment of the homoeologous A05 chromosome. The exact size of this segment could not be determined since a large repeat in this sequence region was not spanned by the long-read sequencing data. In the Express 617 genome assembly this repeat is marked as a stretch of ambiguous bases (Lee et al. 2020). Although the Express 617 genome assembly is based on 54.5x coverage with Pacific Biosciences long-reads, linked reads, optical map-, and high-density genetic map data (Lee et al. 2020) this repeat cluster could not be resolved. With the long-read data of SGDH14 the repeat could also not be resolved due to the lack of spanning reads. To determine the exact size of the HNRT, longer reads which span the repeat cluster are necessary.

### Method for QTL and HE detection

The major QTL C05 was genotyped and mapped previously with the Illumina Infinium Brassica 60K SNP array with a total number of 58,464 markers. The relevant region on C05 with the deleted fragment marked is shown in Table S8 along with the most likely and second most likely position of the SNP markers of Express 617 in Table S9 of Supplementary Information 5. Within the deleted C05 region only two SNP markers were polymorphic (Behnke et al. 2018). Other SNP markers within the C05 confidence interval were scored in SGDH14 as “failed” markers, indicating the HNRT. The DH population of Behnke et al. (2018) did not show a significant segregation distortion for the presence (n=71) and absence (n=67) of the HNRT. The DH lines with the HNRT had a significantly lower seed lignin content than the ones without HNRT (Table 2).

When did the HNRT event occur in SGDH14? To answer this question, the Illumina 60K SNP marker data of the parental genotypes Sollux and Gaoyou of SGDH14 were analysed. Based on the SNP marker data it became obvious that the same SNP markers gave “failed” results for the HNRT region on C05 in both parents. Furthermore, SGEDH13, an offspring of SGDH14 x Express 617, showed the same lignin QTL on C05 and the same “failed” results for the HNRT region on C05 (Yusuf et al. 2022). The same was found for the Chinese cultivar Zheyou 50 and Yusuf et al. (2022) discussed that Zheyou 50 may be an offspring from the SGDH14 x Express 617 cross, since it originates from the Zhejiang Academy of Agricultural Sciences as SGDH14, and carries the same major QTL on C05 and is of canola quality. Although the HNRT event in Sollux, Gaoyou and Zheyou 50 was not confirmed by sequencing, it appears likely that the HNRT event occurred much earlier and indicates that a positive selection may have led to its enrichment. The nGBS data indicated for Sollux and Gaoyou the HNRT event, but the exact borders could not be determined. The HNRT in Sollux is potentially even bigger (∼610 kbp) than in SGDH14. Yield testing of selected high and low lignin bulks of the crosses SGDH14 x Express 617 and of Adriana x SGEDH13 recently indicated an advantage of the low lignin over the high lignin bulks, since a higher yield could be detected (Holzenkamp et al. 2022).

As a consequence of genomic rearrangements, skewed marker segregation patterns often prevent standard mapping procedures accurately localising QTL and causal genes in these regions (Stein et al. 2017). It was proposed to use SNP marker data from loci spanning deletions or duplications, which would be discarded in standard mapping procedures (Stein et al. 2017, Gabur et al. 2018). With this data structural chromosome variation can be identified in QTL analysis and single nucleotide absence polymorphisms (SNaPs) can provide an added value. SNaPs should be used when one parent and progenies of the mapping population show missing markers. The search for presence–absence variation should be implemented in a routine screening to find diversity in breeding material (Stein et al. 2017, Gabur et al. 2018). Surprisingly, the HNRT event was accurately localised by the two SNP markers Bn-scaff_17441_1-p950045 and Bn-scaff_17441_3-p28950. SNP Bn-scaff_17441_1-p950045 was mapped next to Bn-scaff_17441_3-p28950, which are located within the confidence interval of the major lignin QTL on C05 identified by Behnke et al. (2018). Because Bn-scaff_17441_1-p950045 is located within the deleted C05 region based on its physical position in the Express 617 assembly, it should have given a “failed” result for about half of the DH population as the other markers in the HNRT region. However, this was not the case. This marker segregated 1 to 1 in the DH population (Table S8 in Supplementary Information 5).

Obviously, Bn-scaff_17441_1-p950045 detected in homozygous DH lines with the C05 deletion rather the homoeologous SNP on the SGDH14 A05 chromosome, resulting in a 1 to 1 segregation which fits to the presence/absence of the HNRT in the DH population. The same may be valid for the second SNP marker Bn-scaff_17441_1-p958418. The physical position of the second QTL flanking marker Bn-scaff_17441_3-p28950 is located downstream of the C05 deleted region.

### Lignin and seed coat content

In the present study the F2:3 seeds of the SGEF2 population showed a similar bimodal distribution of lignin content as in the DH population SGDH14 x Express 617 of Behnke et al. (2018). Although the SGEF2 population was only tested in one field environment, the results were comparable to those of Behnke et al. (2018). The chi-squared test revealed a 1:2 segregation, which deviates from the expected 1:3 segregation in F2 populations for a recessive mutation. This indicates a segregation distortion towards homozygous genotypes with HNRT. It was not possible to clearly identify and separate homozygous and heterozygous F2-plants for the HNRT translocation, because the designed PCR Primers were not specific enough. The genetic causes for the segregation distortion in the SGEF2 population remains unclear. Later on, designed anchored KASP INDEL markers were able to detect the A05/C05 insertion using subgenome specific polymorphisms on the right border (Figure 2, Figure S7 and S8 of Supplementary information 8). This marker could, however, not anymore be applied to the DNA samples of the F2 plant population.

In general, yellow seeded cultivars show a reduced lignin and seed coat content compared to black seeded cultivars (Wang et al. 2015, Ding et al. 2021). However, the seed material in this study revealed a rather low variation in seed colour, despite showing a wide variation in lignin content. The pigmentation of F2:3 seeds ranged only from dark brown to black, with a tendency to a lighter colour for the low seed lignin genotypes. Hence, the variation in lignin content should be correlated with the seed coat proportion. It was already previously reported, that the seed coat in high lignin lines was thicker than that of low lignin lines (Behnke et al. 2018, Ding et al. 2021, Wang et al. 2015). The seed coat content of SGDH14 (12.1 %) is significantly reduced by a quarter compared to Express 617 (16.3 %) (Behnke et al. 2018). Similar results were determined in this study with the SGEF2 population. The high and the low lignin groups of F2:3 seeds showed significant differences in seed coat content (Figure 1C).

The seed coat consists mainly of three layers from inside to outside: the mucous epidermal cells, the palisade and the endothelial layers (Moïse et al. 2005). Further, Ding et al. (2021) suggested that lignin is deposited in the palisade cells, thereby increasing the thickness of the palisade layer and the overall thickness of the cell wall. This can explain the seed coat thickness, but not the seed coat colour, since in our material the seed colour is similar despite varying seed lignin contents. The material of Yin et al. (2022) showed similar results with no correlation between seed coat thickness and seed colour. Generally, the seed coat should be thin to reach high oil contents. Those genotypes with a thinner seed coat are mostly yellow seeded cultivars. In the present study, this was not the case since SGDH14 showed a dark seed colour and a high oil content. Hu et al. (2013) showed similar results with black seeded oilseed rape lines which were reported to have high oil contents. It was shown by Iwanowska et al. (1994), that the innermost layer of the inner integument, known as the endothelium, accumulates phenolic compounds starting from the early stages of embryo development.

Flavonols and proanthocyanidins (PAs) are derived from the general phenylpropanoid pathway and are end products of different branches of the flavonoid biosynthesis. PAs are the major pigments responsible for the seed colouring in *A. thaliana* and *Brassica* species pathway (Xu et al. 2014). Seed pigments in mature dry seed of *Brassica* species are primarily oxidised PAs (Durkee 1971), which are responsible for the dark seed colour developed during seed maturation (Moïse et al. 2005). SGDH14 exhibits low lignin contents, but similar seed coat pigmentation as the high lignin line Express 617. The absence of C05 *BnaPAL4* expression and thus likely activity in SGDH14 could be an indication that the C05 *BnaPAL4* is specific for the lignin biosynthesis, while the A05 *BnaPAL4* could be involved in PAs synthesis.

In the present study, the high and low lignin groups (Table 1) showed clearly differentiating lignin contents. The overall lignin content in those genotypes was still higher than in the study of Ding et al. (2021) (with min. 0.94 %; max. 4.7 % lignin). The lignin content and seed coat colouring of our low lignin group with the high lignin group of Ding et al. (2021), were similar. This leads to the assumption that yellow seed coat colour appears, when the lignin content drops below 2 %. Moreover, a negative correlation between seed lignin and hemicellulose content was detected. The low lignin lines had a significantly higher hemicellulose content compared to high lignin lines (Table 1). In previous studies it was reported that the synthesis of cellulose and hemicellulose redirects photosynthetic assimilates from oil and protein into sugar biosynthesis, which may result in reduced contents of these two compounds (Wang et al. 2015, Liu et al. 2013). The hypothesis is, that assimilates which are not used in the general phenylpropanoid pathway to produce lignin are redirected into hemicellulose production. Similar redirection to hemicellulose synthesis was reported in rice (Ambavaram et al. 2011) and in aspen trees (Hu et al. 1999). Obviously, there is a crosstalk between the lignin and hemicellulose biosynthetic pathway, which seems not to be the case for cellulose.

The thinner the seed coat of *B. napus* cultivars, the larger the proportion occupied by the embryo, which leads to increasing contents of oil and protein (Slominski et al. 1999). The seed lignin content is confirmed to be significantly negatively correlated with seed oil content (Behnke et al. 2018, Miao et al. 2019). Similar results could be obtained in this study, where we found that oil content did show a significant difference between the low and high lignin groups. The low lignin group had a higher oil content. This was expected when compared to the parental genotypes and to the results of Behnke et al. (2018). Further, protein in defatted meal (%) and the sum of oil and protein content (%) showed a significant difference between the lignin groups, where the low lignin group contains significantly more. Similar results were observed in Behnke et al. (2018). However, the protein content did not show a significant difference between the groups, as in Behnke et al. (2018). Thus, reducing seed lignin content is associated with a reduction in seed coat content and increased oil content, while simultaneously improving the nutritive value of oilseed rape meal through a reduced fibre content.

### Candidate genes and ONT-technology

Genomic and transcriptomic data were used to unravel the genetic basis underlying the contrasting seed lignin content phenotypes between the parental genotypes SGDH14 and Express 617. In accordance with metabolic studies surveying the flavonoid – and general phenylpropanoid pathway in yellow and black seeded *B. napus*, 35 days after flowering (DAF) was chosen for the expression analysis, since those contents were significantly higher at 35 DAF in black seeds compared to yellow seeds (Wang et al. 2018). In this study, the low lignin and reduced fibre line SGDH14 lack the *PAL4, SEC8, MUCI21* and *SDP6* homologues located on C05 compared to the high lignin line Express 617 due to the HNRT event. *PAL4* is the most promising candidate gene for the lignin reduction, since its encoded protein catalyses the first rate limiting step in the general phenylpropanoid pathway leading to lignin synthesis (Fraser and Chapple 2011). Moreover, previous studies have proposed *PAL* as the key enzyme associated with lignin content in oilseed rape (Behnke et al. 2018, Ding et al. 2021, Wang et al. 2015). The expression of *PAL* was significantly higher in high lignin lines than in low lignin lines (Ding et al. 2021). The other candidate genes were related to fibre traits. SEC8 is involved in the localised deposition of seed coat pectin and formation of the central columella of seed epidermal cells (Kulich et al. 2010). *SEC8* and *PAL4* were also related to seed coat development in the study of Zhang et al. (2022). SDP6 is required for glycerol catabolism and is essential for post germinative growth and seedling establishment (Quettier et al. 2008). MUCI21 is necessary for the adherence of mucilage pectin to microfibrils made by Cellulose Synthase 5 (*CESA5*) to form an adherent mucilage layer on the seed coat (Ralet et al. 2016).

In complex allopolyploids like *B. napus*, pairing of homologous chromosomes occurring during meiosis is common and can lead to fragment exchanges between homoeologous chromosomes, resulting in widespread segmental deletions and duplications (Chalhoub et al. 2014). To clearly distinguish highly similar gene sequences and genomic rearrangements from another, long-read sequencing technology is valuable (Lee et al. 2020). In this study, the detection of the HNRT in SGDH14 was possible by long-reads derived from SGDH14 DNA and read mapping against the high quality long-read genome assembly of the winter *B. napus* cultivar Express 617 (Lee et al. 2020). Behnke et al. (2018) used Illumina 60K SNP markers and the Darmor-*bzh* genome assembly which had a higher level of fragmentation since it was constructed prior to the spread of long-read sequencing technologies. This did not allow the immediate detection of the HNRT in SGDH14. With the new long-read technology the sequence specificity of homoeologous chromosomes is enhanced, interference of repetitive sequences is reduced and HE breakpoints can be detected (Lee et al. 2020).

### *BnaPAL4* and paralogues

This study revealed *PAL4* as the most promising candidate gene, which is involved in the general phenylpropanoid pathway and therefore in the lignin synthesis. Since, *B. napus* and *A. thaliana* have multiple *PAL* paralogues (Fraser and Chapple 2011), their seed specificity needs to be determined. In *A. thaliana*, expression analysis showed that *AthPAL1, AthPAL2* and *AthPAL4* are expressed in stems of plants in later growth stages and *AthPAL2* and *AthPAL4* are expressed in seeds (Fraser and Chapple 2011, Raes et al. 2003). In this study, in total 14 *PAL* homologues including four *PAL1*, six *PAL2*, two *PAL3*, and two *PAL4* homologues across eight chromosomes were identified in the Express 617 genome assembly (Lee et al. 2020). The organ-specific expression in developing *B. napus* seeds showed that both *PAL4* homoeologues are much higher expressed than the other paralogues. Ding et al. (2021) found similar results in comparing the seed coat expression in high to low lignin cultivars and stated that *BnaPAL4* was significantly higher expressed in the high lignin lines than in the low lignin lines. Hong et al. (2017) found several DEGs involved in phenylpropanoid and flavonoid biosynthesis during seed development when comparing yellow and black seeded *B. napus* cultivars. The expression results show that *BnaPAL4* C05 was downregulated in low lignin lines as expected (Hong et al. 2017).

In *B. napus* cultivars, typically two *PAL4* homoeologues can be found on chromosomes C05 and A05. In this study, a HNRT in SGDH14 led to the deletion of *BnaPAL4* on C05 and its replacement by a copy of the A05 *BnaPAL4*. The two A05 *BnaPAL4* copies of SGDH14 revealed a ∼1.5 higher expression compared to the combined C05 and A05 *BnaPAL4* expression of Express 617. Thus, although SGDH14 lacks the C05 *PAL4* homoeologue, it still harbours two A05 *PAL4* copies which are expressed in seeds. But apparently these cannot complement SGDH14’s low lignin phenotype. Several hypotheses may explain the effect on lignin content of the two *BnaPAL4* A05 homologues in SGDH14. As stated previously, the C05 *BnaPAL4* and A05 *BnaPAL4* might be involved in different pathways. While the C05 *BnaPAL4* could be specific for the seed lignin biosynthesis, the A05 *BnaPAL4* could be primarily involved in other pathways, e.g. in PA biosynthesis. The analysis of seed coat layers spatio transcription patterns would enable to investigate expression of the two *BnaPAL4* A05 (and A05’’) homologues in SGDH14. They are expressed in the inner integument, endothelium, of the seed coat where PAs are synthesised (Moïse et al. 2005, Auger et al. 2010) while not being expressed in the palisade cells where lignin is synthesised (Jiang and Deyholos 2010, Cosgrove and Jarvis 2012), i.e. they might not contribute to lignin biosynthesis. We observed several SNPs in the putative promoter sequence between the A05 and C05 *BnaPAL4* homoeologues, which might affect the binding of different transcription factors or regulators. This could in turn determine different pathway specificities of the A05 and C05 *BnaPAL4* homoeologues. We propose that *BnaPAL4* on chromosome C05 is stronger associated with the lignin biosynthesis in the seed coat than its homoeologue on A05. Yin et al. (2022) found a similar divergence in function in accumulation of lignins, flavonoids and glucosinolates of two types of cinnamoyl-CoA reductase genes in *B. napus*. Finally, besides *PAL4*, additional candidate genes like *SEC8, SDP6*, and *MUCI21* might play a more important role in the control of seed lignin content than initially thought. However, clearly further functional genomic research is necessary to decipher the role of the identified candidate genes and especially of both *BnaPAL4* homoeologues in lignin biosynthesis in oilseed rape.

## Author Contributions

CM designed the experiment and developed the SGEF2 population. HMS and KH investigated and performed the experiments with subsequent data analysis. PV conducted library preparation and sequencing of ONT data. HMS performed bioinformatic analyses and data curation. HS and KH wrote the initial draft manuscript. HMS, KH, DH and CM revised the manuscript. All authors contributed to the article and approved the submitted version.

## Data Availability Statement

All data generated in this study can be found under the ENA/NCBI Bioproject ID PRJEB55241. The applied scripts in this study are freely available on GitHub: Major_low_ADL_QTL (DOI: 10.5281/zenodo.6970026).

## Acknowledgements

The authors would like to thank Katharina Ziesing, Jutta Schaper and Dietrich Kaufmann for excellent technical assistance. Moreover, we thank Boas Pucker for his bioinformatics support. We are extremely grateful to all researchers who submitted their *B. napus* RNA-Seq data sets to the appropriate databases and published their experimental findings.

