## Supplementary Information 1 for "Homoeologous non-reciprocal translocation explains a major QTL for seed lignin content in oilseed rape (*Brassica napus* L.)"

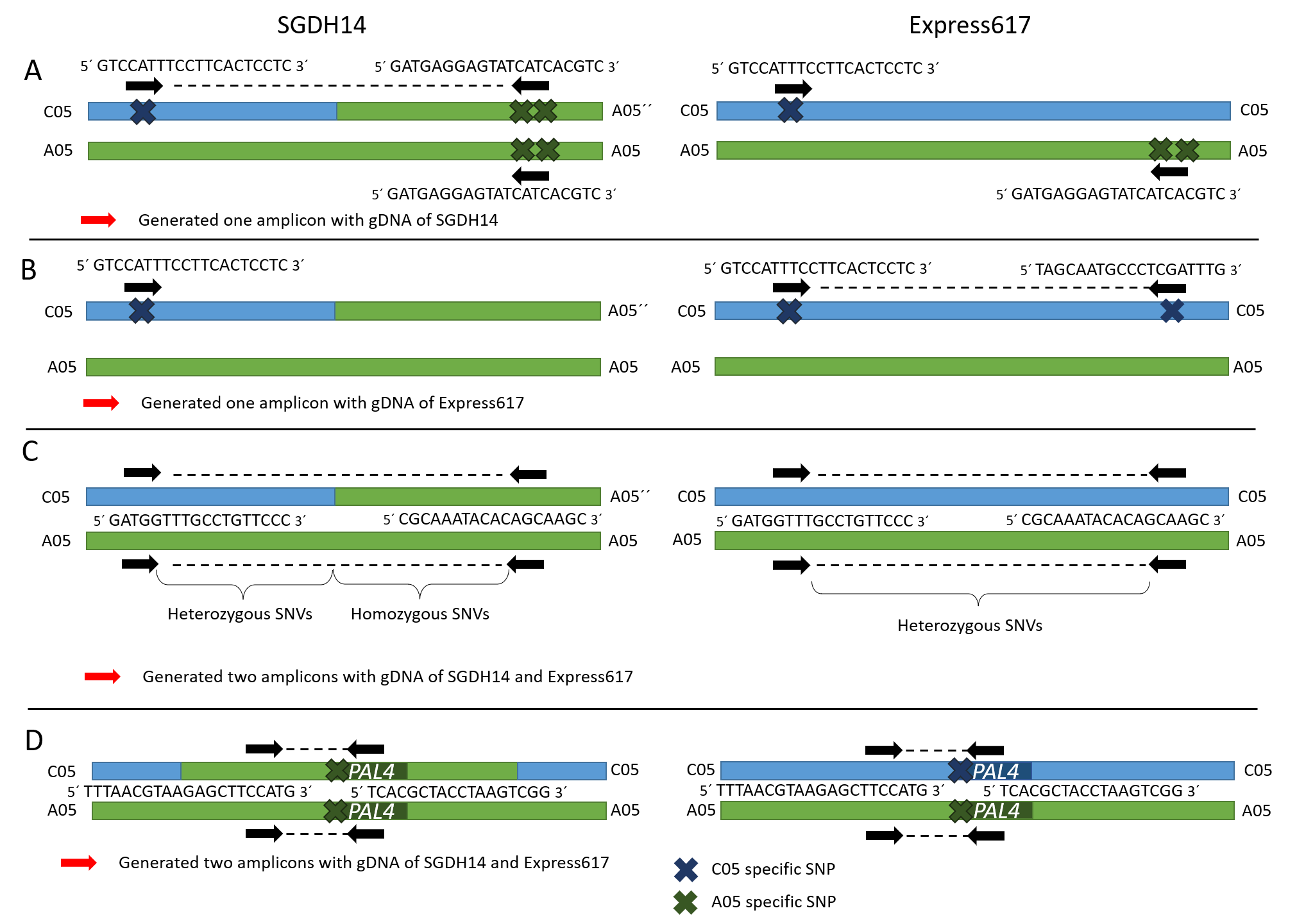

**Supplementary Figure 1: Oligonucleotide design strategies for validation of the HNRT border sequences.** (A) The subgenome-specific for SGDH14, (B) the subgenomes-specific for Express 617 and (C) not subgenome‑specific oligonucleotide design strategies used for the validation of the border sequences are shown, (D) subgenome-specific oligonucleotide design to determine if the *PAL4* gene of the A and/or the C chromosome is present. Oligonucleotides are marked as black arrows. Each strategy was applied for the left and right border, respectively.

**Supplementary Table 1: Oligonucleotides used in this study for the three different strategies and for the amplification of *PAL4*.** PCR products were sequenced to confirm the correct gene was amplified. The oligonucleotides written in bold and italics indicate the positions which are different between the subgenome-specific primers on the A and C chromosome.

| **Strategy** | **Oligonucleotide name** | **Strand** | **Sequence (5’ to 3’)** | **Annealing Temp. [°C]** |
| --- | --- | --- | --- | --- |
| A – left border | SGDH14_spec_fw_L | forward | GTC***CA***TTTCCTTCACTCCTC | 60 |
| A – left border | SGDH14_spec_rev_L | reverse | GATGAGGAGTATC***AT***CACGTC |  |
| A – right border | SGDH14_spec_fw_R | forward | TCAGA***C***GGC***AG***CGTTTAC | 60 |
| A – right border | SGDH14_spec_rev_R | reverse | TTGCCACCACCACC***T***AC |  |
| B – left border | Exp_spec_fw_L | forward | GTC***CA***TTTCCTTCACTCCTC | 60 |
| B – left border | Exp_spec_rev_L | reverse | TAGCAATGCCCTC***G***AT***T***TG |  |
| B – right border | Exp_spec_fw_R | forward | TGGTCAGATG***G***C***TC***CGTTTAC | 60 |
| B – right border | Exp_spec_rev_R | reverse | TTGCCACCACCACC***T***AC |  |
| C – left border | Not_spec_fw_L | forward | GATGGTTTGCCTGTTCCC | 56 |
| C – left border | Not_spec_rev_L | reverse | TCGCTGAATAGTCGCAAG |  |
| C – right border | Not_spec_fw_R | forward | AACCAAATCCGTTGATGC | 56 |
| C – right border | Not_spec_rev_R | reverse | TTGCGTGACTGCTCCAAG |  |
| D – PAL4 | PAL_nspec_fw_1 | forward | TTTAACGTAAGAGCTTCCATG | 55 |
| D – PAL4 | PAL_nspec_rev_2 | reverse | TCACGCTACCTAAGTCGG |  |

**Supplementary Table 2: PCR program for the different subgenomes-specific strategies (A, B) and not subgenome‑specific strategy (C), as well as for the amplification of the *PAL4* gene (D).**

| **Steps** | **Temp. [°C]** | **Time [sec]** | **No. of Cycles** |
| --- | --- | --- | --- |
| PCR program for strategy A and C; PAL4 | | | |
| initial denaturation | 95 | 60 | 1x |
| denaturation | 95 | 15 |  |
| annealing | 54, 56, 58, 60 | 15 | 35x |
| elongation | 72 | 30 |  |
| final elongation | 72 | 180 | 1x |
| hold | 8 | infinite |  |
| PCR program for strategy B | | | |
| **Steps** | **Temp. [°C]** | **Time [sec]** | **No. of Cycles** |
| initial denaturation | 95 | 180 | 1x |
| denaturation | 95 | 30 |  |
| annealing | 70 _ 0.7°C decrease each cycle | 45 | 15x |
| elongation | 72 | 45 |  |
| denaturation | 95 | 30 |  |
| annealing | 60 or 61 | 45 | 25x |
| elongation | 72 | 45 |  |
| elongate | 72 | 300 | 1x |
| halt reaction | 4 | 300 | 1x |
| hold | 20 | infinite | 1x |
