## Supplementary Information 2 for "Homoeologous non-reciprocal translocation explains a major QTL for seed lignin content in oilseed rape (*Brassica napus* L.)"

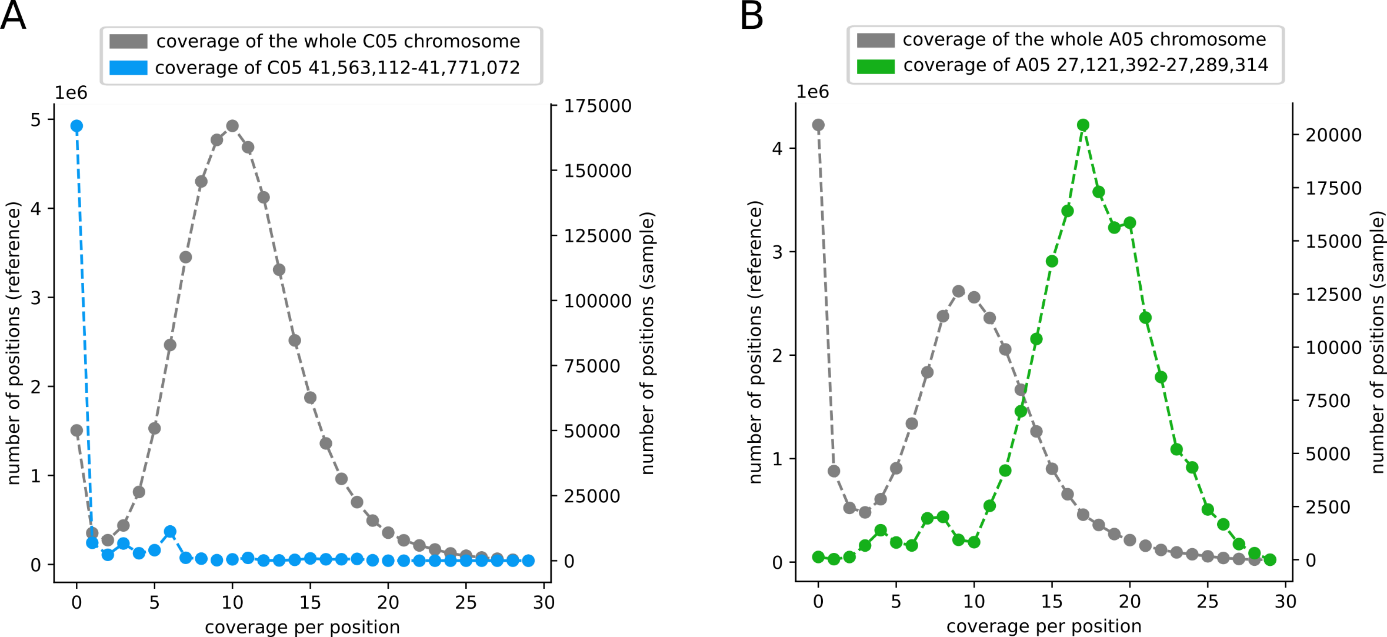

**Supplementary Figure 2: Coverage analysis of the HNRT region of SGDH14 located within the major low lignin QTL.** (A) The coverage per position of the region of interest, namely C05 ranging from 41,563,112 to 41,771,072 (sample), compared to the coverage of the whole C05 chromosome (reference) is shown based on the mapping of SGDH14 derived long-reads against the Express 617 assembly. The average coverage (median) of the whole chromosome is 10, while those of the region of interest is 0. (B) The coverage per position of the region of interest, namely A05 ranging from 27,121,280 to 27,289,335 (sample), compared to the coverage of the whole A05 chromosome (reference) is shown based on the mapping of SGDH14 derived long-reads against the Express 617 assembly. The average coverage (median) of the whole chromosome is 9, while those of the region of interest is 17.

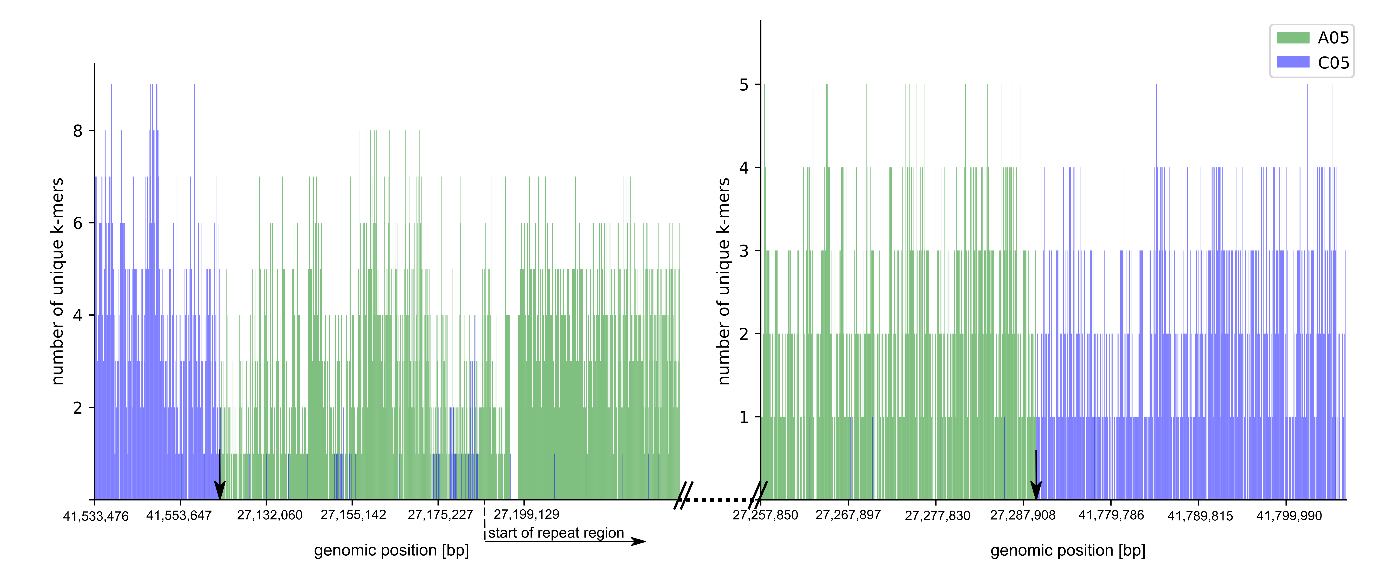

**Supplementary Figure 3: K-mer analysis elucidates chimeric border flanking long-reads of SGDH14.** (Left) Exemplary left border flanking read divided into A05 specific and C05 specific k-mers, which are marked in green and blue respectively. The genomic position based on the Express 617 assembly is given in bp and the number of identified subgenomes specific k-mers is given on the y-axis. The position of the predicted left genomic border is marked with an black arrow. The last base of the last C05 specific k-mer is at 41,563,009 while the first base of the first A05 specific k-mer is at 27,121,379. A dashed line marks the approximate start of the repeat region, while the backslashes and the dotted line indicate that the repeat region is extended beyond the read’s end. (Right) Exemplary right border flanking read divided into A05 specific and C05 specific k-mers. The position of the predicted right genomic border is marked with an arrow. The last base of last A05 specific k-mer is at 27,289,319, while the first base of the first C05 specific k-mer is at 41,771,107.
