## Supplementary Information 4 for "Homoeologous non-reciprocal translocation explains a major QTL for seed lignin content in oilseed rape (*Brassica napus* L.)"

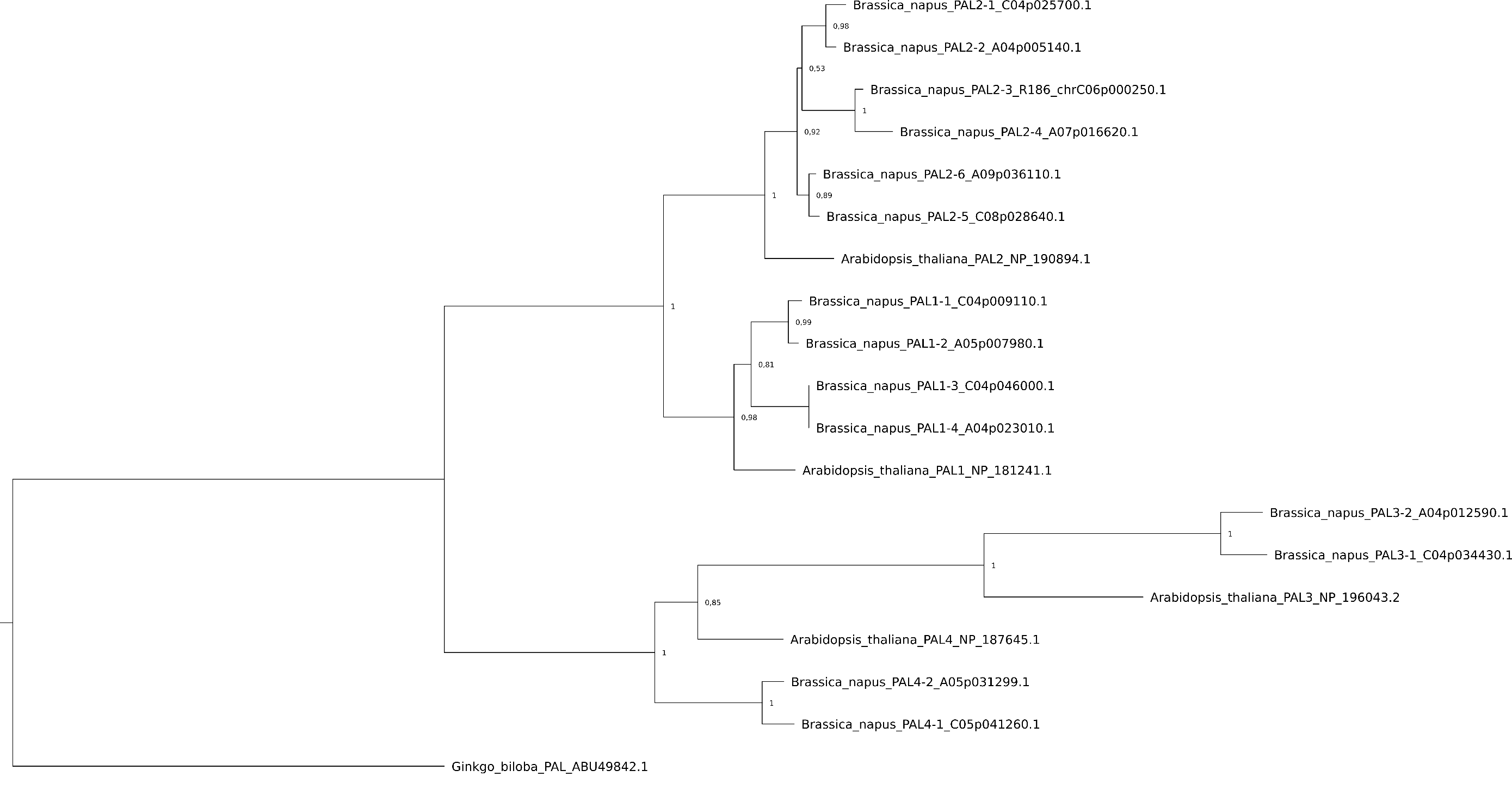

**Supplementary Figure 4: Phylogeny of PAL candidates of *B. napus* Express 617 and previously described PAL sequences.** The maximum‑likelihood tree was constructed with FastTree using 10,000 bootstraps and relative bootstrap-values are shown next to relevant nodes. The phylogenetic tree is based on amino acid sequences. The *Ginkgo biloba* PAL sequence was used to root the tree. The GenBank ID of the respective sequence or the Express 617 gene ID is given for each sequence.

**Supplementary Table 6: Amino acid and coding sequences of *B. napus* Express 617 PALs.**

>BnaPAL1-4_A04p023010.1

MEVNGLSHGGEVDAMLCGGEIKKNATVVGADPLNWGAAAEQMKGSHLDEVKRMVKEFRRPVVNLGGETLTIGQVAAISTLGNGVKVELSETARAGVKASSDWVMESMGKGTDSYGVTTGFGATSHRRTKNGVALQKELIRFLNAGIFGSTKETSHTLPHSATRAAMLVRINTLLQGYSGIRFEILEAITSFLNNNITPSLPLRGTITASGDLVPLSYIAGLLTGRPNSKATGPNGEALNAEEAFKMAGVTSGFFDLQPKEGLALVNGTAVGSGMASMVLFEANVLSVLAEVLSAVFAEVMSGKPEFTDHLTHRLKHHPGQIEAAAIMEHILDGSSYMKLAQKLHEMDPLQKPKQDRYALRTSPQWLGPQIEVIRYATKSIEREINSVNDNPLIDVSRNKAIHGGNFQGTPIGVSMDNTRLAIAAIGKLMFAQFSELVNDYYNNGLPSNLTASRNPSLDYGFKGAEIAMASYCSELQYLANPVTSHVQSAEQHNQDVNSLGLISSRKTSEAVDILKLMSTTFLVGICQAIDLRHLEENLKQTVKNTVSQVAKKVLTTGVNGELHPSRFCEKDLLKVVDREQVYTYADDPCSATYPLIQKLRQVIVDHALVNGESEKNAMTSIFHKIGAFEEELKAVLPKEVEAARAAYDNGTAAIPNRIKECRSYPLYRFVREELGTELLTGEKATSPGEEFDKVFTAICEGKIIDPLMECLDEWNGAPIPIC*

>BnaPAL1-4_A04p023010.1_BnaEXP

ATGGAGGTTAACGGATTATCACACGGAGGAGAAGTCGACGCTATGTTATGCGGCGGAGAGATCAAGAAGAATGCCACGGTGGTTGGGGCTGATCCTCTCAACTGGGGAGCTGCAGCGGAGCAAATGAAAGGGAGCCATTTGGATGAGGTGAAGAGAATGGTGAAGGAGTTTAGGAGGCCAGTGGTGAATCTCGGAGGCGAGACTCTCACGATCGGACAAGTGGCAGCAATCTCGACCCTTGGAAACGGTGTGAAGGTGGAGTTGTCGGAGACGGCGAGAGCCGGTGTGAAAGCGAGTAGTGATTGGGTGATGGAGAGTATGGGCAAAGGAACCGATAGTTATGGTGTTACTACTGGTTTTGGTGCTACTTCTCACCGGAGAACCAAGAACGGTGTCGCTCTTCAGAAAGAGCTTATAAGATTTCTGAACGCCGGAATATTCGGCAGCACGAAGGAAACATCCCACACATTACCACACTCCGCCACAAGAGCCGCCATGCTCGTACGCATCAACACTCTCCTCCAAGGCTACTCAGGCATACGTTTCGAGATCCTCGAGGCCATAACAAGTTTCCTCAACAACAACATCACTCCTTCACTCCCTCTCCGCGGCACCATCACCGCCTCCGGAGACCTCGTCCCCCTCTCCTACATCGCCGGACTCTTAACCGGACGTCCCAACTCCAAAGCCACCGGTCCCAACGGCGAAGCCTTGAACGCAGAGGAAGCCTTCAAAATGGCAGGAGTCACCTCCGGCTTCTTCGACCTCCAGCCTAAAGAAGGCCTCGCCCTAGTCAACGGCACGGCCGTTGGCTCCGGCATGGCGTCGATGGTTCTCTTCGAAGCCAACGTCCTCTCGGTCCTGGCCGAGGTTTTATCAGCCGTCTTCGCCGAGGTGATGAGCGGGAAGCCGGAGTTCACCGATCATTTGACTCACCGGCTGAAGCACCACCCCGGGCAGATAGAAGCCGCGGCTATAATGGAACACATTCTCGACGGGAGCTCGTACATGAAGCTAGCTCAGAAGCTTCACGAGATGGATCCGTTACAAAAACCCAAACAAGACCGTTACGCTCTACGTACCTCTCCCCAATGGCTAGGTCCTCAGATAGAAGTGATCCGTTACGCGACCAAGTCCATCGAGCGTGAGATAAACTCCGTCAACGACAACCCTTTGATCGATGTCTCGAGAAACAAAGCCATTCACGGTGGTAACTTCCAGGGAACACCCATCGGTGTGTCTATGGACAACACACGTTTAGCTATCGCGGCCATAGGGAAGCTCATGTTCGCTCAGTTCTCCGAGCTGGTCAACGATTATTACAACAACGGTCTTCCTTCGAACCTAACCGCTTCGAGGAACCCTAGTTTGGACTACGGTTTCAAAGGAGCGGAGATCGCGATGGCTTCGTACTGTTCAGAGCTTCAGTACTTGGCCAACCCCGTGACTAGCCATGTTCAATCAGCAGAGCAACACAACCAAGACGTGAACTCTTTAGGACTAATCTCATCTCGCAAGACATCAGAAGCTGTAGACATACTCAAACTCATGTCGACAACGTTCCTTGTCGGAATCTGCCAAGCCATCGATCTAAGACACTTGGAGGAGAATCTAAAACAAACGGTCAAGAACACAGTCTCTCAAGTCGCCAAGAAAGTTCTCACCACTGGAGTCAACGGTGAGCTTCACCCATCACGCTTCTGCGAGAAAGACTTACTCAAAGTCGTCGACCGTGAGCAAGTCTACACGTACGCGGACGACCCATGCAGCGCCACGTACCCGTTGATCCAGAAGCTGAGGCAAGTCATCGTTGACCATGCTTTGGTCAACGGTGAGAGCGAGAAGAATGCAATGACTTCGATCTTCCATAAGATTGGAGCCTTCGAGGAGGAGCTCAAGGCGGTGCTACCTAAAGAAGTGGAGGCAGCTAGAGCGGCGTATGATAACGGAACAGCTGCGATACCGAACAGGATCAAGGAGTGTAGGTCGTATCCTTTGTATAGGTTTGTGAGGGAGGAGCTTGGAACAGAGCTTTTGACCGGAGAGAAAGCGACGTCGCCCGGAGAAGAGTTTGATAAGGTTTTCACGGCGATTTGTGAAGGGAAGATTATTGATCCGTTGATGGAGTGTCTCGATGAGTGGAACGGAGCTCCCATTCCTATATGTTAA

>BnaPAL1-3_C04p046000.1

MEVNGLSHGGEVDAMLCGGEIKKNATVVGADPLNWGAAAEQMKGSHLDEVKRMVKEFRRPVVNLGGETLTIGQVAAISTLGNGVKVELSETARAGVKASSDWVMESMGKGTDSYGVTTGFGATSHRRTKNGVALQKELIRFLNAGIFGSTKETSHTLPHSATRAAMLVRINTLLQGYSGIRFEILEAITSFLNNNITPSLPLRGTITASGDLVPLSYIAGLLTGRPNSKATGPNGEALNAEEAFKMAGVTSGFFDLQPKEGLALVNGTAVGSGMASMVLFEANVLSVLAEVLSAVFAEVMSGKPEFTDHLTHRLKHHPGQIEAAAIMEHILDGSSYMKLAQKLHEMDPLQKPKQDRYALRTSPQWLGPQIEVIRYATKSIEREINSVNDNPLIDVSRNKAIHGGNFQGTPIGVSMDNTRLAIAAIGKLMFAQFSELVNDYYNNGLPSNLTASRNPSLDYGFKGAEIAMASYCSELQYLANPVTSHVQSAEQHNQDVNSLGLISSRKTSEAVDILKLMSTTFLVGICQAIDLRHLEENLKQTVKNTVSQVAKKVLTTGVNGELHPSRFCEKDLLKVVDREQVYTYADDPCSATYPLIQKLRQVIVDHALVNGESEKNAMTSIFHKIGAFEEELKAVLPKEVEAARAAYDNGTAAIPNRIKECRSYPLYRFVREELGTELLTGEKATSPGEEFDKVFTAICEGKIIDPLMECLDEWNGAPIPIC*

>BnaPAL1-3_C04p046000.1_BnaEXP

ATGGAGGTTAACGGATTATCACACGGAGGAGAAGTCGACGCTATGTTATGCGGCGGAGAGATCAAGAAGAATGCCACGGTGGTTGGGGCTGATCCTCTCAACTGGGGAGCTGCAGCGGAGCAAATGAAAGGGAGCCATTTGGATGAGGTGAAGAGAATGGTGAAGGAGTTTAGGAGGCCAGTGGTGAATCTCGGAGGCGAGACTCTCACGATCGGACAAGTGGCAGCAATCTCGACCCTTGGAAACGGTGTGAAGGTGGAGTTGTCGGAGACGGCGAGAGCCGGTGTGAAAGCGAGTAGTGATTGGGTGATGGAGAGTATGGGCAAAGGAACCGATAGTTATGGTGTTACTACTGGTTTTGGTGCTACTTCTCACCGGAGAACCAAGAACGGTGTCGCTCTTCAGAAAGAGCTTATAAGATTTCTGAACGCCGGAATATTCGGCAGCACGAAGGAAACATCCCACACATTACCACACTCCGCCACAAGAGCCGCCATGCTCGTACGTATCAACACTCTCCTCCAAGGATACTCCGGCATACGTTTCGAGATCCTCGAGGCCATAACAAGTTTCCTCAACAACAACATCACTCCTTCCCTCCCTCTCCGCGGCACCATCACCGCCTCCGGAGACCTCGTCCCCCTCTCCTACATCGCCGGACTCTTAACCGGACGTCCCAACTCCAAAGCTACCGGTCCCAACGGCGAAGCCCTAAACGCAGAGGAAGCCTTCAAAATGGCTGGAGTCACCTCAGGCTTCTTTGACCTCCAGCCTAAGGAAGGCCTCGCCCTAGTCAACGGCACGGCCGTTGGCTCCGGCATGGCGTCGATGGTTCTCTTCGAAGCCAACGTCCTCTCGGTCCTGGCCGAGGTTTTATCAGCCGTCTTCGCCGAGGTGATGAGCGGGAAGCCGGAGTTCACCGATCATTTGACTCACCGGCTGAAGCACCACCCCGGGCAGATAGAAGCCGCGGCTATAATGGAACACATTCTCGACGGGAGCTCGTACATGAAGCTAGCTCAGAAGCTTCACGAGATGGATCCGTTACAAAAACCCAAACAAGACCGTTACGCTCTACGTACCTCTCCCCAATGGCTAGGTCCTCAGATAGAAGTGATCCGTTACGCGACCAAGTCCATCGAGCGTGAGATAAACTCCGTCAACGACAACCCTTTGATCGATGTCTCGAGAAACAAAGCCATTCACGGTGGTAACTTCCAGGGAACACCCATCGGTGTGTCTATGGACAACACACGTTTAGCTATCGCGGCCATAGGGAAGCTCATGTTCGCTCAGTTCTCCGAGCTGGTCAACGATTATTACAACAACGGTCTTCCTTCGAACCTAACCGCTTCGAGGAACCCTAGTTTGGACTACGGTTTCAAAGGAGCGGAGATCGCGATGGCTTCGTACTGTTCAGAGCTTCAGTACTTGGCCAACCCAGTGACTAGCCATGTTCAATCAGCTGAGCAACACAACCAAGACGTGAACTCTTTAGGACTAATCTCGTCTCGCAAGACATCAGAAGCTGTAGACATACTCAAACTCATGTCTACAACGTTCCTTGTCGGAATCTGCCAAGCCATCGATCTAAGACACTTGGAGGAGAATCTAAAACAAACGGTCAAGAACACAGTCTCTCAAGTCGCCAAGAAAGTTCTCACCACTGGAGTCAACGGTGAGCTTCACCCATCACGCTTCTGCGAGAAAGACTTACTCAAAGTCGTCGACCGTGAGCAAGTCTACACGTACGCGGACGACCCATGCAGCGCCACGTACCCGTTGATCCAGAAGCTGAGGCAAGTCATCGTTGACCATGCTTTGGTCAACGGTGAGAGCGAGAAGAATGCAATGACTTCGATCTTCCATAAGATTGGAGCCTTCGAGGAGGAGCTCAAGGCGGTGCTTCCGAAAGAAGTGGAGGCAGCTAGAGCGGCGTATGATAACGGAACAGCTGCGATACCGAACAGGATCAAGGAGTGTAGGTCGTATCCTTTGTATAGGTTTGTGAGGGAGGAGCTTGGAACAGAGCTTTTGACCGGAGAGAAAGCGACGTCGCCCGGAGAAGAGTTTGATAAGGTTTTCACGGCGATCTGTGAAGGGAAGATTATTGATCCGTTGATGGAGTGTCTCGATGAGTGGAACGGAGCTCCCATTCCTATATGTTAA

>BnaPAL1-2_A05p007980.1

MEINGSSHKVDAMLCGAEIKNVTVAAADPLNWGAAAEQMKGSHLDEVKRMVMEFRKPVVNLGGETLTIGQVAAISTVGNGVKVELSETARAGVKASSDWVMESMGKGTDSYGVTTGFGATSHRRTKNGVALQKELIRFLNAGIFGSTKETCHTLPHSATRAAMLVRINTLLQGYSGIRFEILEAITSFLNTNITPSLPLRGTITASGDLVPLSYIAGLLTGRPNSKATGPNGEALNAEEAFKKAGIPSGFFDLQPKEGLALVNGTAVGSGMASMVLFETNVLSVLAEVLSAVFAEVMSGKPEFTDHLTHRLKHHPGQIEAAAIMEHILDGSSYMKLAKKLHELDPLQKPKQDRYALRTSPQWLGPQIEVIRYATKSIEREINSVNDNPLIDVSRNKAIHGGNFQGTPIGVSMDNTRLAIASIGKLMFAQFSELVNDFYNNGLPSNLTASRNPSLDYGFKGAEIAMASYCSELQYLANPVTTHVQSAEQHNQDVNSLGLISSRKTAEAVDILKLMSTTFLVAICQAVDLRHLEENLKQTVKNTVSQVAKKVLTTGVNGELHPSRFCEKDLLKVVDREQVYTYADDPCSATYPLIQKLRQVIVDHALVNGESEKNAMTSIFHKIGAFEEELKAVLPDEVEAARVAYDNGTSAIPNRIKECRSYPLYRFVREELGTELLTGEKVTSPGEEFDKVFTAICEGKIIDPLMECLSEWNGAPIPIC*

>BnaPAL1-2_A05p007980.1_BnaEXP

ATGGAGATTAACGGATCATCACACAAAGTCGACGCTATGTTATGCGGCGCAGAGATCAAGAATGTCACGGTGGCTGCGGCGGATCCTCTCAACTGGGGGGCTGCGGCGGAGCAAATGAAAGGGAGCCATTTGGATGAAGTGAAGAGAATGGTTATGGAGTTTAGGAAGCCTGTGGTGAATCTCGGAGGAGAGACTCTGACGATCGGACAAGTAGCAGCGATCTCGACCGTTGGAAATGGTGTGAAGGTGGAGCTGTCGGAGACGGCAAGAGCCGGTGTGAAGGCGAGCAGTGATTGGGTGATGGAGAGTATGGGCAAAGGCACTGATAGTTATGGTGTTACTACTGGTTTTGGTGCTACTTCTCATCGTAGAACCAAAAATGGCGTCGCACTTCAGAAGGAGCTTATAAGATTCCTGAACGCCGGAATATTCGGCAGCACGAAAGAAACATGCCACACACTACCACACTCCGCCACGAGAGCCGCCATGCTCGTACGCATCAACACTCTCCTCCAAGGATACTCCGGCATCCGCTTCGAGATCCTCGAGGCCATAACCAGCTTCCTCAACACCAACATCACTCCTTCCCTCCCTCTCCGCGGCACCATCACCGCCTCCGGAGACCTCGTCCCCCTCTCCTACATCGCCGGCCTCCTCACCGGCCGTCCCAACTCCAAAGCCACCGGTCCCAACGGCGAAGCCTTGAACGCAGAGGAAGCCTTCAAGAAGGCTGGAATCCCCTCCGGATTCTTTGACTTGCAGCCCAAGGAAGGTCTCGCCCTAGTCAACGGCACAGCCGTTGGCTCCGGCATGGCCTCGATGGTGCTCTTCGAGACGAACGTCCTCTCGGTCCTGGCCGAGGTTCTCTCAGCTGTCTTCGCCGAGGTGATGAGCGGAAAGCCGGAGTTCACCGACCATTTAACCCACAGGCTTAAACACCACCCCGGCCAGATCGAAGCCGCGGCGATAATGGAGCACATCCTTGACGGGAGCTCGTACATGAAGCTAGCTAAGAAGCTACACGAGTTGGATCCGTTACAGAAACCTAAACAAGACCGTTACGCGCTCCGCACGTCACCTCAATGGCTAGGTCCTCAGATAGAAGTGATCCGTTACGCAACAAAGTCCATCGAACGTGAGATCAACTCCGTCAACGACAACCCCTTGATCGACGTTTCGAGAAACAAGGCCATTCACGGAGGTAACTTCCAGGGAACACCGATAGGTGTCTCCATGGACAACACGCGTCTCGCTATAGCATCGATAGGGAAGCTCATGTTCGCTCAGTTCTCTGAGCTCGTCAACGACTTCTACAACAACGGTCTTCCTTCTAACCTAACCGCCTCGAGGAACCCTAGTTTGGACTACGGTTTCAAAGGAGCGGAGATCGCTATGGCATCCTACTGTTCCGAGCTTCAGTACTTAGCCAATCCAGTTACAACCCATGTTCAATCCGCGGAGCAACACAACCAAGACGTGAACTCTTTAGGACTAATCTCGTCTCGCAAGACGGCAGAAGCTGTTGACATTCTCAAACTCATGTCGACGACGTTCCTTGTCGCAATCTGCCAAGCTGTAGATCTAAGACACTTGGAGGAGAATCTAAAACAAACCGTGAAGAACACGGTCTCTCAAGTGGCTAAGAAAGTTCTAACCACTGGAGTCAACGGTGAGCTTCACCCATCTCGCTTCTGCGAGAAAGATTTACTCAAAGTCGTTGACCGTGAACAAGTCTACACGTACGCGGACGATCCATGCAGCGCTACGTACCCGTTGATCCAGAAGCTAAGGCAAGTTATCGTTGACCATGCTTTGGTCAACGGTGAGAGTGAGAAGAACGCTATGACTTCGATCTTCCACAAGATTGGTGCTTTCGAGGAGGAGCTCAAGGCGGTGCTTCCTGATGAAGTGGAGGCAGCTAGAGTGGCGTACGATAACGGGACATCTGCTATACCGAACCGGATCAAGGAGTGTAGGTCGTATCCTTTGTATAGGTTCGTGAGGGAGGAGCTTGGAACTGAGCTTTTGACCGGAGAGAAAGTCACGTCGCCGGGAGAAGAGTTTGATAAGGTTTTCACGGCGATATGTGAAGGGAAGATCATTGATCCGTTGATGGAGTGTCTCAGTGAGTGGAACGGAGCTCCCATTCCAATCTGCTAA

>BnaPAL1-1_C04p009110.1

MEINGSSYKVDAMLCGGETKNVTVAAADPLNWGAAADQMKGSHLDEVKRMVMEFRKPVVNLGGETLTIGQVAAISTVGNGVKVELSETARAGVKASSDWVMESMGKGTDSYGVTTGFGATSHRRTKNGVALQKELIRFLNAGIFGSTKETCHTLPHSATRAAMLVRINTLLQGYSGIRFEILEAITSFLNTNITPSLPLRGTITASGDLVPLSYIAGLLTGRPNSKATGPNGEALNAEEAFKKAGIPSGFFDLQPKEGLALVNGTAVGSGMASMVLFETNVLSVLAEVLSAVFAEVMSGKPEFTDHLTHRLKHHPGQIEAAAIMEHILDGSSYMKLAQKLHEMDPLQKPKQDRYALRTSPQWLGPQIEVIRYATKSIEREINSVNDNPLIDVSRNKAIHGGNFQGTPIGVSMDNTRLAIASIGKLMFAQFSELVNDFYNNGLPSNLTASRNPSLDYGFKGAEIAMASYCSELQYLANPVTTHVQSAEQHNQDVNSLGLISSRKTAEAVDILKLMSTTFLVAICQAVDLRHLEENLKQTVKNTVSQVAKKVLTTGDNGELHPSRFCEKDLLKVVDREQVYTYADDPCSATYPLIQKLRQVIVDHALVNGESEKNAMTSIFHKIGAFEEELKAVLPDEVEAARVAYDNGTSAIPNRIKECRSYPLYRFVREELGTELLTGEKVTSPGEEFDKVFTAICEGKIIDPLMECLSEWNGAPIPIC*

>BnaPAL1-1_C04p009110.1_BnaEXP

ATGGAGATTAACGGATCATCATACAAAGTCGACGCTATGTTATGCGGCGGAGAGACCAAGAATGTCACGGTGGCTGCGGCGGATCCTCTGAACTGGGGAGCTGCGGCGGATCAAATGAAAGGGAGCCATTTGGATGAAGTGAAGAGAATGGTTATGGAGTTTAGGAAGCCTGTGGTGAATCTCGGAGGAGAGACTCTGACGATCGGACAAGTAGCAGCGATCTCGACCGTTGGAAATGGTGTGAAGGTGGAGCTGTCGGAGACGGCAAGAGCCGGTGTGAAGGCGAGCAGTGATTGGGTGATGGAGAGTATGGGCAAAGGCACTGATAGTTATGGTGTTACTACTGGTTTTGGTGCTACTTCTCATCGTAGAACCAAAAATGGCGTCGCACTTCAGAAGGAGCTTATAAGATTCCTGAACGCCGGAATATTCGGCAGCACGAAAGAAACATGCCACACACTGCCACACTCCGCCACAAGAGCTGCCATGCTCGTACGCATAAACACTCTCCTCCAAGGATACTCCGGCATCCGCTTCGAGATCCTCGAGGCCATAACCAGCTTCCTCAACACCAACATCACTCCTTCCCTCCCTCTCCGCGGCACCATCACCGCCTCCGGAGACCTCGTCCCCCTCTCCTACATCGCCGGCCTCCTCACCGGCCGTCCCAACTCCAAAGCCACCGGTCCCAACGGCGAAGCCTTGAACGCAGAGGAAGCCTTCAAGAAGGCCGGAATCCCCTCTGGATTCTTTGACTTGCAGCCCAAGGAAGGTCTCGCCCTAGTCAACGGCACAGCCGTTGGCTCCGGCATGGCCTCGATGGTTCTCTTCGAGACAAACGTCCTGTCGGTTCTGGCCGAGGTTCTCTCAGCTGTCTTCGCCGAGGTGATGAGCGGCAAGCCGGAGTTCACCGACCATTTAACCCACAGGCTGAAGCACCACCCCGGCCAGATCGAAGCCGCGGCGATAATGGAGCACATCCTCGACGGGAGCTCGTACATGAAGCTAGCTCAGAAGCTACACGAGATGGATCCGTTACAGAAACCTAAACAAGACCGTTACGCGCTCCGCACATCACCTCAATGGCTAGGTCCTCAGATAGAAGTGATCCGTTACGCCACCAAGTCCATCGAGCGTGAGATAAACTCCGTCAACGACAACCCTTTGATCGACGTTTCAAGAAACAAAGCGATTCACGGAGGTAACTTCCAGGGGACACCCATCGGTGTCTCCATGGACAACACGCGTCTCGCTATAGCATCGATAGGGAAGCTCATGTTCGCTCAGTTCTCCGAGCTTGTTAACGACTTCTACAACAACGGTCTTCCTTCTAACCTAACCGCTTCGAGGAACCCTAGTTTGGACTACGGTTTCAAAGGAGCGGAGATCGCTATGGCTTCCTACTGTTCAGAGCTTCAGTACTTGGCCAATCCAGTTACAACCCATGTTCAGTCAGCAGAGCAACACAACCAAGACGTGAACTCTTTAGGACTAATCTCATCACGCAAGACGGCAGAAGCTGTCGACATACTCAAACTCATGTCGACAACGTTCCTCGTCGCAATCTGCCAAGCTGTGGATCTAAGACACTTGGAGGAGAATCTAAAACAAACCGTGAAGAACACTGTCTCTCAAGTGGCCAAGAAAGTTCTAACCACTGGAGACAACGGTGAGCTTCACCCATCTCGCTTCTGCGAGAAAGATTTACTCAAAGTCGTTGACCGTGAACAAGTCTACACGTACGCGGACGATCCATGCAGCGCCACGTACCCGTTGATCCAGAAGCTAAGGCAAGTTATCGTTGACCATGCTTTGGTCAACGGTGAGAGCGAGAAGAACGCTATGACTTCGATCTTCCACAAGATTGGTGCTTTCGAGGAGGAGCTCAAGGCGGTGCTTCCTGACGAAGTGGAGGCAGCTCGAGTGGCGTATGATAACGGGACATCTGCTATACCGAACCGGATCAAGGAGTGTAGGTCGTATCCTTTGTATAGGTTCGTGAGGGAGGAGCTTGGAACTGAGCTTTTGACCGGAGAGAAGGTGACGTCGCCGGGAGAAGAGTTTGATAAGGTTTTCACGGCGATATGTGAAGGGAAGATCATTGATCCGTTGATGGAGTGTCTCAGTGAGTGGAACGGAGCTCCCATTCCAATATGCTAA

>BnaPAL2-2_A04p005140.1

MDQINGSVHQNGKTEAMLLCGGLEKTKVTVAADPLNWGAAAEQMKGSHLDEVKRMVEDYRKPVVNLGGETLTIGQVAAISNVGGGVKVELAEASRAGVKASSDWVMESMGKGTDSYGVTTGFGATSHRRTKNGTALQTELIRFLNAGIFGNTKETCHTLPESATRAAMLVRVNTLLQGYSGIRFEILEAITSLLNHNISPSLPLRGTITASGDLVPLSYIAGLLTGRPNSKATGPNGESLTAEEAFKKAGITSGFFDLQPKEGLALVNGTAVGSGMASMVLFEANVQSVLAEVLSAIFAEVMSGKPEFTDHLTHRLKHHPGQIEAAAIMEHILDGSSYMKLAAKLHEMDPLQKPKQDRYALRTSPQWLGPQIEVIRHATKSIEREINSVNDNPLIDVSRNKAIHGGNFQGTPIGVSMDNTRLAVAAIGKLMFAQFSELVNDFYNNGLPSNLTASNNPSLDYGFKGAEIAMASYCSELQYLANPVTTHVQSAEQHNQDVNSLGLISSRKTSEAVDILKLMSTTFLVAICQAVDLRHLEENLRQTVKNTVSQVAKKVLTTGVNGELHPSRFCEKDLLKVVDREQVFTYVDDPCSATYPLMQKLRQVIVDHALSNGETEKNAVTSIFQKIGAFEEELKMVLPKEVDATREAYGNGTAAIPNRIKECRSYPLYKFVREELGTKLLTGEKVVSPGEEFDKVFTAMCEGKIIDPLMDCLKEWNGAPIPIC*

>BnaPAL2-2_A04p005140.1_BnaEXP

ATGGATCAGATCAACGGATCAGTTCACCAAAACGGCAAAACCGAAGCCATGTTGTTGTGCGGCGGACTAGAGAAGACTAAAGTGACGGTGGCGGCGGATCCGTTGAATTGGGGTGCTGCGGCGGAGCAGATGAAAGGGAGTCATTTGGATGAGGTGAAGAGGATGGTTGAGGATTACCGTAAACCGGTGGTGAATCTCGGCGGAGAAACACTGACGATCGGACAAGTCGCTGCGATCTCGAACGTAGGCGGTGGCGTTAAGGTTGAGCTAGCGGAGGCTTCAAGAGCCGGCGTGAAAGCTAGCAGCGATTGGGTCATGGAGAGTATGGGTAAAGGTACTGACAGTTACGGTGTAACCACCGGGTTTGGAGCTACCTCTCACCGGAGAACAAAAAACGGCACCGCTTTGCAAACCGAACTCATCAGGTTTTTGAACGCCGGAATATTCGGGAACACGAAGGAGACATGCCACACGCTACCGGAATCCGCCACGAGAGCCGCCATGCTCGTCCGAGTCAACACACTCCTCCAAGGATACTCCGGGATCCGATTCGAAATCCTCGAAGCGATCACCAGCCTCCTCAACCACAACATCTCTCCGTCTCTCCCCCTCCGTGGAACCATAACCGCCTCCGGCGATCTCGTCCCCCTCTCCTACATCGCCGGTCTCCTCACCGGCCGTCCAAACTCCAAAGCCACCGGTCCCAACGGCGAATCCCTAACCGCCGAGGAAGCCTTCAAGAAAGCGGGAATCACTTCCGGATTCTTCGATCTACAGCCTAAGGAAGGCTTAGCGCTCGTCAACGGCACGGCGGTTGGATCCGGCATGGCGTCGATGGTTCTCTTCGAAGCGAATGTTCAATCGGTCCTCGCCGAGGTTTTATCCGCGATCTTCGCGGAGGTGATGAGCGGGAAGCCGGAGTTCACCGATCATTTGACTCACCGACTCAAACACCACCCCGGACAAATCGAGGCGGCGGCGATCATGGAGCACATCCTCGACGGAAGCTCGTACATGAAGCTAGCGGCAAAGCTCCACGAGATGGATCCGTTACAAAAACCGAAACAAGACCGTTACGCGCTCCGCACGTCTCCTCAGTGGCTAGGCCCTCAGATCGAAGTGATCCGTCACGCCACGAAGTCAATCGAGCGTGAGATCAACTCCGTTAATGATAATCCGTTAATCGACGTTTCGAGGAACAAAGCGATTCACGGTGGTAACTTCCAGGGGACTCCGATAGGAGTCTCTATGGACAACACACGTTTAGCGGTTGCAGCTATTGGGAAGCTCATGTTCGCTCAGTTCTCGGAGCTTGTTAACGACTTCTACAACAACGGTCTTCCTTCGAATCTAACGGCTTCCAACAACCCAAGCTTGGATTACGGATTCAAAGGAGCGGAGATCGCTATGGCTTCTTACTGCTCTGAGCTTCAGTACCTTGCGAATCCGGTTACAACCCATGTTCAATCAGCTGAGCAGCATAACCAAGATGTTAACTCTTTAGGACTCATCTCGTCTCGTAAGACGTCAGAAGCTGTTGACATTCTCAAGCTCATGTCTACAACGTTCCTCGTAGCGATTTGTCAAGCTGTTGATTTGAGACATCTCGAGGAGAATCTGAGACAGACCGTGAAGAACACGGTTTCTCAAGTGGCGAAGAAAGTGTTGACTACTGGAGTCAACGGGGAGTTGCATCCATCGCGGTTCTGTGAGAAAGACTTGCTTAAGGTCGTTGATCGTGAACAAGTGTTTACATACGTTGACGATCCTTGTAGCGCTACGTACCCATTGATGCAGAAGCTAAGACAAGTTATCGTTGATCACGCGTTATCCAATGGTGAGACTGAGAAGAACGCAGTGACTTCGATCTTTCAAAAGATTGGAGCTTTTGAGGAGGAGCTTAAGATGGTGCTTCCTAAGGAAGTGGATGCGACTAGAGAGGCTTACGGTAATGGAACGGCGGCGATTCCGAACAGGATTAAGGAATGTCGGTCTTATCCATTATATAAGTTTGTGAGGGAAGAGCTCGGGACGAAGTTGCTGACCGGAGAAAAGGTTGTCTCTCCGGGAGAGGAGTTTGATAAGGTGTTCACTGCAATGTGTGAAGGTAAGATTATTGATCCATTGATGGATTGTCTCAAGGAATGGAACGGAGCTCCCATTCCCATATGTTAA

>BnaPAL2-6_A09p036110.1

MDQINSSVHQNGKIEAMLCGGVEKTKVAVAADPLNWGAAAEQMKGSHLDEVKRMVEEYRRPVVNLGGETLTIGQVAAISTVGGGVKVELAEASRAGVKASSDWVMESMGKGTDSYGVTTGFGATSHRRTKNGTALQTELIRFLNAGIFGNTKETCHTLPESATRAAMLVRVNTLLQGYSGIRFEILEAITSFLNHNISPSLPLRGTITASGDLVPLSYIAGLLTGRPNSKATGPNGESLTAEEAFKQAGIASGFFDLQPKEGLALVNGTAVGSGMASMVLFEANVQSVLAEVLSAIFAEVMSGKPEFTDHLTHRLKHHPGQIEAAAIMEHILDGSSYMKLAQKLHEMDPLQKPKQDRYALRTSPQWLGPQIEVIRHATKSIEREINSVNDNPLIDVSRNKAIHGGNFQGTPIGVSMDNTRLAIASIGKLMFAQFSELVNDFYNNGLPSNLTASNNPSLDYGFKGAEIAMASYCSELQYLANPVTSHVQSAEQHNQDVNSLGLISSRKTSEAVDILKLMSTTFLVAICQAVDLRHLEENLRQTVKNTVSQVAKKVLTTGVNGELHPSRFCEKDLLKVVDREQVFTYVDDPCLATYPLMQKLRQVIVDHALSNGETEKNAVTSIFQKIGAFEEELKMVLPKEVDAAREAYGNGTAAIPNRIKECRSYPLYKFVREELGTKLLTGEKVVSPGEEFDKVFTAMCEGKIIDPLMECLKEWNGAPIPIC*

>BnaPAL2-6_A09p036110.1_BnaEXP

ATGGATCAGATTAACAGTTCAGTTCACCAGAACGGCAAGATCGAAGCCATGTTGTGCGGCGGAGTAGAGAAGACGAAGGTGGCTGTGGCGGCGGATCCGTTGAACTGGGGTGCAGCAGCGGAGCAGATGAAAGGGAGTCACTTGGATGAGGTGAAGAGGATGGTTGAGGAGTATCGTAGACCGGTGGTGAATCTCGGAGGAGAGACGCTAACAATCGGACAAGTCGCGGCGATCTCCACCGTAGGCGGAGGTGTTAAGGTTGAGCTAGCGGAGGCTTCGAGAGCCGGCGTGAAAGCTAGCAGTGATTGGGTTATGGAGAGTATGGGCAAAGGTACTGACAGTTACGGTGTCACCACCGGGTTTGGGGCGACCTCTCACCGGAGAACCAAAAACGGCACCGCATTGCAAACCGAACTCATCAGATTTTTGAACGCCGGAATATTCGGTAACACGAAGGAGACATGCCACACGCTACCGGAATCCGCCACGAGAGCCGCCATGCTCGTCCGAGTCAACACTCTCCTCCAAGGATACTCCGGGATCCGATTCGAAATCCTCGAAGCGATCACCAGCTTCCTCAACCACAACATCTCTCCGTCTCTCCCCCTCCGCGGAACCATAACCGCCTCCGGCGATCTCGTCCCCCTCTCCTACATCGCCGGCCTCCTCACCGGCCGTCCGAACTCCAAAGCCACAGGTCCCAACGGCGAATCCCTAACCGCCGAAGAGGCCTTCAAACAAGCCGGAATCGCTTCCGGATTCTTCGATCTACAGCCTAAAGAAGGCTTAGCGCTCGTTAACGGCACGGCGGTTGGATCCGGCATGGCGTCGATGGTTCTATTCGAAGCGAACGTTCAATCGGTGTTAGCGGAGGTCTTATCAGCGATCTTCGCGGAGGTTATGAGCGGGAAGCCTGAGTTCACCGATCATCTGACTCACAGGCTGAAACACCACCCCGGACAAATCGAAGCGGCGGCGATCATGGAGCACATCCTCGACGGAAGCTCGTACATGAAGCTAGCGCAAAAGCTTCACGAGATGGATCCGTTACAAAAACCGAAGCAAGACCGTTACGCTCTCCGTACCTCTCCTCAGTGGCTCGGCCCTCAGATCGAAGTGATCCGTCACGCCACGAAGTCGATCGAGCGTGAGATCAACTCCGTTAACGATAATCCGTTGATCGATGTTTCTAGAAACAAAGCGATTCACGGTGGTAACTTCCAGGGGACTCCGATCGGAGTCTCCATGGATAACACGCGTTTAGCGATCGCTTCGATAGGGAAGCTCATGTTCGCTCAATTCTCGGAGCTTGTTAACGATTTCTATAACAATGGCCTTCCTTCGAATCTAACGGCTTCGAACAATCCAAGCTTGGATTACGGATTCAAAGGAGCGGAGATCGCTATGGCTTCGTATTGCTCTGAGCTTCAGTACTTGGCGAATCCAGTTACAAGCCATGTTCAATCAGCTGAGCAACATAACCAAGATGTTAACTCTTTGGGACTCATCTCGTCTCGCAAAACGTCAGAAGCTGTTGACATTCTCAAGCTGATGTCGACGACGTTCCTTGTGGCTATATGCCAAGCTGTTGATTTGAGACATCTTGAGGAGAATCTGAGACAGACGGTGAAGAACACAGTTTCTCAAGTGGCGAAGAAAGTGTTAACCACTGGAGTCAACGGTGAGCTGCATCCGTCGCGGTTCTGCGAGAAGGACTTGCTTAAGGTTGTTGATCGCGAGCAAGTGTTCACGTACGTGGATGATCCTTGTCTCGCCACGTACCCGTTGATGCAGAAACTGAGACAAGTTATTGTTGATCACGCTTTGTCTAACGGTGAGACTGAGAAGAACGCAGTGACTTCGATCTTTCAAAAGATCGGAGCTTTCGAGGAAGAGCTCAAGATGGTGCTTCCTAAAGAAGTGGATGCGGCTAGAGAGGCTTACGGTAACGGAACGGCGGCGATTCCGAACAGGATTAAGGAATGTCGGTCTTATCCGTTGTATAAGTTCGTGAGGGAAGAGCTTGGAACGAAGTTGTTGACCGGAGAAAAGGTTGTGTCTCCGGGAGAGGAGTTTGATAAGGTGTTCACTGCAATGTGTGAAGGTAAGATCATTGATCCATTGATGGAGTGCCTCAAGGAATGGAACGGAGCTCCGATTCCTATATGCTAA

>BnaPAL2-1_C04p025700.1

MDQINGSVHQNGKTEAMLLCGVVEKTKVTVAADPLNWGAAAEQMKGSHLDEVKRMVEDYRKPVVNLGGETLTIGQVAAISNVGGGVKVELAEASRAGVKASSDWVMESMGKGTDSYGVTTGFGATSHRRTKNGAALQTELIRFLNAGIFGNTKETCHTLPESATRAAMLVRVNTLLQGYSGIRFEILEAITSLLNHNISPSLPLRGTITASGDLVPLSYIAGLLTGRPNSKATGPNGESLTGEEAFKKAGITSGFFDLQPKEGLALVNGTAVGSGMASMVLFEANVQSVLAEVLSAIFAEVMSGKPEFTDHLTHRLKHHPGQIEAAAIMEHILDGSSYMKLAQKLHEMDPLQKPKQDRYALRTSPQWLGPQIEVIRHATKSIEREINSVNDNPLIDVSRNKAIHGGNFQGTPIGVSMDNTRLAVAAIGKLMFAQFSELVNDFYNNGLPSNLTASNNPSLDYGFKGAEIAMASYCSELQYLANPVTSHVQSAEQHNQDVNSLGLISSRKTSEAVDILRLMSTTFLVAICQAVDLRHLEENLRQTVKNTVSQVAKKVLTTGVNGELHPSRFCEKDLLKVVDREQVFTYVDDPCSATYPLMQKLRQVIVDHALSNGEIEKNAVTSIFQKIGAFEEELKMVLPKEVDATREAYANGTAAIPNRIKECRSYPLYKFVREELGTKLLTGEKVVSPGEEFDKVFTAMCEGKIIDPLMDCLKEWNGAPIPIC*

>BnaPAL2-1_C04p025700.1_BnaEXP

ATGGATCAGATCAACGGATCAGTTCACCAAAACGGCAAAACCGAAGCGATGTTGTTGTGCGGCGTAGTAGAGAAGACTAAAGTGACGGTGGCGGCGGATCCGTTAAATTGGGGTGCTGCGGCGGAGCAGATGAAAGGGAGTCATTTAGATGAGGTGAAGAGGATGGTTGAGGATTACCGTAAACCGGTGGTGAATCTCGGCGGAGAAACACTGACGATCGGACAAGTCGCGGCGATCTCGAACGTAGGCGGTGGCGTTAAGGTTGAGCTAGCGGAGGCTTCGAGAGCTGGCGTGAAAGCTAGCAGCGATTGGGTTATGGAGAGTATGGGTAAAGGTACTGACAGTTACGGTGTAACCACTGGGTTTGGAGCTACCTCTCACCGGAGAACCAAAAACGGCGCCGCATTGCAAACCGAACTCATCAGGTTTTTGAACGCCGGAATCTTCGGGAACACGAAGGAGACATGTCACACGCTACCGGAATCCGCCACGAGAGCCGCCATGCTCGTCCGAGTCAACACACTCCTCCAAGGATACTCCGGGATCCGATTCGAAATCCTCGAAGCGATCACCAGCCTCCTCAACCACAACATCTCTCCGTCTCTCCCCCTCCGCGGAACCATAACCGCCTCCGGCGATCTCGTCCCCCTCTCCTACATCGCAGGTCTCCTCACCGGCCGTCCGAACTCCAAAGCCACCGGTCCCAACGGTGAATCCCTAACCGGCGAAGAAGCCTTCAAGAAAGCCGGAATCACTTCCGGATTCTTCGATCTACAGCCTAAGGAAGGCTTAGCGCTCGTCAACGGCACGGCGGTTGGATCCGGCATGGCGTCGATGGTTCTCTTCGAAGCGAATGTTCAATCGGTCCTCGCCGAGGTTTTATCCGCGATCTTCGCGGAGGTGATGAGCGGGAAGCCGGAGTTCACCGATCATTTGACTCACCGACTAAAACATCACCCCGGACAGATCGAAGCGGCGGCGATCATGGAGCACATCCTCGACGGAAGCTCGTACATGAAGCTAGCGCAAAAGCTTCACGAGATGGATCCGTTACAAAAACCGAAACAAGACCGTTACGCGCTCCGCACGTCTCCTCAATGGCTCGGCCCTCAGATCGAAGTGATCCGTCACGCCACGAAGTCAATCGAGCGTGAGATCAACTCAGTTAACGATAATCCGTTAATTGACGTTTCGAGGAACAAAGCGATTCACGGTGGTAACTTCCAGGGGACTCCGATAGGTGTCTCTATGGACAACACACGTTTAGCGGTTGCAGCTATTGGGAAGCTCATGTTCGCTCAGTTCTCTGAGTTGGTGAACGACTTCTATAACAACGGTCTTCCTTCGAATCTAACAGCTTCCAACAACCCAAGTTTAGATTACGGATTCAAAGGAGCAGAGATCGCCATGGCTTCTTACTGCTCTGAGCTTCAGTACCTTGCGAATCCAGTAACAAGCCATGTTCAATCAGCTGAACAGCATAACCAAGATGTTAACTCTTTAGGACTCATCTCCTCTCGTAAGACGTCAGAAGCTGTTGACATTCTCAGGCTGATGTCTACCACGTTCCTCGTAGCGATTTGTCAAGCTGTTGATTTGAGACATCTTGAGGAGAATCTAAGACAAACCGTGAAGAACACTGTTTCTCAAGTGGCGAAGAAAGTGTTGACCACTGGAGTCAACGGGGAGCTGCATCCTTCACGGTTCTGTGAGAAAGACTTGCTTAAGGTCGTTGATCGTGAACAAGTGTTTACATACGTTGACGATCCTTGTAGCGCTACTTACCCGTTGATGCAGAAGTTAAGACAAGTCATTGTCGATCACGCTTTATCCAATGGTGAGATTGAGAAGAACGCAGTGACTTCGATCTTTCAAAAGATTGGGGCTTTTGAGGAGGAGCTTAAGATGGTGCTTCCAAAGGAAGTGGATGCGACTAGAGAGGCTTACGCTAATGGAACGGCGGCGATTCCCAACAGGATTAAGGAATGTCGATCTTATCCGCTGTATAAGTTCGTGAGGGAAGAGCTCGGAACGAAGTTGTTGACCGGAGAAAAGGTTGTCTCTCCGGGAGAGGAGTTTGATAAGGTATTCACTGCAATGTGTGAAGGTAAGATTATTGATCCATTGATGGATTGTCTCAAGGAATGGAACGGAGCTCCCATTCCCATATGTTAA

>BnaPAL2-5_C08p028640.1

MDQTNGSVHQNGKIEAMLCGGVEKTKVAVAADPLNWGAAAEQMKGSHLDELKRMVEEYRRPVVNLGGETLTIGQVAAISTAGGGVKVELAEASRAGVKASSDWVMESMGKGTDSYGVTTGFGATSHRRTKNGTALQTELIRFLNAGIFVNTLLQGYSGIRFEILEAITSFLNHNISPSLPLRGTITASGDLVPLSYIAGLLTGRPNSKATGPNGESLTAEEAFKQAGIASGFFDLQPKEGLALVNGTAVGSGMASMVLFEANVQSVLAEVLSAIFAEVMSGKPEFTDHLTHRLKHHPGQIEAAAIMEHILDGSSYMKLAQKLHEMDPLQKPKQDRYALRTSPQWLGPQIEVIRHATKSIEREINSVNDNPLIDVSRNKAIHGGNFQGTPIGVSMDNTRLAIASIGKLMFAQFSELVNDFYNNGLPSNLTASNNPSLDYGFKGAEIAMASYCSELQYLANPVTSHVQSAEQHNQDVNSLGLISSRKTSEAVDILKLMSTTFLVAICQAVDLRHLEENLRQTVKNTVSQVAKKVLTTGVNGELHPSRFCEKDLLKVVDREQVFTYVDDPCSATYPLMQKLRQVIVDHALSNGETEKNAVTSIFQKIGAFEEELKMVLPKEVDAAREAYGNGTAAIPNRIKECRSYPLYKFVREELGTKLLTGEKVVSPGEEFDKVFTAMCEGKIIDPLMECLKEWNGAPIPIC*

>BnaPAL2-5_C08p028640.1_BnaEXP

ATGGATCAGACTAACGGTTCAGTTCACCAGAACGGCAAGATCGAAGCCATGTTGTGCGGTGGAGTAGAGAAGACAAAAGTGGCTGTGGCGGCGGATCCGTTGAACTGGGGTGCAGCAGCGGAACAGATGAAAGGGAGTCACTTGGATGAGTTGAAGAGGATGGTTGAGGAGTACCGCAGACCGGTGGTGAATCTCGGGGGAGAGACGCTAACAATCGGACAAGTCGCGGCGATCTCCACCGCAGGCGGAGGTGTTAAGGTTGAGCTAGCGGAGGCTTCGAGAGCTGGCGTGAAAGCTAGCAGTGATTGGGTTATGGAAAGTATGGGCAAAGGTACTGACAGTTACGGTGTCACCACCGGGTTTGGAGCGACCTCTCACCGGAGAACCAAAAACGGCACCGCATTGCAAACGGAACTCATCAGATTTTTGAACGCCGGAATATTCGTCAACACTCTCCTCCAAGGATACTCCGGGATCCGATTCGAAATCCTCGAAGCGATCACCAGCTTCCTCAACCACAACATCTCTCCATCTCTCCCCCTCCGTGGAACCATAACCGCCTCCGGCGATCTCGTCCCCCTCTCCTACATCGCCGGCCTCCTCACCGGCCGTCCGAACTCCAAAGCCACAGGTCCCAACGGCGAATCCCTAACCGCCGAAGAGGCCTTCAAACAAGCCGGAATCGCTTCCGGATTCTTCGATCTACAACCCAAGGAAGGCTTAGCGCTCGTTAACGGCACGGCGGTTGGATCCGGCATGGCGTCGATGGTTCTATTCGAAGCGAACGTTCAATCGGTGTTAGCGGAGGTCTTATCCGCGATCTTCGCGGAGGTGATGAGCGGGAAGCCTGAGTTCACCGATCATCTAACGCACAGGCTGAAGCACCACCCCGGGCAAATCGAAGCGGCGGCGATCATGGAGCACATCCTCGACGGAAGCTCGTACATGAAGCTAGCGCAGAAGCTTCACGAGATGGATCCGTTACAAAAGCCGAAACAAGACCGTTACGCTCTCCGTACCTCTCCTCAGTGGCTCGGCCCTCAGATCGAAGTGATCCGTCACGCAACGAAGTCGATCGAGCGTGAGATCAACTCCGTTAACGATAATCCGTTGATCGATGTTTCTAGAAACAAAGCGATTCACGGTGGTAACTTCCAGGGGACTCCAATCGGAGTCTCCATGGATAACACTCGTTTAGCGATTGCTTCGATAGGGAAGCTCATGTTCGCTCAATTCTCGGAGCTTGTTAACGATTTCTACAACAACGGTCTTCCTTCGAATCTAACGGCTTCGAACAATCCAAGCTTGGATTACGGATTCAAAGGAGCTGAGATCGCTATGGCTTCGTATTGCTCTGAGCTTCAGTACTTGGCGAATCCAGTTACAAGCCATGTTCAATCAGCTGAGCAACATAACCAAGATGTGAACTCTTTGGGACTCATCTCGTCTCGCAAAACGTCAGAAGCTGTTGACATCCTCAAGCTTATGTCGACAACGTTCCTTGTGGCTATATGCCAAGCTGTTGATTTGAGACATCTTGAGGAGAATCTGAGACAAACGGTGAAGAACACAGTTTCTCAAGTGGCGAAGAAAGTTTTAACCACTGGAGTCAACGGTGAGCTGCATCCGTCGCGGTTCTGCGAGAAGGACTTGCTTAAGGTCGTTGATCGCGAGCAAGTGTTCACGTACGTGGATGATCCTTGTAGCGCTACGTACCCGTTGATGCAGAAACTGAGACAAGTTATTGTTGATCATGCTTTGTCTAACGGTGAGACTGAGAAGAACGCAGTGACTTCGATCTTTCAAAAGATCGGAGCTTTCGAGGAAGAGCTCAAGATGGTGCTCCCTAAAGAAGTGGATGCGGCTAGAGAGGCTTACGGTAACGGAACGGCGGCGATTCCGAACAGGATTAAGGAATGTCGGTCTTACCCGTTGTATAAGTTCGTGAGGGAAGAGCTTGGAACGAAGCTGTTGACCGGAGAAAAGGTTGTGTCTCCGGGAGAGGAGTTTGATAAGGTGTTCACTGCAATGTGTGAAGGTAAGATCATTGATCCGTTGATGGAGTGCCTCAAGGAATGGAACGGAGCTCCGATTCCGATATGCTAA

>BnaPAL2-3_R186_chrC06p000250.1

MDQTQNDNIEAMLCGGVEKTNVAVAADPLNWGAAAEQMKGSHLDEVKRMVEEYRRPVVNLGGETLTIGQVAAISTVGNGVKVELAEASRDGVKASSDWVMESMGKGTDSYGVTTGFGATSHRRTKNGAALQTELIRFLNAGIFVNTLLQGYSGIRFEILEAITSLLNHNISPSLPLRGTITASGDLVPLSYIAGLLTGRPNSKATGPNGESLTAEEAFKQAGITSGFFDLQPKEGLALVNGTAVGSGMASMVLFEANVQSVLAEVLSAIFAEVMSGKPEFTDHLTHRLKHHPGQIEAAAIMEHILDGSSYMKLAQKLHEMDPLQKPKQDRYALRTSPQWLGPQIEVIRYATKSIEREINSVNDNPLIDVSRNKAIHGGNFQGTPIGVSMDNTRLAVAAIGKLMFAQFSELVNDFYNNGLPSNLTASNNPSLDYGFKGAEIAMASYCSELQYLANPVTSHVQSAEQHNQDVNSLGLISSRKTSEAVDILKLMSTTFLVAICQAVDLRHLEENLRQAVKNTVSQVAKKVLTTGVNGEMHPSRFCERDLLKVVDREQVFTYVDDPCSATYPLMQKLRQVIVDQALANGETEKNVETSIFQKIGAFEEELKTVLPKEVDAAREAYGNGNAAIPNRIKECRSYPLYKFVREELGTKLLTGEKVVSPGEEFDKVFTAMCEGKIIDPLMDCLKEWNGAPIPIC*

>R186_chrC06p000250.1_BnaEXP

ATGGATCAGACCCAAAACGACAACATCGAAGCTATGTTGTGCGGCGGAGTTGAGAAGACGAACGTTGCTGTGGCGGCAGATCCGTTGAACTGGGGTGCTGCGGCGGAGCAGATGAAAGGGAGCCATTTGGATGAGGTGAAGAGGATGGTTGAGGAGTACCGTAGACCGGTGGTGAACCTCGGAGGAGAGACGCTGACGATCGGACAAGTGGCGGCGATCTCCACCGTCGGTAACGGCGTTAAGGTTGAGCTAGCGGAGGCTTCGAGAGATGGAGTGAAAGCTAGCAGCGATTGGGTTATGGAGAGTATGGGCAAAGGTACCGACAGTTACGGTGTCACCACCGGGTTTGGAGCTACCTCTCACCGGAGAACGAAAAACGGAGCTGCATTGCAGACGGAGCTCATCAGATTTTTGAACGCCGGAATATTCGTCAACACACTCCTCCAAGGATACTCCGGGATCCGATTCGAAATCCTCGAAGCGATCACCAGTCTCCTCAACCACAACATCTCTCCATCTCTCCCCCTACGTGGAACCATAACCGCCTCCGGCGATCTCGTCCCCCTCTCCTACATCGCCGGTCTGCTCACCGGACGCCCCAACTCCAAAGCCACCGGTCCCAACGGCGAATCCCTAACCGCCGAAGAAGCCTTCAAACAAGCCGGAATCACTTCCGGATTCTTCGATCTGCAACCTAAGGAAGGTCTCGCGCTCGTTAACGGCACGGCGGTTGGATCCGGCATGGCGTCGATGGTTCTATTCGAAGCGAATGTTCAATCCGTTTTAGCGGAGGTTTTATCGGCGATCTTCGCGGAGGTGATGAGCGGGAAGCCTGAGTTCACCGATCATTTGACTCACCGACTCAAACATCACCCCGGACAAATCGAAGCGGCGGCGATCATGGAGCACATCCTCGACGGAAGCTCGTACATGAAGCTAGCTCAAAAGCTTCACGAGATGGATCCGTTACAGAAGCCCAAACAAGACCGTTACGCTCTCCGTACTTCTCCTCAATGGCTCGGCCCTCAGATCGAAGTGATCCGTTACGCCACGAAATCAATCGAGCGTGAGATCAACTCCGTTAACGATAATCCGTTGATCGACGTTTCTAGGAACAAAGCCATTCACGGTGGTAACTTCCAGGGGACGCCGATTGGAGTTTCGATGGACAACACGCGTTTAGCGGTTGCAGCCATTGGGAAGCTCATGTTTGCTCAGTTCTCTGAGCTGGTCAACGATTTCTACAATAATGGCCTTCCTTCGAATCTCACAGCTTCTAACAATCCAAGTTTGGATTACGGTTTCAAAGGAGCAGAGATCGCTATGGCTTCTTACTGCTCTGAGCTTCAGTACTTGGCGAATCCAGTTACTAGCCATGTTCAATCAGCTGAACAGCATAACCAAGACGTTAACTCTTTAGGACTCATCTCATCTCGTAAGACCTCAGAAGCTGTTGACATTCTCAAGCTAATGTCTACCACGTTCCTTGTTGCTATATGCCAAGCTGTTGACTTGAGACATCTTGAGGAGAATCTGAGACAGGCGGTGAAGAACACAGTTTCTCAAGTGGCGAAGAAAGTGTTGACTACTGGAGTCAACGGGGAGATGCATCCTTCACGGTTCTGTGAGAGAGACTTGCTTAAGGTCGTTGACCGTGAGCAAGTGTTCACGTACGTGGATGATCCTTGTAGCGCAACTTACCCACTGATGCAGAAGCTAAGACAAGTCATTGTTGATCAAGCTTTAGCCAACGGTGAGACTGAGAAGAACGTAGAGACTTCAATTTTTCAAAAGATTGGAGCTTTTGAGGAGGAGCTTAAGACGGTTCTTCCTAAGGAAGTGGATGCGGCTAGAGAGGCTTACGGTAATGGAAACGCAGCGATTCCGAACAGGATTAAGGAGTGTCGGTCTTATCCGTTGTATAAGTTCGTGAGGGAGGAGCTTGGAACGAAGTTGTTGACGGGAGAAAAGGTTGTGTCTCCGGGAGAGGAGTTTGATAAGGTGTTCACTGCTATGTGTGAAGGTAAGATTATTGATCCACTGATGGATTGTCTCAAGGAATGGAACGGAGCTCCGATTCCGATATGCTAA

>BnaPAL2-4_A07p016620.1

MDQTQNDNIETMLCGGVEKTNVAVAADPLNWGAAAEQMKGSHLDEVKRMVEEYRRPVVNLGGETLTIGQVAAISTVGNGVKVELAEASRAGVKASSDWVMESMGKGTDSYGVTTGFGATSHRRTKNGVALQTELIRFLNAGIFVNTLLQGYSGIRFEILEAITSLLNHNISPSLPLRGTITASGDLVPLSYIAGLLTGRPNSKATAPNGESLTAEEAFKQAGIASGFFDLQPKEGLALVNGTAVGSGMASMVLFEANVQSVLAEVLSAIFAEVMSGKPEFTDHLTHRLKHHPGQIEAAAIMEHILKGSSYMKLAQKVHEMDPLQKPKQDRYALRTSPQWLGPQIEVIRYATRSIEREINSVNDNPLIDVSRNKAIHGGNFQGTPIGVSMDNTRLAVAAIGKLMFAQFSELVNDFYNNGLPSNLTASNNPSLDYGFKGAEIAMASYCSELQYLANPVTSHVQSAEQHNQDVNSLGLISSRKTSEAVDILKLMSTTFLVAICQAVDLRHLEENLRQAVKNTVSLVAKKVLTTGVNGEMHPSRFCERDLLKVVDREQVFTYVDDPCSASYPLMQKLRQVIVDHALANGETEKNVETSIFEKIGAFEEELKTVLPKEVDAAREAYGNGNAAIPNRIKECRSYPLYKFVREELGTKLLTGEKVVSPGEEFDKVFIAMCEGKIIDPLMDCLKEWNGAPIPIC*

>BnaPAL2-4_A07p016620.1_BnaEXP

ATGGATCAGACACAAAACGACAACATCGAAACTATGTTGTGCGGCGGAGTTGAGAAGACGAACGTGGCTGTGGCGGCAGATCCGTTGAACTGGGGTGCTGCGGCGGAGCAGATGAAAGGGAGCCACTTGGATGAGGTGAAGAGGATGGTTGAGGAGTACCGTAGACCGGTGGTGAACCTCGGAGGAGAGACGCTAACGATCGGACAAGTGGCGGCGATCTCCACCGTCGGTAACGGCGTTAAGGTTGAGCTAGCGGAGGCTTCGAGAGCGGGAGTGAAAGCTAGCAGCGATTGGGTTATGGAGAGTATGGGCAAAGGTACTGACAGTTACGGTGTCACCACCGGGTTTGGAGCTACCTCTCACCGTAGAACGAAAAACGGGGTCGCATTGCAGACAGAACTCATCAGATTTTTGAACGCCGGAATATTCGTCAACACACTCCTCCAAGGATACTCCGGGATCCGATTCGAAATCCTCGAAGCGATCACCAGTCTCCTCAACCACAACATCTCCCCATCTCTCCCCCTCCGTGGAACCATAACCGCCTCCGGCGACCTCGTCCCCCTCTCCTACATCGCCGGTCTCCTCACCGGACGCCCCAACTCCAAAGCCACCGCTCCCAACGGCGAATCCCTAACCGCCGAAGAAGCCTTCAAACAAGCCGGAATCGCTTCCGGATTCTTCGATCTTCAACCCAAGGAAGGTCTCGCGCTCGTTAACGGCACGGCGGTTGGATCCGGCATGGCCTCGATGGTTCTATTCGAAGCGAACGTTCAGTCCGTTTTAGCGGAGGTTTTATCGGCGATCTTCGCGGAGGTGATGAGCGGGAAGCCTGAGTTCACCGATCATTTGACTCACCGACTCAAACATCACCCCGGACAAATCGAAGCGGCGGCGATCATGGAGCACATCCTCAAGGGAAGCTCGTACATGAAGCTAGCTCAAAAGGTTCACGAGATGGATCCGTTACAGAAGCCCAAACAAGACCGTTACGCTCTCCGTACTTCTCCTCAATGGCTCGGCCCTCAGATCGAAGTGATCCGTTACGCGACGAGATCAATCGAGCGTGAGATCAACTCCGTTAACGATAATCCGTTGATCGACGTTTCTAGGAACAAAGCGATTCACGGTGGTAACTTCCAGGGGACGCCGATTGGAGTTTCGATGGACAACACGCGTTTAGCGGTTGCGGCCATTGGGAAGCTCATGTTTGCTCAGTTCTCGGAGCTTGTTAACGATTTCTACAACAATGGTTTACCTTCGAACTTAACAGCTTCGAATAATCCAAGTTTGGATTACGGTTTCAAAGGAGCTGAGATCGCTATGGCTTCTTACTGCTCTGAGCTTCAGTACTTAGCGAATCCAGTAACTAGCCATGTTCAATCAGCTGAGCAGCATAACCAAGACGTTAACTCTTTAGGACTCATCTCATCTCGTAAGACCTCAGAAGCTGTTGACATTCTCAAGCTGATGTCTACGACGTTCCTTGTTGCTATATGCCAAGCTGTTGACTTGAGACATCTTGAGGAGAATCTGAGACAGGCGGTGAAGAACACAGTTTCTCTAGTGGCGAAGAAAGTGTTGACCACTGGAGTCAACGGGGAGATGCATCCGTCACGGTTCTGTGAGAGAGACTTGCTTAAGGTCGTTGACCGTGAGCAAGTGTTCACGTACGTGGATGATCCTTGCAGCGCAAGTTACCCGTTGATGCAGAAGCTAAGACAAGTCATTGTTGATCACGCTTTAGCCAACGGTGAGACTGAGAAGAATGTGGAGACTTCAATCTTTGAAAAGATTGGAGCTTTTGAGGAGGAGCTTAAGACGGTTCTTCCTAAGGAAGTGGATGCGGCTAGAGAGGCTTACGGTAATGGAAACGCAGCGATTCCGAACAGGATTAAGGAGTGTCGGTCTTATCCGTTGTATAAGTTCGTGAGGGAGGAGCTTGGAACGAAGTTGTTGACCGGAGAAAAGGTTGTGTCTCCGGGAGAGGAGTTTGATAAGGTGTTCATTGCTATGTGTGAAGGTAAGATTATTGATCCACTGATGGATTGTCTCAAGGAATGGAACGGAGCTCCGATTCCGATATGCTAA

>BnaPAL4-2_A05p031299.1

MDLCKQNNNHIVAVSADPLNWNAAAEALKGSHLEEVKRMVEDYRKGAVRLGGETLTIGQVAAVASGGVTVELAEEARAGVKASSDWVMESMNRGTDSYGVTTGFGATSHRRTKQGGALQKELIRFLNAGIFGAGVGDTSLTLPKSATRAAMLVRVNTLLQGYSGIRFEILEAITKLLNNEITPCIPLRGSITASGDLVPLSYIAGLLTGRPNSKAVGPAGETLTASDAFKLAGVPSFFELQPKEGLALVNGTGVGSGLASMVLFETNVLAVLSEVMSAMFAEVMQGKPEFTDHLTHKLKHHPGQIEAAAIMEHILHGSSYVKEAQQLHELDPLQKPKQDRYALRTSPQWLGPQIEVIRAATKMIEREINSVNDNPLIDVSRNKALHGGNFQGTPIGVAMDNTRLAIASIGKLMFAQFSELVNDFYNNGLPSNLSGGRNPSLDYGFKGAEIAMASYCSELQFLANPVTNHVQSAEQHNQDVNSLGLISSRKTEEAVDILKLMSTTYLVALCQAVDLRHIEENLKKAVKAAVSQVAKRVLTVGVNGELHPSRFTERDVLQVVDREHVFSYADDPCSFAYPLMQRLRHVLVDHALEDPDREANVSTSVFQKIEAFEAELKVVLPKEVERVRVEYEGGCSVVGNRIKECRSYPLYRFVREELETELLSGESVRSPGEEFDKVFSAICDGKVIDPLLECLKEWNGAPVPIC

>BnaPAL4-2_A05p031299.1_BnaEXP

atggacttgtgcaaacaaaacaacaaccacatcgtcgctgtttccgccgatccgttgaactggaacgcggcggcggaagctttgaaagggagccacttggaggaggtgaaacgtatggtggaagattacagaaaaggtgcggtgcggttaggaggagagacgctgacgattggtcaagttgcagccgtggctagcggaggagtgacggtggagctggcggaggaggctcgtgccggagttaaggcgagtagcgactgggtgatggagagcatgaaccgtggcacggacagttatggagtcaccacagggtttggtgcaacgtcccatagaagaactaaacaaggcggtgcacttcaaaaggagcttattaggttcttgaacgccggaatattcggtgccggcgtcggagacacgtcactcacgctacctaagtcggcaactagagcagctatgctcgtccgtgtcaacactctcctccaaggctactctggaatacgctttgagattcttgaagccattacaaagcttcttaacaacgaaatcactccatgcatccctctccgtggcagcatcaccgcatccggcgaccttgttcctctctcttacatcgccggactgctcaccggccgtcccaattcaaaagccgtgggtcccgccggtgagactctcactgcctccgatgcctttaagctagccggagtaccgtccttttttgagctgcagccaaaggaaggactggctcttgtgaacgggacaggggtcggatcgggtctggcttcgatggttctgtttgagaccaatgttttggcggttttgtctgaagttatgtctgcgatgttcgcggaggtcatgcaagggaagccggagtttactgatcatctaacgcataagctcaagcaccatcctggtcagattgaagccgctgcgattatggaacatatattacatggaagctcttacgttaaagaagctcagcagcttcatgaactggatccgcttcaaaaacctaaacaagatcggtacgcgctaaggacgtcaccacagtggctaggaccacaaatcgaagtgatcagagcagcgaccaagatgatagagcgcgagatcaactcggtcaacgacaaccctctcatcgacgtctccagaaacaaagcactacacggcggtaacttccaaggcacgcctatcggtgtcgccatggacaacacccgcttagccatcgcttccatcggcaaactcatgttcgcgcagttctccgagctcgtgaacgacttctacaacaacggcttgccttcaaacctatccggcgggagaaaccctagcctcgactacggtttcaaaggcgcggagatcgccatggcctcttactgctccgagctccagttcttggctaaccccgtgacgaaccacgtccagagcgccgagcagcataaccaagacgttaactcgctagggctgatctcgagtcggaagacagaagaagctgttgatatcctcaagctgatgtccacgacctacttggttgccttatgccaagccgttgatctgcgtcatatcgaagagaatctcaagaaagctgttaaagccgcggtgagccaggtggcgaaacgggtgttaacggttggtgttaacggcgagctgcatccgtctaggttcacggagcgtgatgtgctccaagtggttgatagagagcacgtgttctcatacgcggacgatccttgcagctttgcttatccgttgatgcagaggcttaggcacgttcttgtggaccacgctttggaggatccggaccgcgaggctaatgtttcgacgtcggtttttcagaagatagaagcgtttgaggcggagctgaaggtggttttacctaaggaagtggagcgtgttagggtggagtatgagggagggtgttcggttgtgggtaaccggattaaggaatgtcggtcttatccgttgtaccggtttgtgagggaggagctggagactgagctgttgagtggagagagtgttaggtcgccgggtgaggagtttgataaagtgttctcggcgatttgtgatgggaaggttattgatcctttgttggagtgtctcaaggagtggaacggagctccggttccgatctgttga

>BnaPAL4-1_C05p041260.1

MELCEQNNNHVVAVSADPLNWNAAAEALKGSHLEEVKRMVEDYRKGAVRLGGETLTIGQVAAVASGGVMVELAEEARAGVKASSDWVMESMNRGTDSYGVTTGFGATSHRRTKQGGALQKELIRFLNAGIFGAGAGDSSLTLPKSATRAAMLVRVNTLLQGYSGIRFEILEAITKLLNNEITPCIPLRGSITASGDLVPLSYIAGLLTGRPNSKAVGPAGETLTASDAFKLAGVPSFFELQPKEGLALVNGTGVGSGLASMVLFEANVLAVLSEVMSAMFAEVMQGKPEFTDHLTHKLKHHPGQIEAAAIMEHILHGSSYVKEAQQLHELDPLQKPKQDRYALRTSPQWLGPQIEVIRAATKMIEREINSVNDNPLIDVSRNKALHGGNFQGTPIGVAMDNTRLAIASIGKLMFAQFSELVNDFYNNGLPSNLSGGRNPSLDYGFKGAEIAMASYCSELQFLANPVTNHVQSAEQHNQDVNSLGLISSRKTEEAVEILKLMSTTYLVALCQAVDLRHIEENLKKAVKAAVSQVAKRVLTVGVNGELHPSRFTERDVLQVVDREHVFSYADDPCSFAYPLMQRLRHVLVDHALEDPDREADASASVFQKIGAFETELKVVLPKEVERVRGEYEGGRSVVGNRIKECRSYPLYRFVREELETELLSGESVRSPGEVFDKVFSAICDGKVIDPLLECLKEWNGAPVPIC*

>BnaPAL4-1_C05p041260.1_BnaEXP

ATGGAGTTATGCGAACAAAACAACAACCACGTCGTCGCTGTCTCCGCCGATCCGTTGAACTGGAACGCGGCGGCGGAAGCTTTGAAAGGGAGCCACTTGGAGGAGGTGAAACGTATGGTGGAAGATTACAGAAAAGGGGCGGTGCGGTTAGGAGGAGAGACGCTGACGATTGGTCAAGTTGCCGCCGTTGCTAGCGGAGGAGTGATGGTGGAGCTGGCGGAGGAGGCTCGTGCCGGAGTTAAGGCGAGTAGCGACTGGGTGATGGAGAGCATGAACCGTGGCACGGACAGTTATGGAGTCACCACGGGGTTTGGTGCAACGTCCCATAGAAGAACTAAACAAGGCGGTGCACTTCAAAAGGAGCTTATTAGGTTCTTGAACGCCGGGATATTCGGTGCCGGCGCCGGAGACTCGTCACTCACGCTACCTAAGTCGGCGACTAGAGCAGCTATGCTCGTCCGTGTCAACACTCTCCTCCAAGGCTACTCTGGAATACGTTTTGAGATTCTTGAAGCCATTACAAAGCTTCTTAACAACGAAATCACTCCATGCATCCCTCTCCGTGGCAGCATCACTGCATCCGGCGACCTTGTTCCCCTCTCTTACATCGCCGGACTACTCACCGGCCGTCCCAATTCAAAAGCCGTGGGTCCCGCCGGTGAGACTCTCACTGCCTCTGATGCCTTTAAGCTAGCCGGAGTACCGTCCTTTTTCGAGCTGCAGCCAAAGGAAGGACTGGCTCTTGTGAACGGGACAGGGGTCGGATCGGGTCTGGCCTCGATGGTTCTGTTTGAGGCCAACGTTTTGGCGGTTTTGTCTGAAGTTATGTCTGCCATGTTCGCGGAGGTTATGCAAGGCAAACCGGAGTTTACTGATCATCTAACGCATAAGCTCAAGCACCATCCTGGTCAGATTGAAGCCGCTGCGATTATGGAACATATATTACATGGAAGCTCTTACGTTAAAGAAGCTCAACAGCTCCATGAACTGGATCCGCTTCAAAAACCTAAACAAGATCGGTACGCGCTAAGGACGTCACCACAATGGCTAGGACCACAAATCGAAGTGATCAGAGCAGCGACCAAGATGATAGAGCGTGAGATCAACTCGGTCAACGACAACCCTCTCATAGACGTCTCCAGAAACAAAGCACTACACGGCGGTAACTTCCAAGGCACGCCTATCGGTGTTGCCATGGACAACACCCGCTTAGCCATCGCTTCCATCGGCAAACTCATGTTCGCGCAGTTCTCCGAGCTCGTGAACGACTTCTACAACAACGGCTTGCCTTCAAACCTATCCGGCGGGAGAAACCCTAGCCTCGACTACGGTTTCAAAGGCGCGGAGATAGCCATGGCCTCTTACTGCTCCGAGCTCCAGTTCTTGGCTAACCCCGTGACTAACCACGTCCAGAGCGCCGAGCAGCATAACCAAGACGTTAACTCGCTAGGGCTGATCTCGAGCCGGAAGACAGAAGAAGCTGTTGAGATCCTCAAGCTGATGTCCACGACCTACTTGGTTGCGCTCTGCCAAGCCGTTGATCTGCGTCATATCGAAGAGAATCTCAAGAAAGCGGTTAAAGCCGCTGTGAGCCAGGTGGCGAAACGGGTGTTAACGGTTGGTGTTAACGGCGAGCTGCATCCGTCGAGGTTCACGGAGCGGGATGTGCTCCAAGTGGTTGATAGAGAGCACGTGTTCTCCTACGCGGACGATCCTTGCAGCTTTGCTTACCCGTTGATGCAGAGGCTTAGGCACGTTCTTGTGGACCACGCTTTAGAGGATCCGGACCGCGAGGCTGATGCTTCGGCGTCGGTTTTTCAGAAGATAGGAGCGTTTGAGACGGAGCTGAAGGTGGTTTTACCTAAGGAGGTGGAGCGTGTTAGGGGTGAGTATGAGGGAGGGCGGTCGGTTGTGGGTAACCGGATTAAGGAATGTCGGTCTTATCCGTTGTACCGGTTTGTGAGGGAGGAGCTGGAGACTGAGCTGTTGAGTGGAGAGAGTGTTAGGTCGCCCGGTGAAGTGTTTGATAAAGTGTTCTCGGCGATTTGTGATGGGAAGGTTATTGATCCTTTGTTGGAGTGTCTCAAGGAGTGGAACGGAGCTCCGGTTCCGATCTGTTGA

>BnaPAL3-2_A04p012590.1

MEFCQPNKNNGSASSDPLNWNVAAEALKGSHVEDVKKMVEDYRKGTVRLGGETLTIGQVAAVASKGTTVELLEEARAGVKASSEWVMESINRGTDTYGITTGFGSSSRRRTNQGAALQKELIRYLNTGIFATGDEDDVLSNILPRPATRAAMLIRVNTLLQGYSGIRFEILEAITKFLNHKITLRLPLRGTITASGDLVPLSYIAGLLTGRPNSRSVGPSDEILTALEAFKLAGISSPFELRPKEGLALVNGTAVGSAMASIVLYEANVLAVFSEVASAMFVEVMHGKPEFTDHLVHKLKHHPGQIEAAAIMEHILDGSSFVKEAIRLHEIDPLQKPKQDRYALRTSPQWLGPQIEVIRAATKMIEREINSVNDNPLIDVLRNKAIHGGNFQGTPIGVAMDNARLAIGSIGKLMFAQFTELVNDFYNNGLPSNLSGGRNPSLDYGFKGAEVPMAAYCSELQFLANPVTNHVQSTEQHNQDVNSLGLVSSHKTAEAVNILKLMSATYLVALCQAYDLRHLEDNLKETIKAVVNQTAERHVFTLSKPFLEQNILGVIDREYVFSYVYDLSSLTNPLMQKLRSILFDHALAEPEHETDSGFRKIGTFETELKSLLHNEVERVWTEYEKGNFVVANRIKECRSYPLYRFVREELETRLLTGGSVRTPGEDFDEVFKAISKGKLIDPLFECLKEWNGAPIPIS*

>BnaPAL3-2_A04p012590.1_BnaEXP

ATGGAATTTTGTCAACCAAACAAAAACAACGGAAGTGCGTCTAGTGATCCACTTAATTGGAACGTAGCGGCCGAGGCTTTGAAAGGGAGCCACGTGGAGGATGTGAAGAAGATGGTGGAGGATTATAGGAAAGGAACGGTGCGGCTCGGTGGAGAGACGCTGACTATCGGTCAGGTTGCGGCCGTTGCGAGCAAAGGGACGACGGTGGAGCTTTTGGAGGAGGCTCGTGCCGGCGTGAAGGCTAGTAGTGAGTGGGTGATGGAAAGCATTAACCGTGGCACGGACACTTATGGAATCACCACTGGATTTGGTTCTTCTTCTCGTAGGAGGACCAATCAAGGTGCTGCTCTTCAAAAAGAGCTTATTAGGTACTTGAACACCGGAATATTCGCTACAGGCGACGAAGATGACGTTTTGTCAAACATTCTTCCTCGTCCGGCAACAAGAGCGGCGATGCTCATCCGTGTCAACACACTCCTCCAAGGCTACTCCGGTATACGCTTCGAAATCCTTGAAGCCATCACAAAGTTCCTCAACCACAAAATCACACTGCGCCTCCCTCTCCGAGGCACCATCACCGCCTCTGGTGACCTCGTTCCTCTATCGTACATCGCCGGTCTCCTCACCGGACGGCCCAACTCCCGATCCGTGGGTCCATCCGATGAGATCCTTACTGCCTTAGAGGCCTTCAAGCTAGCTGGAATATCTTCTCCTTTTGAGCTCCGGCCTAAAGAAGGGCTTGCGCTCGTGAACGGCACCGCGGTTGGGTCCGCTATGGCCTCAATAGTACTATACGAGGCCAACGTTCTGGCAGTTTTTTCTGAAGTTGCTTCCGCCATGTTTGTTGAGGTTATGCATGGGAAACCCGAGTTTACCGATCATCTTGTGCATAAACTCAAGCACCATCCTGGTCAGATCGAAGCCGCGGCAATCATGGAACATATCTTAGACGGAAGCTCTTTTGTCAAAGAAGCTATACGTCTCCATGAGATTGATCCGCTCCAAAAACCTAAACAAGATCGCTACGCTCTGCGAACATCGCCACAGTGGCTTGGACCGCAGATTGAGGTGATCAGAGCCGCGACGAAAATGATTGAACGAGAGATAAACTCTGTCAACGATAACCCTTTAATCGATGTGCTGCGAAACAAAGCTATCCACGGTGGGAATTTCCAGGGGACACCAATAGGTGTAGCCATGGACAACGCTCGTCTGGCCATTGGTTCCATCGGGAAACTAATGTTTGCGCAATTCACGGAACTCGTCAATGATTTCTATAACAACGGTCTACCTTCGAATCTATCCGGTGGGAGAAACCCTAGCCTTGACTACGGGTTCAAAGGCGCGGAAGTCCCCATGGCTGCTTATTGTTCCGAGCTTCAGTTCTTGGCTAACCCTGTGACAAACCATGTCCAAAGCACTGAGCAACATAATCAAGATGTTAACTCTCTTGGTTTGGTCTCCAGCCACAAGACTGCAGAAGCTGTGAATATCCTCAAGCTTATGTCAGCTACTTACTTAGTAGCTTTATGCCAAGCCTATGATCTAAGACATCTTGAAGATAATCTCAAGGAAACGATTAAGGCGGTTGTGAACCAAACCGCGGAAAGACACGTATTCACGCTCAGCAAACCGTTCCTCGAACAGAACATCCTCGGAGTTATCGACCGCGAATATGTCTTTTCCTATGTTTATGACCTAAGCAGCCTCACTAACCCTCTAATGCAGAAGCTGAGAAGCATTCTTTTCGACCATGCTTTAGCTGAACCGGAACATGAGACGGATTCGGGTTTTCGAAAAATAGGAACATTCGAAACAGAGCTGAAATCTCTTCTACATAACGAAGTTGAAAGAGTGTGGACCGAGTATGAAAAGGGTAACTTTGTTGTGGCTAACCGGATCAAAGAGTGTAGATCGTATCCGTTGTACCGGTTTGTACGGGAGGAGCTAGAGACGAGGCTACTAACCGGAGGGAGTGTCCGGACACCAGGTGAAGATTTCGACGAAGTTTTTAAAGCCATCTCTAAGGGAAAACTCATAGATCCTCTGTTTGAATGCCTCAAGGAATGGAACGGAGCTCCAATCCCAATCTCCTAA

>BnaPAL3-1_C04p034430.1

MLIRVNTLLQGYSGIRFEILEAITKFLNHKITPRLPLRGTITASGDLVPLSYIAVLLTGRPNSRSVGPSGEILAALEAFKLAGISSPFELRPKEGLALVNGTAVGSAMASIVLYEANVLAVFSEVASAMFAEVMHGKPEFSDHLVHKLKHHPGQMEAAAITEHILDGSSFVKEAIRLHEIDPLQKPKQDRYALRTSPQWLGPQIEVIRAATKMIEREINSVNDNPLIDVSRNKAIHGGNFQGTPIGVAMDNARLAIGSIGKLMFAQFTELVNDFYNNGLPSNLSGGRNPSLDYGFKGAEVPMAAYYSELQFLANPVTNHVQSTEQHNQDVNSLGLISSHKTAEAVNILKLMSATYLVALCQAYDLRHLEDNLKETIKAVVNQTAEKHAFTLSKPFIEQNILGVIDREYVFSYVYDLSSLANPLMQKLRSVLFDHALAEPEHETDSGFRKIGTFETELKSLLPNEVERVWTEYENGNFVVANRIKECRSYPLSRFVREELETRLLTGGSVRTPGEDFDEVFKAISKGKLIDPLFECLKEWNGAPIPIS*

>BnaPAL3-1_C04p034430.1_BnaEXP

ATGCTCATCCGTGTCAACACACTCCTCCAAGGCTACTCCGGTATACGCTTCGAAATCCTTGAAGCCATCACAAAGTTCCTCAACCACAAAATCACACCGCGCCTCCCTCTCCGAGGCACCATCACCGCCTCTGGTGACCTCGTTCCTCTATCGTACATCGCCGTTCTCCTCACCGGACGGCCCAACTCCCGATCCGTGGGTCCATCCGGTGAGATCCTTGCCGCCTTAGAGGCCTTCAAGCTAGCTGGAATATCTTCTCCTTTTGAGCTCCGGCCTAAAGAAGGGCTTGCGCTCGTGAACGGCACCGCGGTTGGGTCTGCTATGGCCTCAATAGTGCTATACGAGGCCAACGTTTTGGCAGTGTTCTCGGAAGTTGCTTCTGCCATGTTTGCCGAGGTTATGCATGGGAAACCCGAGTTTTCCGATCATCTTGTGCATAAACTCAAGCACCATCCTGGTCAGATGGAAGCCGCGGCAATCACGGAACATATCTTAGACGGAAGCTCTTTTGTCAAAGAAGCTATACGTCTCCACGAGATTGATCCGCTCCAAAAACCTAAACAAGATCGCTACGCTCTGCGAACATCGCCACAGTGGCTTGGACCTCAGATTGAGGTGATCCGAGCCGCGACGAAAATGATCGAACGAGAGATAAACTCTGTCAACGATAACCCCTTAATCGATGTGTCGCGAAATAAAGCTATCCACGGTGGGAATTTCCAAGGGACACCAATAGGTGTCGCCATGGACAACGCTCGTCTAGCCATTGGTTCCATCGGGAAACTAATGTTTGCGCAGTTCACGGAACTCGTCAATGATTTCTACAACAACGGTCTACCTTCGAATCTATCCGGTGGGAGAAACCCTAGCCTTGACTACGGGTTTAAAGGCGCGGAAGTCCCCATGGCTGCTTATTATTCCGAGCTTCAGTTCTTGGCTAACCCTGTGACAAACCATGTCCAAAGCACTGAACAACATAATCAAGATGTTAACTCTCTTGGTTTGATCTCCAGCCACAAAACTGCCGAAGCTGTGAATATCCTCAAGCTTATGTCAGCTACTTACTTAGTAGCTTTATGCCAAGCCTATGATCTAAGACATCTTGAAGACAATCTCAAGGAAACGATTAAAGCGGTTGTGAACCAAACCGCGGAAAAACACGCATTCACTCTCAGCAAACCGTTCATCGAACAGAACATCCTCGGAGTTATCGACCGCGAATATGTCTTTTCCTATGTTTATGACCTAAGTAGCCTCGCTAACCCTCTAATGCAGAAGCTGAGAAGCGTTCTTTTCGACCATGCTTTAGCTGAACCGGAGCATGAGACGGATTCGGGTTTTCGAAAAATAGGAACGTTCGAAACAGAGCTGAAATCTCTTCTACCTAACGAAGTTGAAAGAGTGTGGACCGAGTACGAAAATGGTAACTTTGTTGTGGCCAACCGAATCAAAGAGTGTAGATCGTATCCGTTGTCCCGGTTTGTACGGGAGGAGCTAGAGACGAGGCTACTAACCGGAGGGAGTGTCCGGACACCAGGAGAAGATTTTGACGAAGTTTTTAAAGCCATCTCTAAGGGAAAACTCATAGATCCTCTGTTTGAATGTCTCAAAGAATGGAACGGAGCTCCAATTCCAATCTCCTAA

**Supplementary Table 7: Organ-specific expression of *B. napus* Express 617 *PAL* candidate genes.** The median transcripts per millions (TPMs) of each *PAL* candidate gene per organ is listed. Paired-end RNA-Seq data generated in this study derived from seeds (35 DAF) of Express 617 (E617) and SGDH14 (S14) are marked with an asterisk. Publicly available paired-end *B. napus* RNA-Seq data sets were used for the remaining organs. The number of analysed data sets per organ is stated via (n = X). The colour gradient from white to blue indicates the expression strength with dark blue symbolising high expression. DAF, days after flowering; DAP, days after pollination; SAM, shoot apical meristem.

|  | *PAL4-1  C05p041260.1* | *PAL4-2 A05p031299.1* | *PAL3-1  C04p034430.1* | *PAL3-2 A04p012590.1* | *PAL2-1  C04p025700.1* | *PAL2-2  A04p005140.1* | *PAL2-3  R186_chrC06p000250.1* | *PAL2-4  A07p016620.1* | *PAL2-5  C08p028640.1* | *PAL2-6  A09p036110.1* | *PAL1-1 C04p009110.1* | *PAL1-2  A05p007980.1* | *PAL1-3  C04p046000.1* | *PAL1-4  A04p023010.1* |
| --- | --- | --- | --- | --- | --- | --- | --- | --- | --- | --- | --- | --- | --- | --- |
| SAM (n=16) | 1.3 | 8.4 | 0.0 | 0.0 | 2.3 | 0.6 | 0.0 | 0.0 | 0.0 | 2.2 | 4.2 | 1.5 | 8.7 | 0.1 |
| Anther prophase 1 (n=12) | 0.1 | 0.8 | 0.0 | 0.0 | 3.1 | 2.4 | 0.0 | 0.0 | 0.0 | 3.7 | 0.1 | 0.0 | 0.1 | 0.2 |
| Anther bolting (n=6) | 0.7 | 54.0 | 0.0 | 0.1 | 3.0 | 1.8 | 0.0 | 0.0 | 0.0 | 4.5 | 27.9 | 14.8 | 19.9 | 0.0 |
| Anther flowering (n=4) | 0.0 | 70.0 | 0.0 | 0.0 | 9.0 | 5.8 | 0.0 | 0.0 | 0.0 | 13.1 | 33.1 | 31.6 | 25.8 | 20.5 |
| Stamen (n=1) | 0.1 | 30.5 | 0.0 | 0.0 | 0.4 | 1.0 | 0.0 | 0.0 | 0.0 | 1.1 | 9.3 | 1.5 | 13.4 | 4.1 |
| Ovule (n=1) | 0.8 | 135.2 | 0.0 | 0.0 | 18.3 | 20.6 | 0.0 | 0.0 | 0.0 | 19.5 | 25.4 | 9.1 | 72.7 | 17.5 |
| Pistil (n=3) | 7.1 | 11.9 | 0.0 | 0.0 | 4.6 | 2.8 | 0.0 | 0.0 | 0.0 | 6.7 | 12.2 | 7.0 | 26.4 | 3.7 |
| Sepal (n=1) | 0.0 | 2.0 | 0.0 | 0.0 | 7.8 | 5.9 | 3.1 | 0.0 | 0.0 | 19.6 | 6.9 | 0.0 | 5.9 | 0.0 |
| Petal (n=2) | 217.4 | 29.2 | 0.0 | 0.0 | 4.2 | 1.2 | 0.8 | 0.6 | 0.0 | 3.7 | 37.3 | 232.9 | 7.4 | 0.0 |
| Silique 10-20DAF (n=13) | 9.4 | 22.3 | 0.0 | 0.0 | 10.3 | 6.0 | 0.0 | 0.0 | 0.0 | 12.8 | 29.1 | 12.1 | 13.7 | 7.7 |
| Silique 25DAF (n=6) | 6.4 | 20.2 | 0.0 | 0.0 | 11.1 | 11.3 | 0.0 | 0.0 | 0.0 | 18.1 | 42.6 | 3.3 | 12.5 | 23.8 |
| Silique 30DAF (n=6) | 10.7 | 33.8 | 0.0 | 0.0 | 13.7 | 13.1 | 0.0 | 0.0 | 0.0 | 18.7 | 44.7 | 6.1 | 11.0 | 17.6 |
| Silique 40DAF (n=2) | 6.5 | 22.2 | 0.0 | 0.0 | 7.3 | 8.1 | 0.0 | 0.0 | 0.0 | 10.2 | 19.5 | 11.9 | 2.9 | 6.4 |
| Seed_35DAF_E617* (n=3) | 54.9 | 87.7 | 0.0 | 0.0 | 9.6 | 11.7 | 0.0 | 0.0 | 0.0 | 18.1 | 19.4 | 31.2 | 6.9 | 6.6 |
| Seed_35DAF_S14* (n=3) | 0.7 | 217.0 | 0.0 | 0.0 | 18.6 | 15.2 | 0.0 | 0.0 | 0.0 | 18.2 | 34.1 | 54.2 | 6.5 | 12.6 |
| Seed coat 14DAF (n=7) | 0.9 | 2.5 | 0.0 | 0.0 | 7.7 | 4.9 | 0.0 | 0.0 | 0.0 | 5.4 | 1.5 | 0.0 | 3.7 | 1.5 |
| Seed coat 21DAF (n=6) | 38.6 | 69.5 | 0.0 | 0.0 | 19.5 | 14.1 | 0.0 | 0.0 | 0.0 | 16.8 | 24.7 | 0.0 | 34.1 | 12.2 |
| Seed coat 28DAF (n=6) | 120.7 | 151.6 | 0.0 | 0.1 | 19.2 | 14.0 | 0.0 | 0.0 | 0.0 | 20.9 | 42.5 | 0.0 | 95.5 | 42.2 |
| Seed coat 35DAF (n=6) | 92.4 | 90.8 | 0.0 | 0.1 | 18.3 | 16.2 | 0.0 | 0.0 | 0.0 | 18.0 | 48.4 | 0.0 | 91.0 | 42.0 |
| Seed coat 42DAF (n=6) | 111.9 | 82.6 | 0.0 | 0.0 | 14.6 | 13.2 | 0.0 | 0.0 | 0.0 | 21.4 | 245.5 | 0.0 | 63.1 | 27.8 |
| Embryo (n=6) | 13.5 | 20.4 | 0.0 | 0.0 | 10.5 | 9.8 | 0.0 | 0.0 | 0.0 | 25.4 | 9.5 | 3.2 | 2.1 | 10.9 |
| Endosperm (n=8) | 15.2 | 33.4 | 0.0 | 0.0 | 4.7 | 4.1 | 0.0 | 0.0 | 0.0 | 5.5 | 3.1 | 3.0 | 10.4 | 1.3 |
| Seedling (n=9) | 0.2 | 7.5 | 0.0 | 0.1 | 13.2 | 10.5 | 0.0 | 0.0 | 0.0 | 23.6 | 15.5 | 10.9 | 18.6 | 0.0 |
| Stem (n=19) | 126.9 | 227.9 | 0.0 | 0.0 | 29.0 | 25.1 | 0.0 | 0.0 | 0.0 | 67.9 | 131.6 | 59.7 | 117.5 | 19.9 |
| Shoot (n=2) | 2.1 | 8.2 | 0.0 | 0.0 | 1.8 | 0.9 | 0.0 | 0.0 | 0.0 | 3.9 | 4.9 | 2.3 | 5.4 | 6.0 |
| Shoot apex (n=2) | 11.7 | 8.1 | 0.0 | 0.0 | 4.6 | 1.6 | 0.0 | 0.0 | 0.0 | 6.4 | 4.3 | 1.1 | 3.4 | 1.3 |
| Root seedling (n=13) | 8.4 | 7.6 | 0.0 | 0.5 | 18.4 | 14.1 | 0.0 | 0.0 | 0.0 | 56.7 | 44.3 | 25.8 | 31.2 | 3.6 |
| Root 30DAP (n=20) | 6.5 | 7.4 | 0.0 | 0.6 | 66.3 | 55.6 | 0.0 | 0.0 | 0.0 | 82.5 | 92.6 | 21.6 | 77.8 | 54.7 |
| Root 60DAP (n=2) | 0.5 | 13.8 | 0.0 | 0.2 | 132.0 | 148.0 | 8.0 | 0.0 | 0.0 | 252.5 | 290.0 | 17.5 | 364.5 | 1.0 |

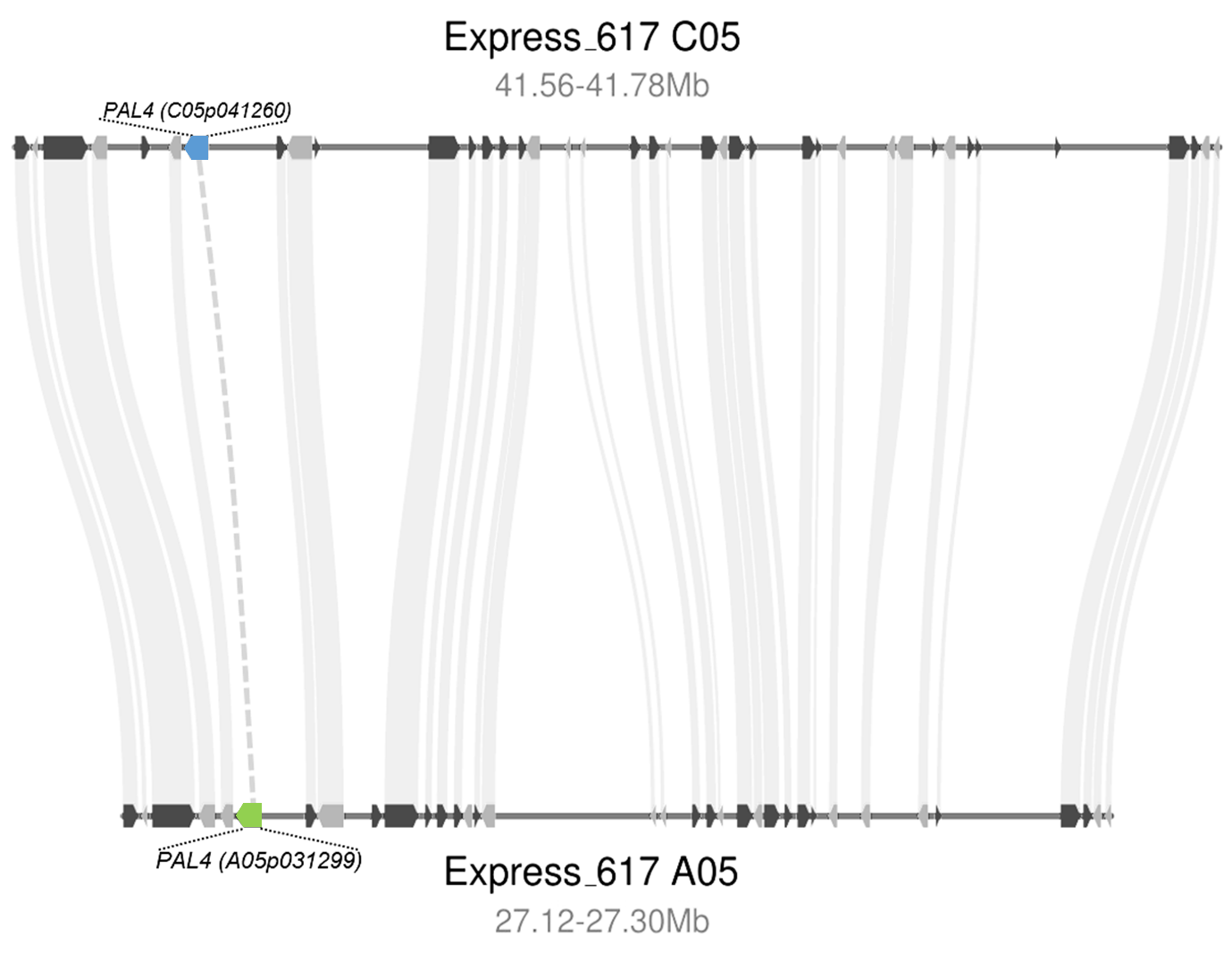

**Supplementary Figure 5: *B. napus* Express 617 carries a previously not annotated A05 *PAL4* homolog and shows high local synteny between the homologous A05 region to the C05 region.** The synteny plot was constructed based on the Express 617 assembly and covers the C05 region (41,56-41,78 million base pairs (Mb)) located inside the major low lignin QTL, as well as the corresponding homologous A05 region ranging from 27,12-27,30 million base pairs (Mb). Genes located on the forward strand are marked in black, while genes located on the reverse strand are marked in grey. The *PAL4* copy on C05 is highlighted in blue, while the manually annotated *PAL4* copy on A05 is marked in green. Respective novel syntenic genes are marked by a dashed line.

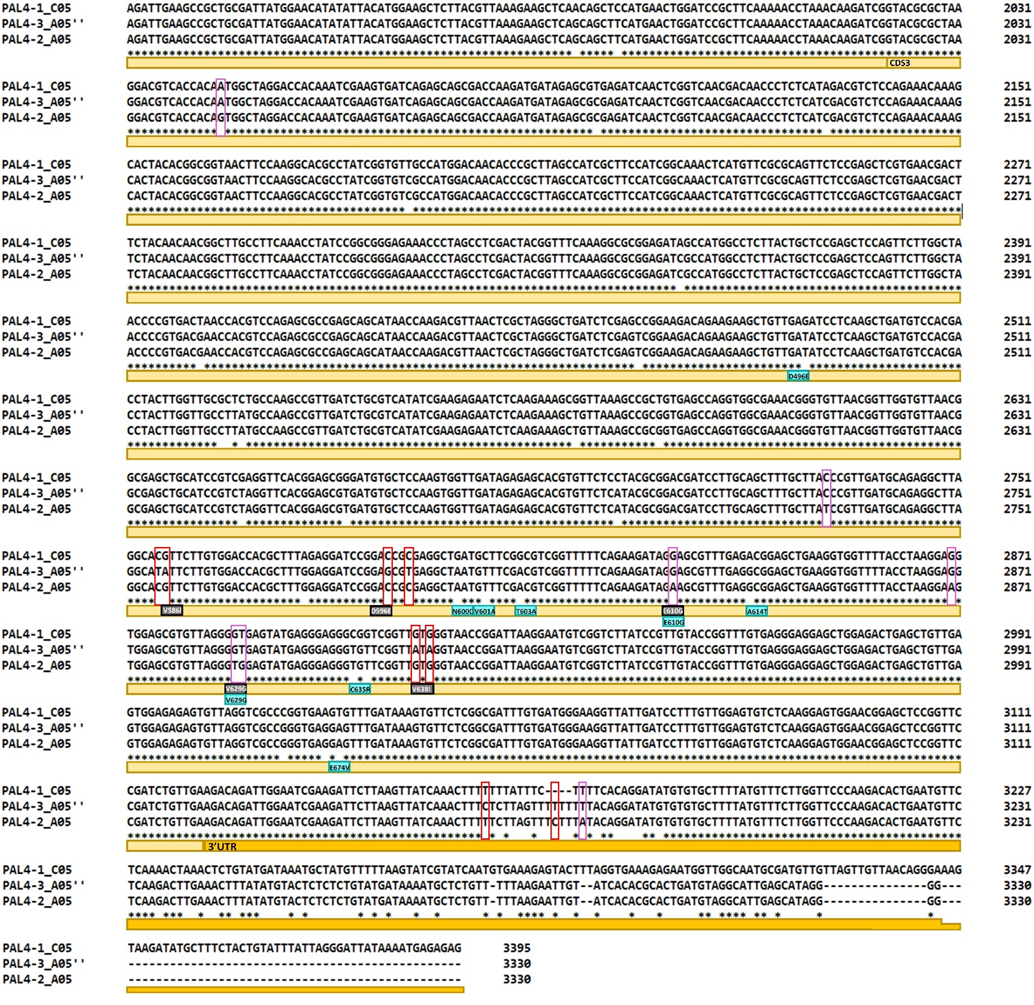

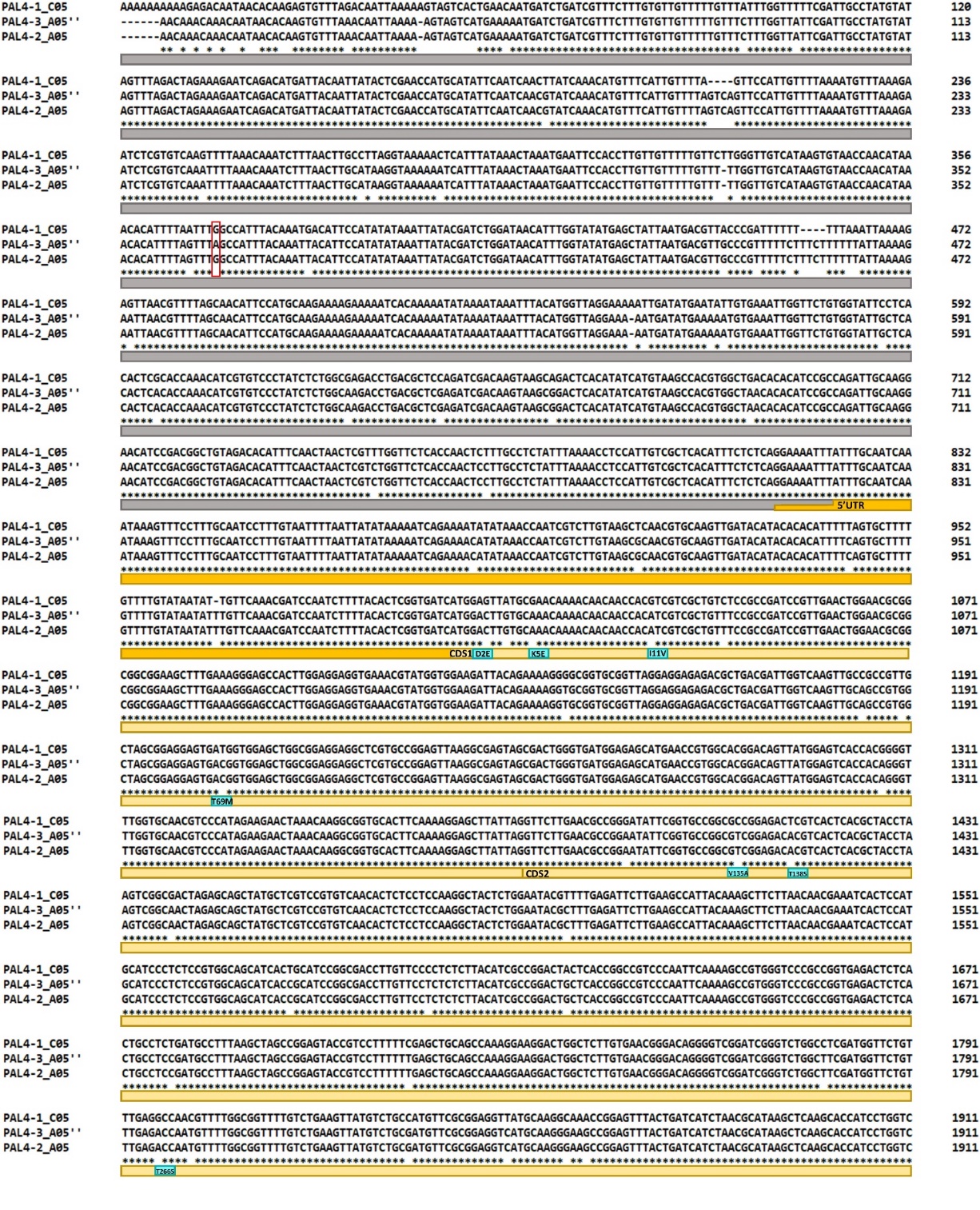

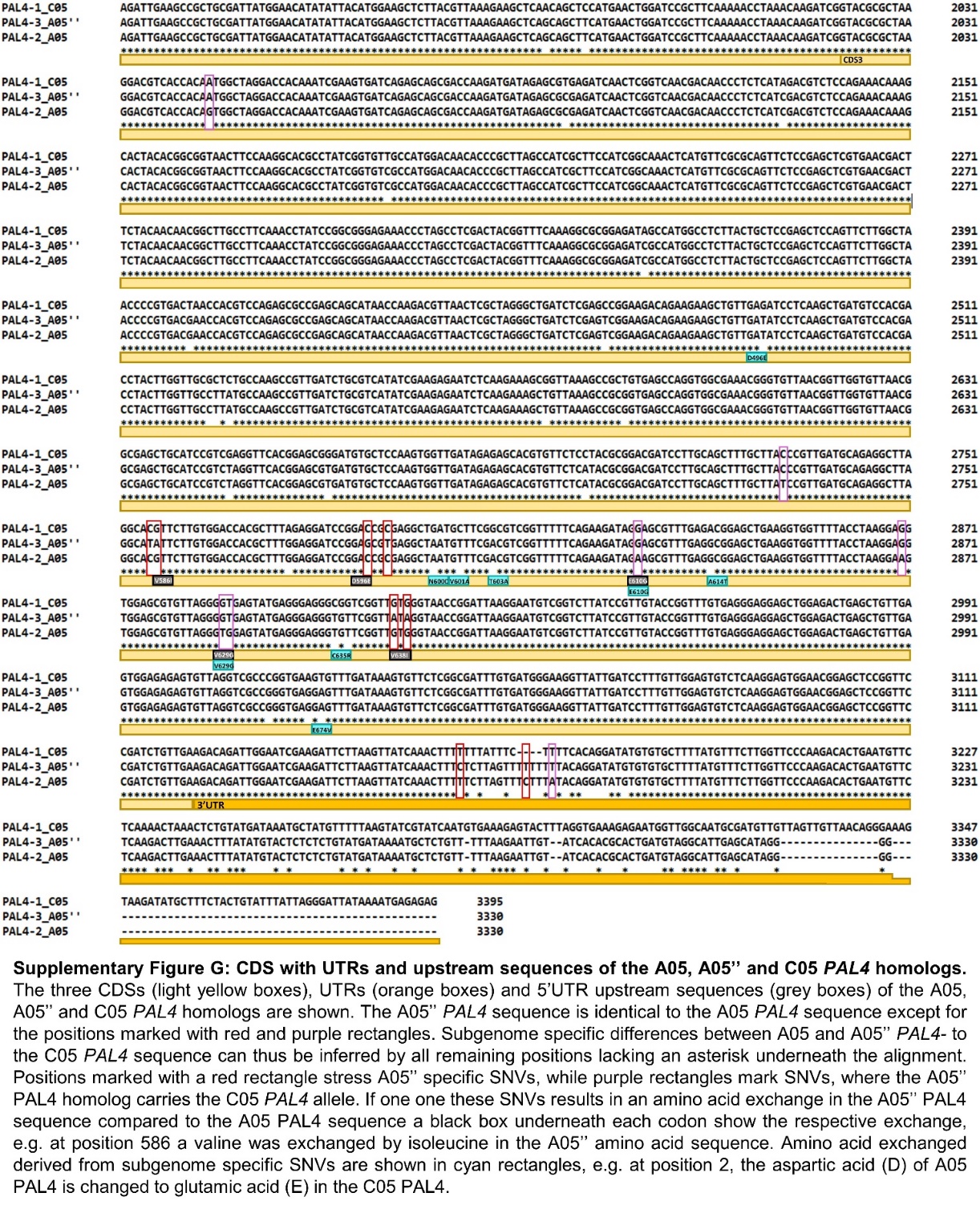

**Supplementary Figure 6: The CDS with UTRs and upstream sequences of the A05, A05’’, and C05 *PAL4* homologs.** The CDS (light yellow box), UTRs (orange boxes) and 5’UTR upstream sequences (grey boxes) of the A05, A05’’ and C05 *PAL4* homologs are shown. The A05’’ *PAL4* sequence is identical to the A05 *PAL4* sequence except for the positions marked with red and purple rectangles. Subgenome-specific differences between A05 and A05’’ *PAL4-* to the C05 *PAL4* sequence can thus be inferred by all remaining positions lacking an asterisk underneath the alignment. Positions marked with a red rectangle stress A05’’ specific SNPs, while purple rectangles mark SNPs, where the A05’’ PAL4 homolog carries the C05 *PAL4* allele. If one one these SNPs results in an amino acid exchange in the A05’’ PAL4 sequence compared to the A05 PAL4 sequence a black box underneath each codon show the respective exchange, e.g. at position 586 a valine was exchanged by isoleucine in the A05’’ amino acid sequence. Amino acid exchanged derived from subgenomes-specific SNPs are shown in cyan rectangles, e.g. at position 2, the aspartic acid (D) of A05 PAL4 is changed to glutamic acid (E) in the C05 PAL4. Clustal Omega (Sievers et al. 2011) was applied for the construction of the sequence alignment.
