## Supplementary Information 8 for "Homoeologous non-reciprocal translocation explains a major QTL for seed lignin content in oilseed rape (*Brassica napus* L.)"

P1

P2

R1

```
Express617_C05_RB      GCTTAATCTGAAACACGGGCTTGGGTATCAAGTTTACCCTAACGGAGACGTCTTAGAAGG     60
Express617_A05_RB      GCTGAATCTCAAACACGGGCTTGGGTACCAAGTTTACCCTAACGGTGACGTCTTCGAAGG     60
SGDH14_C05_RB          GCTGAATCTCAAACACGGGCTTGGGTACCAAGTTTACCCTAACGGTGACGTCTTCGAAGG     60
SGDH14_A05_RB          GCTGAATCTCAAACACGGGCTTGGGTACCAAGTTTACCCTAACGGTGACGTCTTCGAAGG     60
                       *** ***** ***************** ***************** ******** *****

Express617_C05_RB      CTCTTGGATTCAGGGTTGGGGAGAAGGGCCAGGGAAGTACACTTGGGGTAACGGGAACAT     120
Express617_A05_RB      CTCTTGGATTCAGGGTTTGGGAGAAGGGCCAGGGAAGTACACGTGGGGGAACGGGAACAT     120
SGDH14_C05_RB          CTCTTGGATTCAGGGTTGGGGAGAAGGGCCAGGGAAGTACACTTGGGGTAACGGGAACAT     120
SGDH14_A05_RB          CTCTTGGATTCAGGGTTTGGGAGAAGGGCCAGGGAAGTACACGTGGGGGAACGGGAACAT     120
                       ***************** ************************ ***** ***********

Express617_C05_RB      CTATCTTGGGGATATGAAAGGTGGGAA      147
Express617_A05_RB      CTATCTTGGGGATATGAAAGGTGGGAA      147
SGDH14_C05_RB          CTATCTTGGGGATATGAAAGGTGGGAA      147
SGDH14_A05_RB          CTATCTTGGGGATATGAAAGGTGGGAA      147
                       ***************************
```

<G> 3‘ Anchor R1 reverse primer

[A/C 3‘ SNP P1 forward primer; alternative [A/T] 3‘ SNP P2 forward primer

GAAG-GGTT 22 bp non subgenome-specific region

**Supplementary Figure 7: Anchored KASP assay design for homoeologous non-reciprocal translocation in SGDH14.** An anchored KASP INDEL marker was designed to detect the A05/C05 insertion using the marked mismatches in both sequences on the right border.

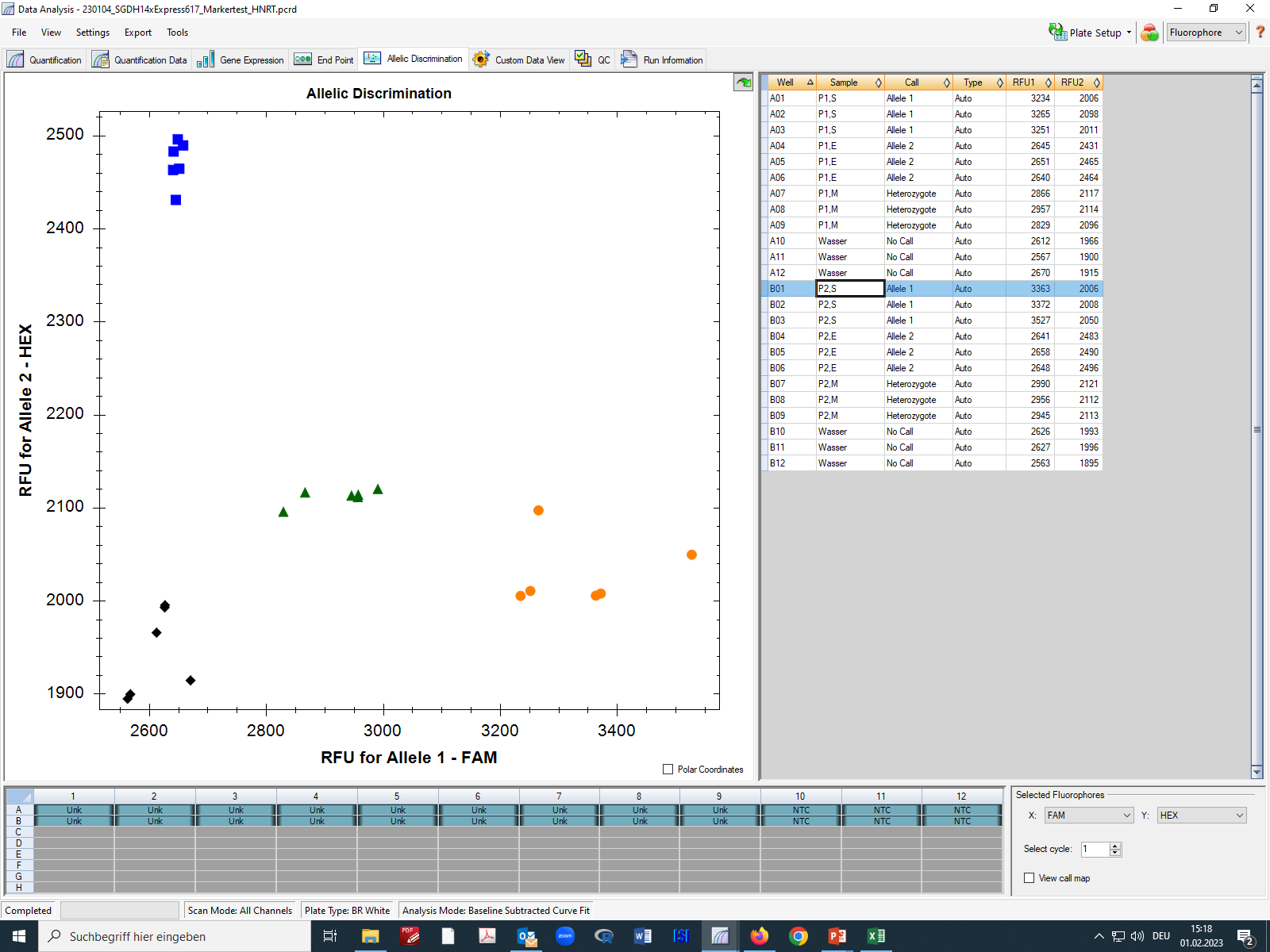

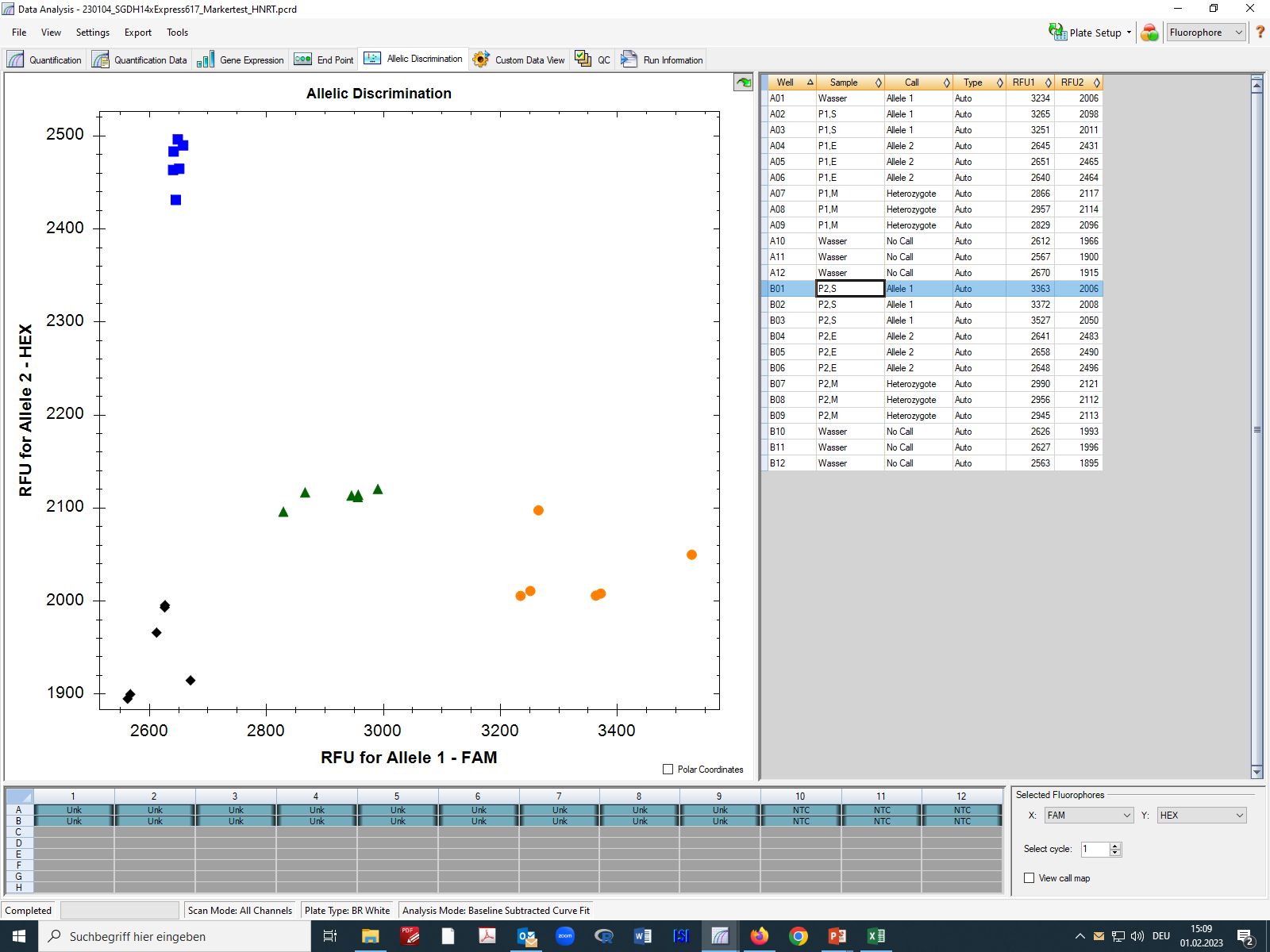

P1; E = Express; n=3

P2; E = Express; n=3

P1; S = SGDH14; n=3

P2; S = SGDH14; n=3

P1; M = Mix of DNA; n=3

P2; M = Mix of DNA; n=3

**Supplementary Figure 8: Anchored KASP assay results for homoeologous non-reciprocal translocation in SGDH14.** KASP anchored SNP result of Express 617 (blue rectangles), SGDH14 (orange dots), and a mixed samples of both genotypes (green triangles) were used for KASP assay. The black diamonds mark the water controls. Two forward primers were tested (P1 and P2), which performed equally well. RFU, relative fluorescence units.
